# A framework for detecting noncoding rare variant associations of large-scale whole-genome sequencing studies

**DOI:** 10.1101/2021.11.05.467531

**Authors:** Zilin Li, Xihao Li, Hufeng Zhou, Sheila M. Gaynor, Margaret S. Selvaraj, Theodore Arapoglou, Corbin Quick, Yaowu Liu, Han Chen, Ryan Sun, Rounak Dey, Donna K. Arnett, Lawrence F. Bielak, Joshua C. Bis, Thomas W. Blackwell, John Blangero, Eric Boerwinkle, Donald W. Bowden, Jennifer A. Brody, Brian E. Cade, Matthew P. Conomos, Adolfo Correa, L. Adrienne Cupples, Joanne E. Curran, Paul S. de Vries, Ravindranath Duggirala, Barry I. Freedman, Harald H. H. Göring, Xiuqing Guo, Rita R. Kalyani, Charles Kooperberg, Brian G. Kral, Leslie A. Lange, Ani Manichaikul, Lisa W. Martin, Braxton D. Mitchell, May E. Montasser, Alanna C. Morrison, Take Naseri, Jeffrey R. O’Connell, Nicholette D. Palmer, Patricia A. Peyser, Bruce M. Psaty, Laura M. Raffield, Susan Redline, Alexander P. Reiner, Muagututi‘a Sefuiva Reupena, Kenneth M. Rice, Stephen S. Rich, Jennifer A. Smith, Kent D. Taylor, Ramachandran S. Vasan, Daniel E. Weeks, James G. Wilson, Lisa R. Yanek, Wei Zhao, NHLBI Trans-Omics for Precision Medicine (TOPMed) Consortium, TOPMed Lipids Working Group, Jerome I. Rotter, Christen J. Willer, Pradeep Natarajan, Gina M. Peloso, Xihong Lin

## Abstract

Large-scale whole-genome sequencing studies have enabled analysis of noncoding rare variants’ (RVs) associations with complex human traits. Variant set analysis is a powerful approach to study RV association, and a key component of it is constructing RV sets for analysis. However, existing methods have limited ability to define analysis units in the noncoding genome. Furthermore, there is a lack of robust pipelines for comprehensive and scalable noncoding RV association analysis. Here we propose a computationally-efficient noncoding RV association-detection framework that uses STAAR (variant-set test for association using annotation information) to group noncoding variants in gene-centric analysis based on functional categories. We also propose SCANG (scan the genome)-STAAR, which uses dynamic window sizes and incorporates multiple functional annotations, in a non-gene-centric analysis. We furthermore develop STAARpipeline to perform flexible noncoding RV association analysis, including gene-centric analysis as well as fixed-window-based and dynamic-window-based non-gene-centric analysis. We apply STAARpipeline to identify noncoding RV sets associated with four quantitative lipid traits in 21,015 discovery samples from the Trans-Omics for Precision Medicine (TOPMed) program and replicate several noncoding RV associations in an additional 9,123 TOPMed samples.

## Introduction

Genome-wide association studies (GWASs) have successfully identified thousands of common genetic variants for complex diseases and traits; however, these common variants only explain a small fraction of heritability^1^. Recent studies suggest that the missing heritability of complex traits and diseases and causal variants may be accounted for in part by RVs (minor allele frequency (MAF) < 1%)^2-4^. Although whole-exome sequencing (WES) studies have identified exome-wide significant RV associations for complex diseases and traits^5,6^, more than 98% of the genetic variants are located in the noncoding genome^6^. Many common variants identified by GWAS as being associated with phenotypes are located in noncoding regions^7,8^. Further, the ENCODE project shows that a significant fraction of noncoding regions are functionally active^9,10^, indicating that rare noncoding regions may have an effect on diseases or traits.

An increasing number of whole-genome sequencing (WGS) association studies, such as the Genome Sequencing Program (GSP) of the National Human Genome Research Institute (NHGRI) and the Trans-Omics for Precision Medicine (TOPMed) Program of the National Heart, Lung, and Blood Institute (NHLBI), permit the study of the genetic contributions of noncoding RVs to complex traits and diseases. It is of substantial interest to use these rich WGS data to explore the role of noncoding RVs in the genetic underpinning of common human diseases.

Single-variant analyses are not appropriate for analysis of rare variants because in realistic settings they lack power^11-13^. To improve power, variant set tests have been proposed that assess the effects of sets of multiple RVs jointly. These tests include burden, SKAT, and most recently STAAR (variant-set test for association using annotation information), which incorporates multiple functional annotations for genetic variants^14-16^. A key challenge of these approaches is the selection of RVs to form variant sets. Several methods have been proposed to create coding and noncoding variant sets for RV association analysis of WGS/WES studies^16-20^. However, these methods have limited utility for defining analysis units in the noncoding genome^21^. For example, for gene-centric analysis, STAAR has been used with two noncoding genetic categories of regulatory regions (masks): using promoters and enhancers in GeneHancer^22^ overlaid with Cap Analysis of Gene Expression (CAGE) sites^23,24^; for non-gene-centric analysis, fixed-size sliding windows can be used to scan the genome. As the signal regions (variant-phenotype-association regions) are unknown in practice and their sizes vary across the genome, the fixed-size sliding window approach is likely to lead to power loss when the prespecified window sizes are too big or too small compared with the actual sizes of signal regions. Furthermore, it is often knowledge- and effort-demanding to functionally annotate variants from a WGS/WES study of interest. Limited tools exist for multi-faceted functional annotation and analytic integration of WGS/WES data for rare variant association tests (RVATs). Finally, there is a lack of robust pipelines to perform scalable and comprehensive noncoding RV association analysis in large-scale WGS data with hundreds of millions of noncoding RVs that have been sequenced across the genome. Much uncertainty remains on the best practices for computationally-efficient RV analysis at the scale of large WGS studies.

To respond to the aforementioned needs, we propose a computationally-efficient noncoding rare variant association-detection framework for WGS data by making three new contributions toward automatically selecting interpretable and powerful variant sets. First, in gene-centric analysis, we propose additional strategies for grouping noncoding variants based on functional annotations, including untranslated regions, upstream regions, downstream regions, promoters, enhancers of protein-coding genes, and long noncoding RNA genes within STAAR. For promoters and enhancers, we offer additional options of overlaying promoters and GeneHancer-based enhancers with not only CAGE sites but also with DNase Hypersensitivity (DHS) sites^9^. Second, in non-gene-centric analysis, instead of using fixed-size sliding windows in STAAR we propose SCANG-STAAR, a flexible data-adaptive window size RVAT method that extends the SCANG (scan the genome) method^18^ by incorporating multiple functional annotations through STAAR^16^, while accounting for both relatedness and population structure through a generalized linear mixed model framework^25^ for quantitative and dichotomous traits^26,27^. Third, we develop *STAARpipeline*, a pipeline that (1) functionally annotates both noncoding and coding variants of a WGS study and builds an annotated genotype dataset using the multi-faceted functional annotation database FAVOR^16^ (Functional Annotations of Variants - Online Resource), through FAVORannotator; and (2) performs RVATs using the proposed methods for both gene-centric analysis and non-gene-centric analysis.

We applied the proposed framework to detect noncoding RVs associated with four quantitative lipid traits: low-density lipoprotein cholesterol (LDL-C); high-density lipoprotein cholesterol (HDL-C); triglycerides (TG) and total cholesterol (TC) using 21,015 discovery samples and 9,123 replication samples from the NHLBI TOPMed Freeze 5 WGS data. We performed conditional analysis by conditioning on known lipids-associated variants and identified several novel replicated RVs sets associated with lipids.

## Results

### Overview of Noncoding RVATs

We propose a computationally-efficient noncoding RVAT framework for phenotype-genotype association analyses of whole-genome sequencing data, focusing on rare variant association analysis in the noncoding genome. This regression-based framework allows adjusting for covariates, population structure, and relatedness by fitting linear and logistic mixed models for quantitative and dichotomous traits^26,27^. A central component of it is the development of strategies to aggregate noncoding rare variants using both flexible gene-centric and non-gene-centric approaches to empower RVATs. For the gene-centric approach, we group noncoding RVs for each gene using eight genetic categories of regulatory regions provided by functional annotations and apply STAAR, which incorporates multiple *in-silico* variant functional annotation scores that prioritize functional variants using multi-dimensional variant biological functions^16^. For the non-gene-centric analysis, instead of using sliding windows with fixed sizes, we propose SCANG-STAAR, a procedure using dynamic windows with data-adaptive sizes and incorporating multi-dimensional functional annotations. We also perform analytical follow-up to dissect RV association signals independent of a given set of known variants via conditional analysis (**Figure 1**).

**Figure 1.**
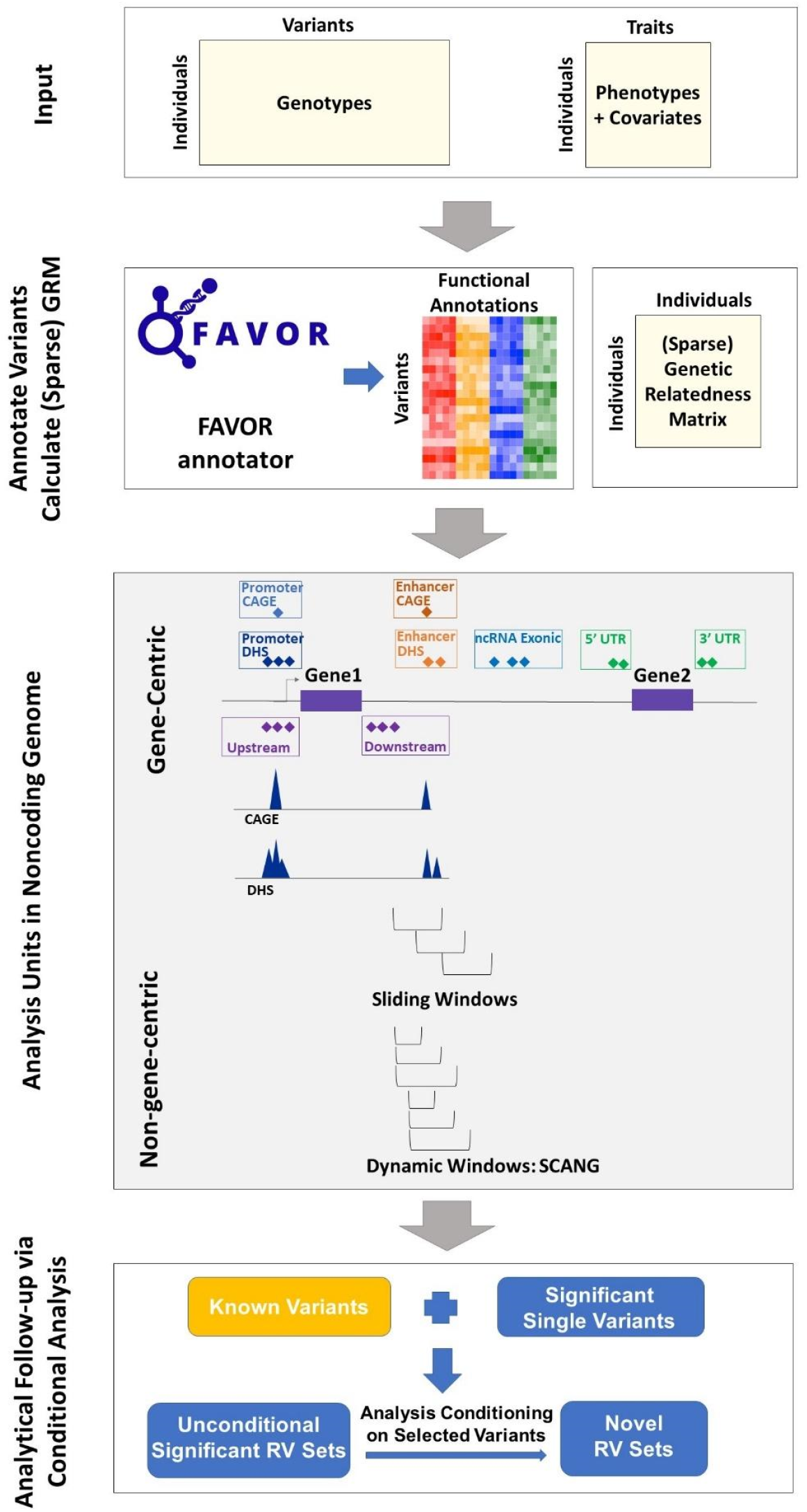
Workflow of *STAARpipeline*. (a) Prepare the input data of *STAARpipeline*, including genotypes, phenotypes and covariates. (b) Annotate all variants in the genome using FAVORannotator through FAVOR database and calculate the (sparse) genetic relatedness matrix. (c) Define analysis units in the noncoding genome: eight genetic categories of regulatory regions, sliding windows and dynamic windows using SCANG. (d) Obtain genome-wide significant associations and perform analytical follow-up via conditional analysis.

### Gene-centric analysis of the noncoding genome

In gene-centric analysis of noncoding variants, we provide eight genetic categories of regulatory regions to aggregate noncoding rare variants: (1) promoter RVs overlaid with CAGE sites, (2) promoter RVs overlaid with DHS sites, (3) enhancer RVs overlaid with CAGE sites, (4) enhancer RVs overlaid with DHS sites, (5) untranslated region (UTR) RVs, (6) upstream region RVs, (7) downstream region RVs and (8) noncoding RNA (ncRNA) RVs. The promoter RVs are defined as RVs in the +/- 3-kilobase (kb) window of transcription start sites with the overlap of CAGE sites or DHS sites. The enhancer RVs are defined as RVs in GeneHancer predicted regions with the overlap of CAGE sites or DHS sites^9,22-24^. We define the UTR, upstream, downstream, and ncRNA RVs by GENCODE VEP categories^28,29^. For the UTR mask, we include RVs in both 5’ and 3’ UTR regions. For the ncRNA mask, we include the exonic and splicing ncRNA RVs. We consider the protein-coding gene for the first seven categories provided by Ensembl^30^ and the ncRNA genes provided by GENCODE^28,29^.

For each noncoding mask, we calculate its *P* value using the STAAR method that empowers RVATs by incorporating multiple variant functional annotation scores^16^. Functional annotations consist of diverse biological information of genomic elements. Incorporating this external biological information provided by functional annotations can increase the association analysis power^31^. For example, annotation principal components (aPCs) provide multi-dimensional summaries of variant annotations and capture the multi-faceted biological impact, calculated by the first principal component of the set of individual functional annotation scores interpreting similar biological functionality^16^. We incorporate ten aPCs and three integrative scores (CADD^32^, LINSIGHT^33^, and FATHMM-XF^34^) as weights in constructing STAAR statistics^16^. Details of these 13 functional annotations are given in **Supplementary Table 1**. Specifically, we calculate the *P* value of each variant set using STAAR-O^16^, an omnibus test aggregating multiple annotation-weighted burden test^14^, SKAT^15^, and ACAT-V^35^ in the STAAR framework.

### Non-gene-centric analysis using dynamic windows with SCANG-STAA

We improve the STAAR-based fixed-size sliding window RVAT^16,17^ by proposing a dynamic window based SCANG-STAAR method, which extends the procedure SCANG^18^ by incorporating multi-dimensional functional annotations to flexibly detect the locations and the sizes of signal windows across the genome. Specifically, as location of regions associated with a disease or trait are often unknown in advance and their sizes may vary across the genome, RVAT’s default use of a pre-specified fixed-size sliding window method can lead to power loss, if the pre-specified window sizes do not align with the true signal window sizes.

The dynamic window RVAT method, SCANG^18^, overcomes the limitation of the fixed-size sliding window method using scan statistics that flexibly detect the sizes and the locations of RV association by scanning the whole genome continuously while allowing for overlapping windows of different sizes by shifting forward a given size window by a small number of variants each time and selecting the windows that maximize the test power, while controlling for the genome-wise (family-wise) error rate by accounting for the correlations of tests from overlapping windows. However, SCANG does not incorporate variant functional annotations and may therefore lose power if annotation information helps identify true signals. We propose SCANG-STAAR by extending SCANG to incorporate multi-dimensional variant functional annotations using STAAR to ameliorate power loss.

In dynamic window analysis, we extend the SCANG-SKAT procedure (SCANG-S) to SCANG-STAAR-S by using the STAAR-SKAT (STAAR-S) *P* value, which in each overlapping window incorporates multiple variant functional annotations, instead of using just the MAF-based SKAT *P* value. In SCANG-STAAR-S we first calculate a threshold that controls the genome-wise type I error at a given *α* level, based on the minimum value of the STAAR-S *P* value from all moving windows of different sizes in a range of windows (**Online Methods**). The procedure then selects the candidate significant windows whose set-based *P* value beats that threshold. When this results in multiple overlapping windows, we localize the detected significant window as the window whose *P* value is smaller than both the threshold and any window that overlaps with it. We then calculate the genome-wide *P* value of the detected windows by accounting for multiple comparisons of overlapping windows and controlling for the genome-wise (family-wise) error rate (**Online Methods)**.

Besides the SCANG-STAAR-S method, we also provide the SCANG-STAAR-B procedure, based on the STAAR-Burden *P* value. Compared with SCANG-STAAR-B, SCANG-STAAR-S has two advantages in detecting noncoding associations using dynamic windows in practice. First, the effects of causal variants in the noncoding genome tend to be in different directions, especially in the intergenic region. Second, due to the different correlation structures of the two test statistics for overlapping windows, the genome-wide significance threshold of SCANG-STAAR-B is lower than that of SCANG-STAAR-S. For example, to control the genome-wise error rate at 0.05 level in our analysis of LDL-C, the *P* value threshold of SCANG-STAAR-S and SCANG-STAAR-B are 3.80 × 10^−9^ and 2.31 × 10^−10^, respectively. We additionally provide the SCANG-STAAR-O procedure, which is based on an omnibus *P* value of SCANG-STAAR-S and SCANG-STAAR-B calculated by ACAT method^36^. However, different from STAAR-O, we do not incorporate the ACAT-V test in the omnibus test, since the ACAT-V test is designed for sparse alternatives. Hence, it always detects the region with the smallest size that contains the most significant variant in the dynamic window procedure.

### Analytical follow-up via conditional analysis

We also perform conditional analysis as an analytical follow-up to identify RV association signals independent of known single variant associations. We first select a list of known variants by including the previously identified trait-associated variants, for example, variants indexed in the GWAS Catalog^36^. We then perform stepwise selection to select the subset of independent variants from the known variants list to be used in the conditional analysis. We perform iterative conditional association analysis until the *P* values of all variants in the known variant list are larger than a cut-off (1 × 10^−4^, **Online Methods**). Instead of adjusting for all known trait-associated variants in the entire chromosome, we adjust for variants in an extended region of the specific variant, for example, a +/- 1-megabase (Mb) window beyond the variant of interest. Finally, we perform conditional analysis of each variant set by fitting the regression model adjusting for the selected known variants near the variant set (for example, in a +/- 1-Mb window).

### STAARpipeline and computation cost

Our R package *STAARpipeline* performs scalable phenotype-genotype association analyses of functionally annotated WGS data using the developed RVAT methods. A further package, *STAARpipelineSummary* summarizes the rare variant findings generated by *STAARpipeline*, including results of both unconditional and conditional analysis and visualization of analysis results.

Specifically, to perform RVATs for a given WGS study, we first need to functionally annotate the variants and create variant sets. To achieve this, we use FAVORannotator, a workflow that annotates the variants of a given WGS study using the FAVOR database and generates annotated genotype files for use in *STAARpipeline*. Across the genome, *STAARpipeline* runs gene-centric noncoding and sliding window tests using STAAR and dynamic window analysis using SCANG-STAAR. *STAARpipeline* can also perform RV analysis of coding variants and single variant analysis of common and low-frequency variants (**Discussion**).

All analyses can be computed with modest time and memory resources, even for large-scale WGS/WES datasets such as TOPMed, GSP and UK Biobank. We benchmarked *STAARpipeline*’s WGS association analysis of n=30,138 pooled related TOPMed lipids samples including both discovery and replication data in: 15 hours using 200 2.10 GHz computing cores with 11 Gb memory of gene-centric noncoding analysis; or 11 hours using 200 cores with 11 Gb memory of sliding window analysis; or 20 hours using 800 cores with 15 Gb memory of dynamic window analysis (including SCANG-STAAR-S, SCANG-STAAR-B and SCANG-STAAR-O). *STAARpipelineSummary* summarizes the results from *STAARpipeline* and provides analytical follow-up via conditional analysis. Summarizing the genome-wide TOPMed results took 24 hours using one core with 25 Gb memory.

### Association analysis of lipid traits in the TOPMed WGS data

We applied *STAARpipeline* to identify RV-sets associated with four quantitative lipid traits (LDL-C, HDL-C, TG and TC) using TOPMed WGS data^4,16,20^. DNA samples were sequenced at >30X target coverage^4^. The discovery phase consisted of six study cohorts with 21,015 samples sequenced in TOPMed Freeze 5. The replication phase consisted of eight remaining study cohorts with 9,123 samples in TOPMed Freeze 5 (**Supplementary Note, Supplementary Table 2**). Sample-level and variant-level quality control (QC) were performed^4,20^. Race/ethnicity was defined using a combination of self-reported race/ethnicity and study recruitment information^37^. The discovery cohorts consisted of 5,849 (27.8%) Black or African American, 12,313 (58.6%) White, 675 (3.2%) Asian American, 1,075 (5.1%) Hispanic/Latino American, and 1,103 (5.3%) Samoan participants. Among all samples in the discovery phase, 3,610 (17.2%) had first degree relatedness, 546 (2.6%) had second degree relatedness, and 472 (2.2%) had third degree relatedness (**Supplementary Figure 1**). There were 215 million single-nucleotide variants (SNVs) observed in the discovery phase, and 205 million (94.9%) were rare variants (MAF < 1%). Among these 205 million rare variants, 202 million (98.8%) were noncoding variants defined by GENCODE VEP. Details of the study-specific demographics, summaries of lipid levels, and variant number distributions are given in **Supplementary Tables 2-3** and **Supplementary Figure 2**.

For each phenotype, we applied rank-based inverse normal transformation of phenotypes. We adjusted for age, age^2^, sex, race/ethnicity, study, and the first 10 ancestral PCs, and controlled for relatedness through heteroscedastic linear mixed models with sparse genetic relatedness matrices (GRMs) plus study-race/ethnicity-specific group-specific residual variance components (**Online Methods**). We accounted for the presence of medications of LDL-C and TC as before^20^. We tested for an association between lipid traits and RVs (MAF < 1%) in each variant set. In gene-centric analysis, we defined the eight analysis units as the previously-described: seven noncoding genetic categories of protein-coding genes and one category for ncRNA genes. In non-gene-centric analysis, we performed a 2-kb sliding window analysis with 1-kb skip length and a dynamic window analysis using SCANG-STAAR-S of all moving windows containing 40 to 300 variants^18^. In unconditional analysis we used Bonferroni-corrected genome-wide significance thresholds of *α* = 0.05/(20,000 × 7) = 3.57 × 10^−7^ accounting for 7 different noncoding masks across protein-coding genes; *α* = 0.05/20,000 = 2.50 × 10^−6^ accounting for ncRNA genes, and *α* = 0.05/(2.66 × 10^6^) = 1.88 × 10^−8^ accounting for 2.66 million 2-kb sliding windows across the genome. We controlled the genome-wise (family-wise) error rate for SCANG-STAAR-S dynamic window analysis at *α* = 0.05 level^18^. We selected individual variants to be adjusted for in conditional analysis from the list of phenotype-associated common and low-frequency variants (MAF ≥ 1%) indexed in GWAS Catalog^36^. Then we obtained the independent known variants using the algorithm described before in the analytical follow-up via conditional analysis section (**Online Methods, Supplementary Table 4**).

In gene-centric noncoding unconditional analysis of the discovery samples, *STAARpipeline* identified 43 genome-wide significant associations with at least one of the four lipid levels (**Supplementary Table 5, Supplementary Figures 3a-d, 4a-d, 5a-d, 6a-d**). After conditioning on known lipid-associated variants, 14 out of the 43 associations remained significant at the Bonferroni-corrected level 0.05/43 = 1.16 × 10^−3^ (**Table 1**). In the replication data, and adjusting for known lipid-associated variants, 4 of these 14 associations achieved significance at Bonferroni-corrected level 0.05/14 = 3.57 × 10^−3^. These included enhancer DHS RVs in *APOA1* and HDL-C, promoter CAGE RVs in *APOE* and TG, and enhancer CAGE or DHS RVs in *APOE* and TG. After further adjustment for known individual rare variants (minor allele count, MAC ≥ 20, **Supplementary Table 6**), none of the associations remained significant at the same significance level of 3.57 × 10^−3^ (**Supplementary Table 7**).

**Table 1.**
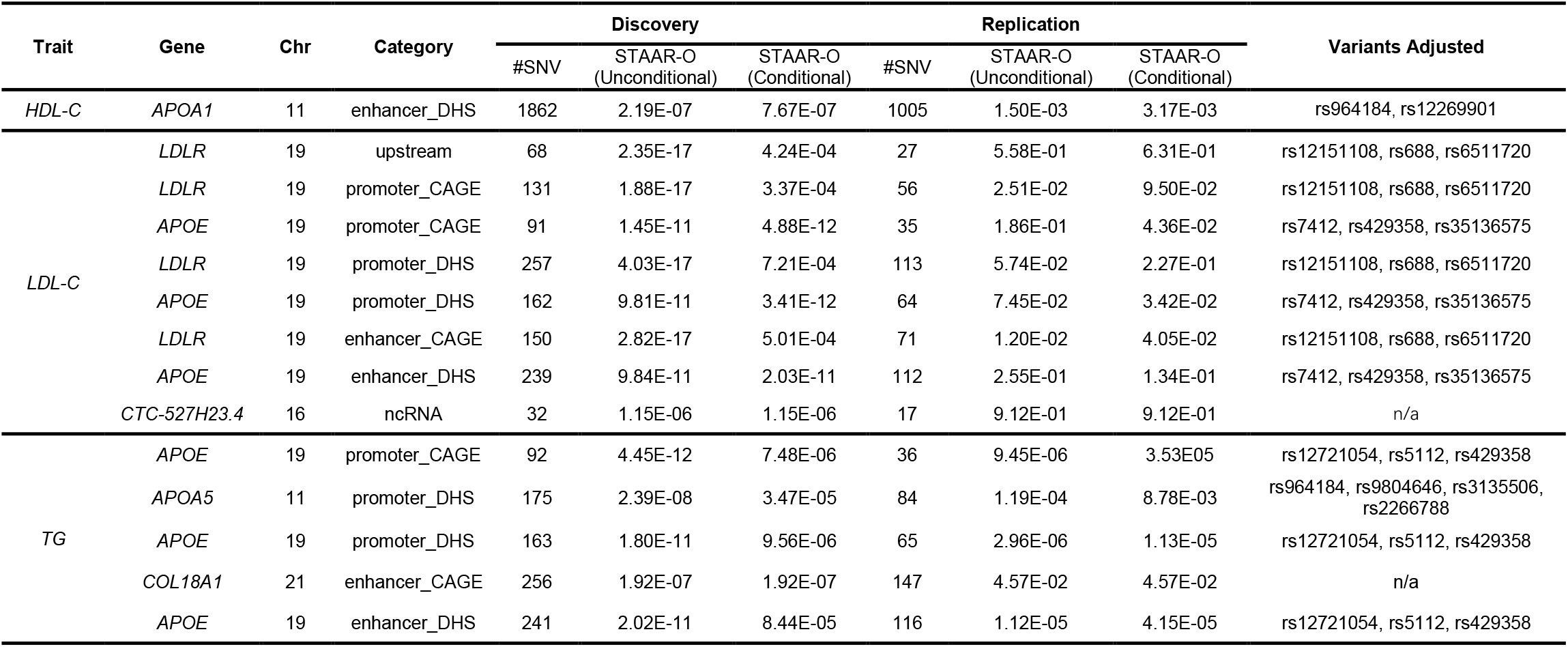
Gene-centric noncoding analysis results of both unconditional analysis and analysis conditional on known common and low-frequency variants. 21,015 discovery samples and 9,123 replication samples from the NHLBI Trans-Omics for Precision Medicine (TOPMed) program are considered in the analysis. Results for the conditionally significant genes (unconditional STAAR-O P < 3.57 × 10^−7^ and conditional STAAR-O P < 1.16 × 10^−3^ for 7 different noncoding masks across protein-coding genes; unconditional STAAR-O P < 2.50 × 10^−6^ and conditional STAAR-O P < 1.16 × 10^−3^ for ncRNA genes) using discovery samples are presented in the table. Chr (Chromosome); Category (Functional category); #SNV (Number of rare variants (MAF < 1%) of the particular functional category in the gene); STAAR-O (STAAR-O *P* value); HDL-C (High-density lipoprotein cholesterol); LDL-C (Low-density lipoprotein cholesterol); TG (Triglycerides); TC (Total cholesterol); Variants Adjusted (Adjusted variants in conditional analysis); n/a, no variant adjusted in the conditional analysis.

In unconditional analysis of the discovery data, using the 2-kb sliding window procedure we identified 140 windows as genome-wide significant (**Supplementary Table 8, Supplementary Figures 3e-f, 4e-f, 5e-f, 6e-f**). Among these 140 significant sliding windows, 14 are located in noncoding regions and, after conditioning on known lipid-associated variants, all remained significant at the Bonferroni-corrected level 0.05/140 = 3.57 × 10^−4^ (**Table 2**). In replication data 9 of the 14 associations were significant at the Bonferroni-corrected level 0.05/14 = 3.57 × 10^−3^ after adjusting for known phenotype-specific variants. When we further adjusted these 9 associations for known individual rare variants (MAC ≥ 20), associations for two intronic sliding windows (*PAFAH1B2* and TG) remained significant at the same level of 3.57 × 10^−3^ (**Supplementary Table 9**).

**Table 2.**
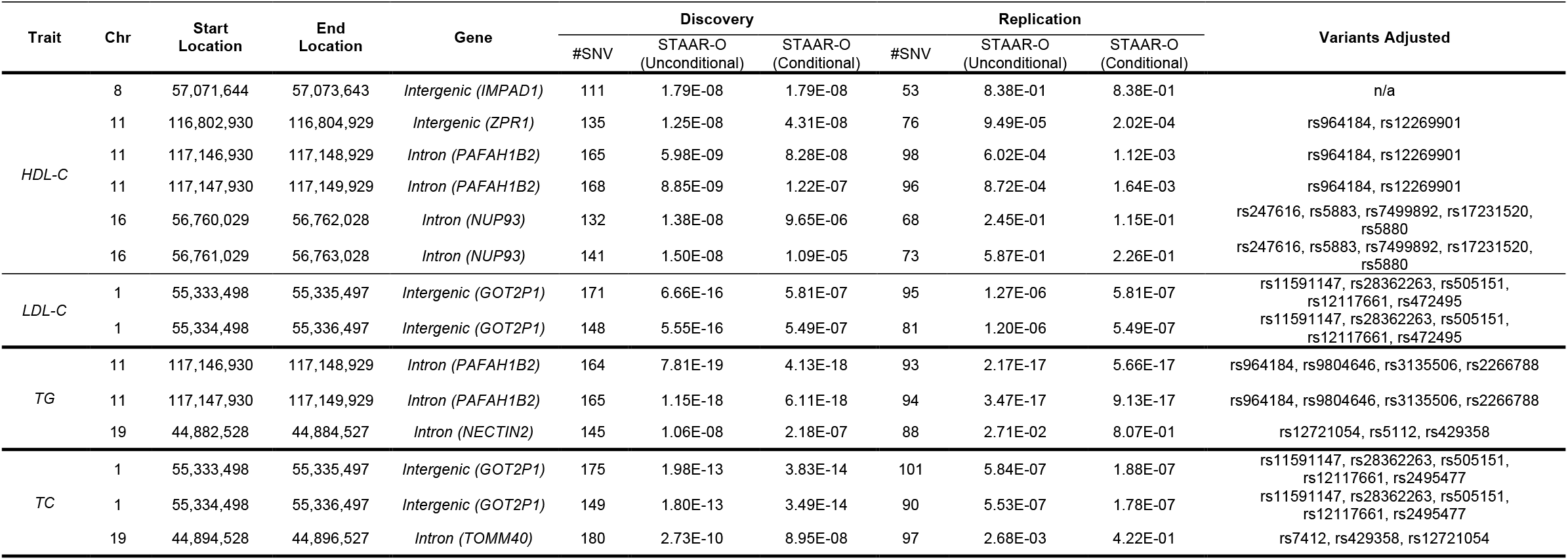
2-kb sliding window analysis results of unconditional analysis and analysis conditional on known common and low-frequency variants. 21,015 discovery samples and 9,123 replication samples from the NHLBI Trans-Omics for Precision Medicine (TOPMed) program are considered in the analysis. Results for the conditionally significant sliding windows (unconditional STAAR-O P < 1.88 × 10^−8^; conditional STAAR-O P < 3.57 × 10^−4^) using discovery samples are presented in the table. Chr (Chromosome); Start Location (Start location of the 2kb sliding window); End Location (End location of the 2-kb sliding window); # SNV (Number of rare variants (MAF < 1%) in the 2-kb sliding window; STAAR-O (STAAR-O *P* value); HDL-C (High-density lipoprotein cholesterol); LDL-C (Low-density lipoprotein cholesterol); TG (Triglycerides); TC (Total cholesterol); Variants Adjusted (Adjusted variants in conditional analysis); n/a, no variant adjusted in the conditional analysis. Physical positions of each window are on build hg38.

| Trait | Chr | Start Location | End Location | Gene | Discovery |  |  | Replication |  |  | Variants Adjusted |
| --- | --- | --- | --- | --- | --- | --- | --- | --- | --- | --- | --- |
|  |  |  |  |  | #SNV | STAAR-O (Unconditional) | STAAR-O (Conditional) | #SNV | STAAR-O (Unconditional) | STAAR-O (Conditional) |  |
| HDL-C | 8 | 57,071,644 | 57,073,643 | *Intergenic (IMPAD1)* | 111 | 1.79E-08 | 1.79E-08 | 53 | 8.38E-01 | 8.38E-01 | n/a |
|  | 11 | 116,802,930 | 116,804,929 | *Intergenic (ZPR1)* | 135 | 1.25E-08 | 4.31E-08 | 76 | 9.49E-05 | 2.02E-04 | rs964184, rs12269901 |
|  | 11 | 117,146,930 | 117,148,929 | *Intron (PAFAH1B2)* | 165 | 5.98E-09 | 8.28E-08 | 98 | 6.02E-04 | 1.12E-03 | rs964184, rs12269901 |
|  | 11 | 117,147,930 | 117,149,929 | *Intron (PAFAH1B2)* | 168 | 8.85E-09 | 1.22E-07 | 96 | 8.72E-04 | 1.64E-03 | rs964184, rs12269901 |
|  | 16 | 56,760,029 | 56,762,028 | *Intron (NUP93)* | 132 | 1.38E-08 | 9.65E-06 | 68 | 2.45E-01 | 1.15E-01 | rs247616, rs5883, rs7499892, rs17231520, rs5880 |
|  | 16 | 56,761,029 | 56,763,028 | *Intron (NUP93)* | 141 | 1.50E-08 | 1.09E-05 | 73 | 5.87E-01 | 2.26E-01 | rs247616, rs5883, rs7499892, rs17231520, rs5880 |
| LDL-C | 1 | 55,333,498 | 55,335,497 | *Intergenic (GOT2P1)* | 171 | 6.66E-16 | 5.81E-07 | 95 | 1.27E-06 | 5.81E-07 | rs11591147, rs28362263, rs505151, rs12117661, rs472495 |
|  | 1 | 55,334,498 | 55,336,497 | *Intergenic (GOT2P1)* | 148 | 5.55E-16 | 5.49E-07 | 81 | 1.20E-06 | 5.49E-07 | rs11591147, rs28362263, rs505151, rs12117661, rs472495 |
| TG | 11 | 117,146,930 | 117,148,929 | *Intron (PAFAH1B2)* | 164 | 7.81E-19 | 4.13E-18 | 93 | 2.17E-17 | 5.66E-17 | rs964184, rs9804646, rs3135506, rs2266788 |
|  | 11 | 117,147,930 | 117,149,929 | *Intron (PAFAH1B2)* | 165 | 1.15E-18 | 6.11E-18 | 94 | 3.47E-17 | 9.13E-17 | rs964184, rs9804646, rs3135506, rs2266788 |
|  | 19 | 44,882,528 | 44,884,527 | *Intron (NECTIN2)* | 145 | 1.06E-08 | 2.18E-07 | 88 | 2.71E-02 | 8.07E-01 | rs12721054, rs5112, rs429358 |
| TC | 1 | 55,333,498 | 55,335,497 | *Intergenic (GOT2P1)* | 175 | 1.98E-13 | 3.83E-14 | 101 | 5.84E-07 | 1.88E-07 | rs11591147, rs28362263, rs505151, rs12117661, rs2495477 |
|  | 1 | 55,334,498 | 55,336,497 | *Intergenic (GOT2P1)* | 149 | 1.80E-13 | 3.49E-14 | 90 | 5.53E-07 | 1.78E-07 | rs11591147, rs28362263, rs505151, rs12117661, rs2495477 |
|  | 19 | 44,894,528 | 44,896,527 | *Intron (TOMM40)* | 180 | 2.73E-10 | 8.95E-08 | 97 | 2.68E-03 | 4.22E-01 | rs7412, rs429358, rs12721054 |

In unconditional analysis of the discovery data using the dynamic window procedure SCANG-STAAR-S we identified 90 genome-wide significant associations (**Supplementary Table 10**). Among them, 10 are located in noncoding regions and remained significant at Bonferroni-corrected level 0.05/90 = 5.56 × 10^−4^ after conditioning on known lipid-associated variants (**Table 3**). In the replication data, and after adjusting for known phenotype-specific variants, 7 were significant at the Bonferroni-corrected level 0.05/10 = 5 × 10^−3^. After further adjustment for known individual rare variants (MAC ≥ 20), 3 associations remained significant, including RVs in an intronic region of *PAFAH1B2* and TG, RVs in an intronic region of *SIDT2* and TG, and RVs in an intronic region of *CEP164* and TG (**Supplementary Table 11**).

**Table 3.**
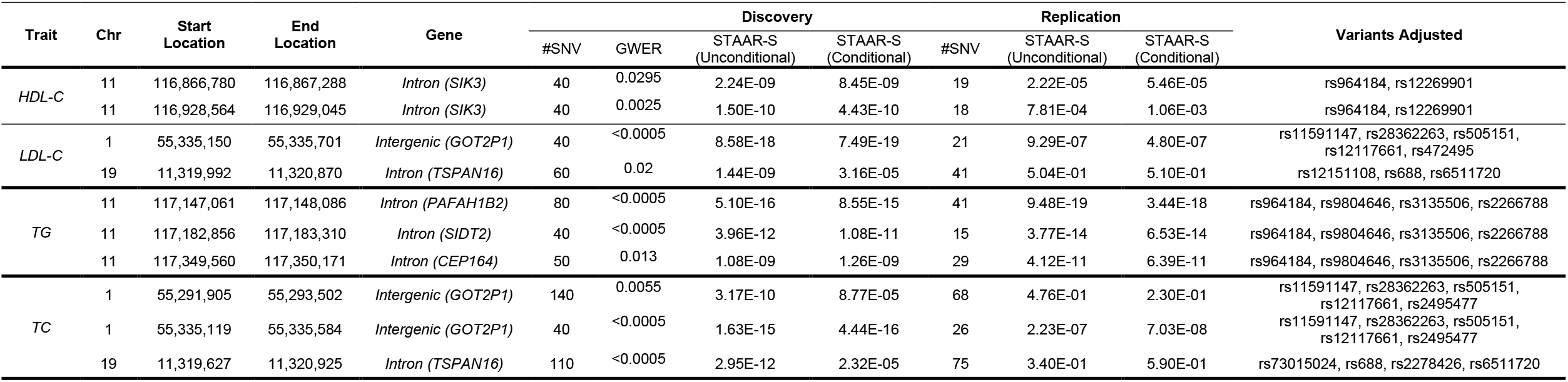
Dynamic window analysis results of unconditional analysis and analysis conditional on known common and low-frequency variants. 21,015 discovery samples and 9,123 replication samples from the NHLBI Trans-Omics for Precision Medicine (TOPMed) program are considered in the analysis. Results for the conditionally significant sliding windows (unconditional genome-wide error rate *G*WER < 0.05; conditional STAAR-S P < 5.56 × 10^−4^) using discovery samples are presented in the table. Chr (Chromosome); Start Location (Start location of the dynamic window); End Location (End location of the dynamic window); #SNV (Number of rare variants (MAF < 1%) in the dynamic window; GWER (genome-wide error rate); STAAR-S (STAAR-S *P* value); HDL-C (High-density lipoprotein cholesterol); LDL-C (Low-density lipoprotein cholesterol); TG (Triglycerides); TC (Total cholesterol); Variants Adjusted (Adjusted variants in conditional analysis). Physical positions of each window are on build hg38.

## Discussion

We developed a comprehensive association analysis framework for detecting noncoding rare variant set associations in large-scale WGS studies. Crucially, our framework explicitly solves the problem of defining variant sets, which is a significant challenge in practical analysis but not often discussed in other set-based inference methodology work. Our approach allows for continuous and binary traits and accounts for both population structure and relatedness through generalized linear mixed models using gene-centric analysis and non-gene-centric analysis. For gene-centric analysis, we proposed several strategies to define analysis units of rare variants in the noncoding genome, including seven genetic categories of regulatory regions for protein-coding genes, ncRNA genes, and perform RVATs of each noncoding mask using STAAR. For non-gene-centric analysis, to overcome the limitations of fixed-size sliding windows, we proposed SCANG-STAAR, a data-adaptive-size dynamic window scan procedure that incorporates multi-faceted functional annotations. We proposed *STAARpipeline* to perform RVATs using these methods for both noncoding and coding variants using unconditional analysis, as well as conditional analyses, which provides an analytical follow-up to distinguish novel RV association signals independent of known variants.

We developed *STAARpipeline*, a fast and resource-efficient tool for RV association analysis of WGS data that scales linearly on hundreds of thousands of samples, for both quantitative and dichotomous phenotypes. *STAARpipeline* allows researchers to conveniently functionally annotate a WGS/WES study using the variant functional annotation database FAVOR and the FAVORannotator workflow. *STAARpipeline* optimizes computational feasibility of RV association analysis in two steps. First, *STAARpipeline* reduces the computation burden of fitting the null mixed model using the estimated sparse GRM^16,38^. Second, *STAARpipeline* performs the RV association tests by taking advantage of sparse genotype dosages of RVs^39^.

In a WGS RV analysis of lipid traits in TOPMed, we identified and replicated using our *STAARpipeline* several conditional associations with lipid traits in the noncoding genome, including RVs in an intronic region of *PAFAH1B2* and TG, RVs in an intronic region of *SIDT2* and TG, and RVs in an intronic region of *CEP164* and TG, which were not detected by previous analysis of TOPMed Freeze 3 data^16,20^. Several coding rare variants in *PAFAH1B2* have been previously detected associated with TG^40^, our findings detected additionally significant RV association in the noncoding region of *PAFAH1B2*. Two intronic common variants in *SIDT2* have been reported associated with TG^41^, additional intronic rare variant association in *SIDT2* was detected using *STAARpipeline*.

For non-gene-centric analysis, we proposed improvements to the sliding window analysis using the dynamic window analysis of SCANG-STAAR. Compared with sliding window analysis using a fixed window size and skip length, SCANG-STAAR can increase power by considering all possible sub-windows of different sizes and selecting those windows that maximize power, while incorporating multi-faceted functional annotations. On the other hand, since SCANG-STAAR considers many more overlapping windows than the sliding window procedure, the genome-wide significance threshold is smaller than that of the sliding window procedure, potentially reducing power. For example, to control the genome-wise error rate at 0.05 level in our analysis of LDL-C, the *P* value threshold of SCANG-STAAR-S is 3.80 × 10^−9^ while the Bonferroni-corrected threshold of the 2-kb sliding window procedure is 1.88 × 10^−8^. When the window size of the signal region is close to the sliding window size, the sliding window procedure may detect associations missed by the dynamic window procedure because of this gap of the *P* value thresholds. In *STAARpipeline* we pragmatically provide both procedures.

In addition to noncoding rare variants association analysis, *STAARpipeline* also provides single variant analysis for common and low-frequency variants and gene-centric analysis for coding rare variants. The single variant analysis in *STAARpipeline* provides individual *P* values of variants given a MAF or MAC cut-off, for example, MAC ≥ 20. The gene-centric coding analysis provides five genetic categories to aggregate coding rare variants of each protein-coding gene: (1) putative loss of function (stop gain, stop loss and splice) RVs, (2) missense RVs, (3) disruptive missense RVs, (4) putative loss of function and disruptive missense RVs, and (5) synonymous RVs. The putative loss of function, missense, and synonymous RVs are defined by GENCODE VEP categories^29,30^. The disruptive variants are further defined by MetaSVM^42^, which measures the deleteriousness of missense mutations. As in the noncoding RV association analysis, single variant and gene-centric coding analyses also scale well in computation time and memory for large-scale WGS data. Using 30,138 related TOPMed samples these two analyses respectively took 3 hours and 5 hours for 100 cores with 6 Gb memory. Thus, *STAARpipeline* provides an efficient and comprehensive analysis tool for both coding and noncoding variant association discovery in large-scale sequencing studies.

With the emergence of large-scale WGS data, there is a pressing need to identify genetic components of complex traits in the noncoding genome. Here we introduce a powerful and scalable framework, *STAARpipeline*, for noncoding RV association detection across the genome. *STAARpipeline* provides several strategies to aggregate noncoding rare variants to empower RV association analysis in the noncoding region. We demonstrate the computational efficiency of *STAARpipeline* in application to the WGS association analysis of lipid traits on 30,138 TOPMed samples. The optimization approaches of *STAARpipeline* make it scalable for even larger data sets. Thus, our framework provides an essential solution for noncoding RV association detection in large-scale WGS data analysis and dissects the genetic contribution of noncoding rare variants to complex diseases.

## URLs

*STAARpipeline* (version 0.9.6), https://github.com/xihaoli/STAARpipeline and https://content.sph.harvard.edu/xlin/software.html.

*STAARpipelineSummary* (version 0.9.6), https://github.com/xihaoli/STAARpipelineSummary and https://content.sph.harvard.edu/xlin/software.html.

*FAVOR*, http://favor.genohub.org/.

*FAVORannotator*, https://github.com/zhouhufeng/FAVORannotator.

## Acknowledgments

This work was supported by grants R35-CA197449, P01-CA134294, U19-CA203654, R01-HL113338, and U01-HG009088 (X. Lin), R01-HL142711 and R01-HL127564 (P.N. and G.M.P), R35-HL135824 (C.J.W.), 75N92020D00001, HHSN268201500003I, N01-HC-95159, 75N92020D00005, N01-HC-95160, 75N92020D00002, N01-HC-95161, 75N92020D00003, N01-HC-95162, 75N92020D00006, N01-HC-95163, 75N92020D00004, N01-HC-95164, 75N92020D00007, N01-HC-95165, N01-HC-95166, N01-HC-95167, N01-HC-95168, N01-HC-95169, UL1-TR-000040, UL1-TR-001079, UL1-TR-001420, UL1TR001881, DK063491, R01HL071051, R01HL071205, R01HL071250, R01HL071251, R01HL071258, R01HL071259, and UL1RR033176 (J.I.R. and X.G.), U01-HL72518, HL087698, HL49762, HL59684, HL58625, HL071025, HL112064, NR0224103, and M01-RR000052 (to the Johns Hopkins General Clinical Research Center), NO1-HC-25195, HHSN268201500001I, 75N92019D00031, and R01-HL092577-06S1 (R.S.V. and L.A.C.), the Evans Medical Foundation and the Jay and Louis Coffman Endowment from the Department of Medicine, Boston University School of Medicine (R.S.V.), HHSN268201800001I and U01-HL137162 (K.M.R.), R01-HL133040 (D.E.W., M.S.R., and T.N.), R35-HL135818 and R01-HL113338 (S.R.), KL2TR002490 (L.M.R.), R01-HL92301, R01-HL67348, R01-NS058700, R01-AR48797, and R01-AG058921 (N.D.P. and D.W.B.), R01-DK071891 (N.D.P., B.I.F., and D.W.B.), M01-RR07122 and F32-HL085989 (to the General Clinical Research Center of the Wake Forest University School of Medicine), the American Diabetes Association, P60-AG10484 (to the Claude Pepper Older Americans Independence Center of Wake Forest University Health Sciences), U01-HL137181 (J.R.O.), HHSN268201600018C, HHSN268201600001C, HHSN268201600002C, HHSN268201600003C, and HHSN268201600004C (C.L.K.), R01-HL113323, U01-DK085524, R01-HL045522, R01-MH078143, R01-MH078111, and R01-MH083824 (H.H.H.G., R.D., J.E.C., and J.B.), 18CDA34110116 from American Heart Association (P.S.d.V.), HHSN268201800010I, HHSN268201800011I, HHSN268201800012I, HHSN268201800013I, HHSN268201800014I, and HHSN268201800015I (A.C.), HHSN268201700001I, HHSN268201700002I, HHSN268201700003I, HHSN268201700005I, and HHSN268201700004I (E.B.), U01-HL072524, R01-HL104135-04S1, U01-HL054472, U01-HL054473, U01-HL054495, U01-HL054509, and R01-HL055673-18S1 (D.K.A.). Molecular data for the Trans-Omics in Precision Medicine (TOPMed) program was supported by the National Heart, Lung and Blood Institute (NHLBI). Core support including centralized genomic read mapping and genotype calling, along with variant quality metrics and filtering were provided by the TOPMed Informatics Research Center (3R01HL-117626-02S1; contract HHSN268201800002I). Core support including phenotype harmonization, data management, sample-identity QC, and general program coordination were provided by the TOPMed Data Coordinating Center (R01HL-120393; U01HL-120393; contract HHSN268201800001I). We gratefully acknowledge the studies and participants who provided biological samples and data for TOPMed. We gratefully acknowledge the support from The Samoan Obesity, Lifestyle and Genetic Adaptations Study (OLaGA) Group. The full study specific acknowledgements are detailed in **Supplementary Note**.

## Author contributions

Z.L., X. Li and X. Lin designed the experiments., Z.L., X. Li, H.Z. and X. Lin performed the experiments. Z.L., X. Li, H.Z., S.M.G., M.S.S., T.A., C.Q., Y.L., H.C., R.S., R.D., D.K.A., L.F.B., J.C.B., T.W.B, J.B., E.B., D.W.B., J.A.B., B.E.C., M.P.C., A.C., L.A.C., J.E.C., P.S.d.V., R.D., B.I.F., H.H.H.G., X.G., R.R.K., C.L.K., B.G.K., L.A.L., A.W.M., L.W.M., B.D.M., M.E.M., A.C.M., T.N., J.R.O., N.D.P., P.A.P., B.M.P., L.M.R., S.R., A.P.R., M.S.R., K.M.R., S.S.R., J.A.S., K.D.T., R.S.V., D.E.W., J.G.W., L.R.Y., W.Z., J.I.R., C.J.W., P.N., G.M.P. and X. Lin acquired, analyzed or interpreted data. G.M.P., P.N. and NHLBI TOPMed Lipids Working Group provided administrative, technical or material support. Z.L., Li, S.M.G. and X. Lin drafted the manuscript and revised according to co-authors’ suggestions. All authors critically reviewed the manuscript, suggested revisions as needed, and approved the final version.

## Additional information

Supplementary information for this paper includes **Supplementary Figures** (6 figures), **Supplementary Tables** (11 tables) and **Supplementary Note**.

## Online Methods

### Notations and model

Suppose there are *n* subjects with *M* total variants sequenced across the whole genome. For subject *i*, let *Y*_*i*_ denote a continuous or dichotomous trait with mean *µ*_*i*_; ***X***_*i*_ = (*X*_*i*1_, …, *X*_*iq*_)^*T*^ denote *q* covariates, such as age, gender, ancestral principal components; and ***G***_*i*_ = (*G*_*i*1_, …, *G*_*iq*_)^*T*^ denote the genotype information of the *p* genetic variants in a given variant set.

We consider the Generalized Linear Model for unrelated samples,

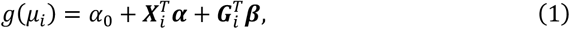

where *g* (*µ*) = *µ* for a continuous trait, *g* (*µ*) = logit(*µ*) for a dichotomous trait, *α*_0_ is an intercept, ***α*** = (*α*_1_, …, *α*_*q*_)^*T*^ is a vector of regression coefficients for ***X***_*i*_, and ***β***= (*β*_1_, …, *β*_*q*_)^*T*^ is a vector of regression coefficients for ***G***_*i*_.

We consider the following Generalized Linear Mixed Model^25,26,43^ for related samples,

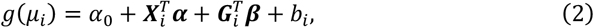

where the random effects *b*_*i*_ account for remaining population structure unaccounted by ancestral principal components and relatedness. Let ***b*** =(*b*_1_, …, *b*_*n*_)^*T*^ ∼ *N* (**0**, *θ***Ф**) with variance components *θ* and a genetic relatedness matrix **Ф**^16,38^. Our goal is testing the null hypothesis of whether the variant-set is associated with the phenotype, adjusting for covariates and relatedness, which corresponds to *H*_0_: ***β*** = 0, that is, *β*_1_ = *β*_2_ = … = *β*_*p*_ = 0.

### Variant set test using STAAR

The *STAARpipeline* calculates the variant set *P* value of each analysis unit using the STAAR method that incorporates multiple variant functional annotation scores^16^. Assume there are *K* annotations and 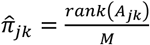, where *A*_*jk*_ is the *k*th annotation for the *j*th variant (*k* = 1, …, *K*; *j* = 1, …, *p*). For *k* = 0, we assume 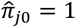. Assume *w*_*jl*_ = *Beta* (MAF_*j*_; *a*_1*l*_, *a*_2*l*_, where (*a*_11_, *a*_21_) = (1,25), (*a*_12_, *a*_22_) = (1,1) and MAF_*j*_ is the MAF of the *j*th variant (*j* = 1, …, *p*). The burden test statistic using *k*th variant functional annotation and *l*th beta density as the weight is given by

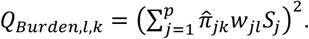

The SKAT test statistic using *k*th variant functional annotation and *l*th beta density as the weight is given by

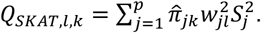

(*k* = 0, …, *K*; l = 1,2). The ACAT-V test statistic using *k*th variant functional annotation and *l*th beta density as the weight is given by

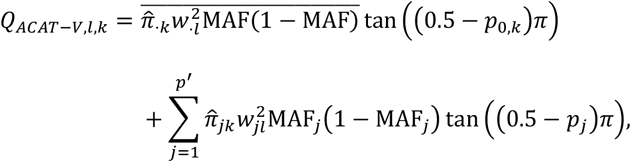

where 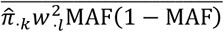 is the average of the weights 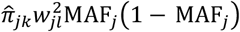 among the extremely rare variants with MAC ≤ 10, and *p*′ is the number of variants with MAC > 10 in the variant set.

Let *p*_*Burden,l,k*_ be the *P* value of *Q*_*Burden,l,k*_, *p*_*SKAT,l,k*_ be the *P* value of *Q*_*SKAT,l,k*_, and *p*_*ACAT*−*V,l,k*_ be the *P* value of *Q*_*ACAT*−*V,l,k*_ (*k* = 0, …, *K*; *l* = 1,2). We define STAAR-Burden (STAAR-B), STAAR-SKAT (STAAR-S), and STAAR-ACAT-V (STAAR-A) as 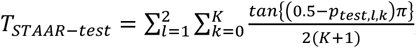, and the corresponding *P* value is calculated by 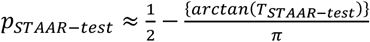, where *test* ∈{*Burden, SKAT, ACAT* − *V*}. The STAAR-O test statistic is defined as

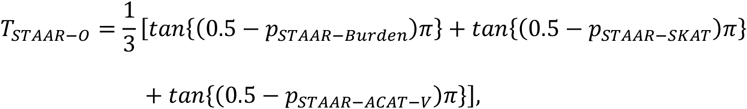

and the corresponding *P*-value is calculated by

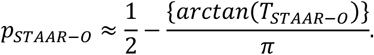

In gene-centric and sliding window analysis, we use the STAAR-O test for each analysis unit.

### Dynamic window analysis using SCANG-STAAR

The *STAARpipeline* performs dynamic window analysis using the SCANG-STAAR procedure, which extends the dynamic window rare variant test procedure SCANG by incorporating multiple variant functional annotations using the STAAR method. Under the global null hypothesis, there is no variant associated with the phenotype across the genome. Under the alternative hypothesis, there exists at least one region associated with the phenotype. SCANG-STAAR procedure provides a valid test by using the minimum value of the *P* value of all candidate moving windows of different sizes

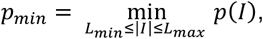

where *p*(*I*) is the *P* value of region *I*, |*I*| is the number of variants in a window *I*, and *L*_*min*_ and *L*_*max*_ are the smallest and largest number of variants in the searching windows, respectively. For SCANG-STAAR-S and SCANG-STAAR-B procedures, *p*(*I*) is the STAAR-S and STAAR-B *P* value of window *I*, respectively. For SCANG-STAAR-O, *p*(*I*) is the omnibus *P* value of STAAR-S and STAAR-B calculated by ACAT method^35^. Similar to the SCANG procedure, SCANG-STAAR controls the genome-wise type I error at a given *α* level by using the (1 − *α*)th quantile of the empirical distribution of *p*_*min*_ as an empirical threshold *h*(*α, p*_*min*_, *L*_*min*_, *L*_*max*_)^18^. We reject the null hypothesis if the *P* value of any window is smaller than *h*(*α, p*_*min*_, *L*_*min*_, L_max_). If this results in only one window, the detected window is 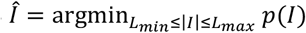. If this results in multiple overlapping windows, we localize the signals as the window whose *P* value is smaller than both the threshold and the windows that overlap with it.

### Conditional analysis

The *STAARpipeline* performs conditional analysis to identify RV association independent of known variants. We first select a list of known variants by including the trait-associated variants identified in literature, for example, variants indexed in GWAS Catalog^36^ or significant variants in large-scale GWAS. The significant variants detected in individual analysis using the same data could also be added into the known variants list to ensure the RV signals are not captured by the significant individual variants. We then use the following stepwise selection strategy to select a subset of independent variants representing the known variant list as the variants adjusted in the conditional analysis:

1. Calculate the individual *P* value of all variants in the known variants list and select the most significant variant.
2. For each step, calculate the *P* values of all the remaining variants conditional on the variant(s) that have already been selected. For each variant, we only condition on the selected variants within a specified region of that variant, such as the +/- 1-Mb window.
3. Select the variant with minimum conditional *P* value that is lower than the cutoff *P* value, for example, 1 × 10^−4^.
4. Repeat steps 2-3 until no variants can be selected.

Finally, we calculate the conditional *P* value of each significant RV analysis unit by adjusting for the selected variants residing in an extended region (for example, +/- 1-Mb window) of the analysis unit.

### Statistical analysis of lipid traits in the TOPMed data

The TOPMed WGS data consist of ancestrally diverse and multi-ethnic related samples^4,44^. Race/ethnicity was defined using a combination of self-reported race/ethnicity and study recruitment information (**Supplementary Note**)^37^. The discovery cohorts consist of 5,849 (27.8%) Black or African American, 12,313 (58.6%) White, 675 (3.2%) Asian American, 1,075 (5.1%) Hispanic/Latino American and 1,103 (5.3%) Samoans. The replication cohorts consist of 2,265 (24.8%) Black or African American, 5,615 (61.5%) White, and 1,243 (13.6%) Hispanic/Latino American.

We applied STAARpipeline to identify RV sets associated with four quantitative lipid traits (LDL-C, HDL-C, TG and TC) using the TOPMed WGS data. LDL-C and TC were adjusted for the presence of medications as before^20^. Linear regression model adjusting for age, age^2^, sex was first fit for each study-race/ethnicity-specific group. In addition, for Old Order Amish, we also adjusted for *APOB* p.R3527Q in LDL-C and TC analyses and adjusted for *APOC3* p.R19Ter in TG and HDL-C analyses^20^. The residuals were rank-based inverse normal transformed and rescaled by the standard deviation of the original phenotype within each group. We then fit a heteroscedastic linear mixed model (HLMM) for the rank normalized residuals, adjusting for 10 ancestral PCs, study-ethnicity group indicators, and a variance component for empirically derived kinship matrix plus separate group-specific residual variance components to account for population structure and relatedness. The output of HLMM was then used to perform following variant set analyses for rare variants (MAF < 1%) by scanning the genome, including gene-centric analysis using seven variant categories (promoter RVs overlaid with CAGE sites, promoter RVs overlaid with DHS sites, enhancer RVs overlaid with CAGE sites, enhancer RVs overlaid with DHS sites, UTR RVs, upstream RVs and downstream RVs) for each protein coded gene, ncRNA RVs, 2-kb sliding windows with 1-kb skip length, and dynamic windows with variants number between 40 and 300. The WGS RVAT analysis was performed using R packages STAAR (version 0.9.6), STAARpipeline (version 0.9.6) and STAARpipelineSummary (version 0.9.6).

### Genome build

All genome coordinates are given in NCBI GRCh38/UCSC hg38.

### Code availability

*STAARpipeline* is implemented as an open-source R package available at https://github.com/xihaoli/STAARpipeline and https://content.sph.harvard.edu/xlin/software.html. *STAARpipelineSummary* is implemented as an open-source R package available at https://github.com/xihaoli/STAARpipelineSummary and https://content.sph.harvard.edu/xlin/software.html.

### Data availability

This paper used the TOPMed Freeze 5 Whole Genome Sequencing data and lipids phenotype data. The genotype and phenotype data are both available in dbGAP. The discovery phase used the data from the following six study cohorts, where the accession numbers are provided in parenthesis: Framingham Heart Study (phs000974.v1.p1), Old Order Amish (phs000956.v1.p1), Jackson Heart Study (phs000964.v1.p1), Multi-Ethnic Study of Atherosclerosis (phs001416.v1.p1), Genome-wide Association Study of Adiposity in Samoans (phs000972) and Women’s Health Initiative (phs001237). The replication phase used the data from the following eight study cohorts: Atherosclerosis Risk in Communities Study (phs001211), Cleveland Family Study (phs000954), Cardiovascular Health Study (phs001368), Diabetes Heart Study (phs001412), Genetic Study of Atherosclerosis Risk (phs001218), Genetic Epidemiology Network of Arteriopathy (phs001345), Genetics of Lipid Lowering Drugs and Diet Network (phs001359) and San Antonio Family Heart Study (phs001215). The sample sizes, ethnicity and phenotype summary statistics of these cohorts are given in **Supplementary Table 3**.

The functional annotation data are publicly available and were downloaded from the following links: GRCh38 CADD v1.4 (https://cadd.gs.washington.edu/download), ANNOVAR dbNSFP v3.3a (https://annovar.openbioinformatics.org/en/latest/user-guide/download), LINSIGHT (https://github.com/CshlSiepelLab/LINSIGHT), FATHMM-XF (http://fathmm.biocompute.org.uk/fathmm-xf), CAGE (https://fantom.gsc.riken.jp/5/data), GeneHancer (https://www.genecards.org), and Umap/Bismap (https://bismap.hoffmanlab.org). In addition, recombination rate and nucleotide diversity were obtained from Gazal et al^45^. The tissue-specific functional annotations were downloaded from ENCODE (https://www.encodeproject.org/report/?type=Experiment).

