## Supplementary Files for "A framework for detecting noncoding rare variant associations of large-scale whole-genome sequencing studies"

Li et al

#### Supplementary Figures

**Supplementary Figure 1. Relatedness of subjects within and across studies in the discovery and replication samples of the TOPMed lipids study.** See **Supplementary Note** for study abbreviations.

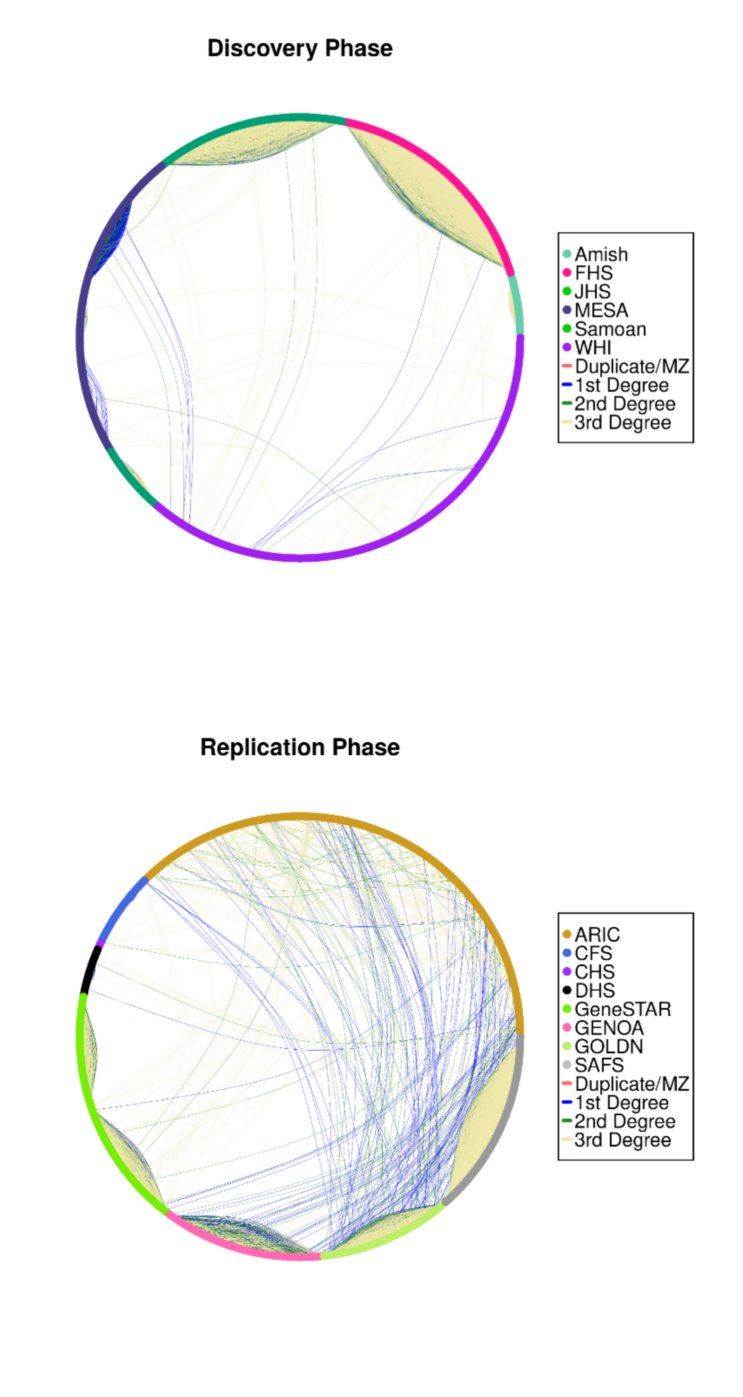

**Supplementary Figure 2. Rare variant (MAF < 0.01) distribution in the discovery phase using TOPMed cohorts (n=21,015).** Variant categories are defined by GENCODE VEP categories.

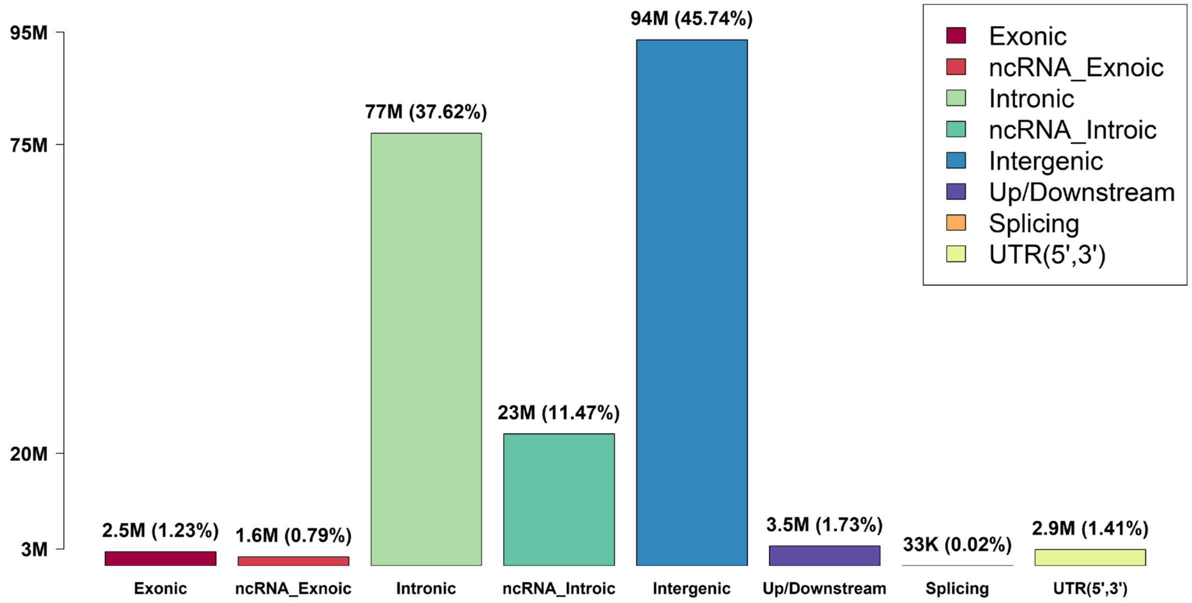

**Supplementary Figure 3. Manhattan plots and Q-Q plots for unconditional gene-centric noncoding analysis and sliding window analysis of high-density lipoprotein cholesterol (HDL-C) in the discovery phase (n=21,015). a**, Manhattan plots for unconditional gene-centric noncoding analysis of protein-coding gene. The horizontal line indicates a genome-wide STAAR-O *P*-value threshold of $3.57 \times 10^{-7}$. Different symbols represent the STAAR-O *P*-value of the protein-coding gene using different functional categories (upstream, downstream, UTR, promoter_CAGE, promoter_DHS, enhancer_CAGE, enhancer_DHS). Promoter_CAGE and promoter_DHS are the promoter with overlap of Cap Analysis of Gene Expression (CAGE) sites and DNase hypersensitivity (DHS) sites for a given gene, respectively. **b**, Quantile-quantile plots for unconditional gene-centric noncoding analysis of protein-coding gene. Different symbols represent the STAAR-O *P*-value of the gene using different functional categories (upstream, downstream, UTR, promoter_CAGE, promoter_DHS, enhancer_CAGE, enhancer_DHS). **c,** Manhattan plots for unconditional gene-centric noncoding analysis of ncRNA gene. The horizontal line indicates a genome-wide STAAR-O *P*-value threshold of $2.50 \times 10^{-6}$. **d,** Quantile-quantile plots for unconditional gene-centric noncoding analysis of ncRNA gene. **e,** Manhattan plot for 2-kb sliding windows. The horizontal line indicates a genome-wide *P*-value threshold of $1.88 \times 10^{-8}$. **f**, Quantile-quantile plot for 2-kb sliding windows.

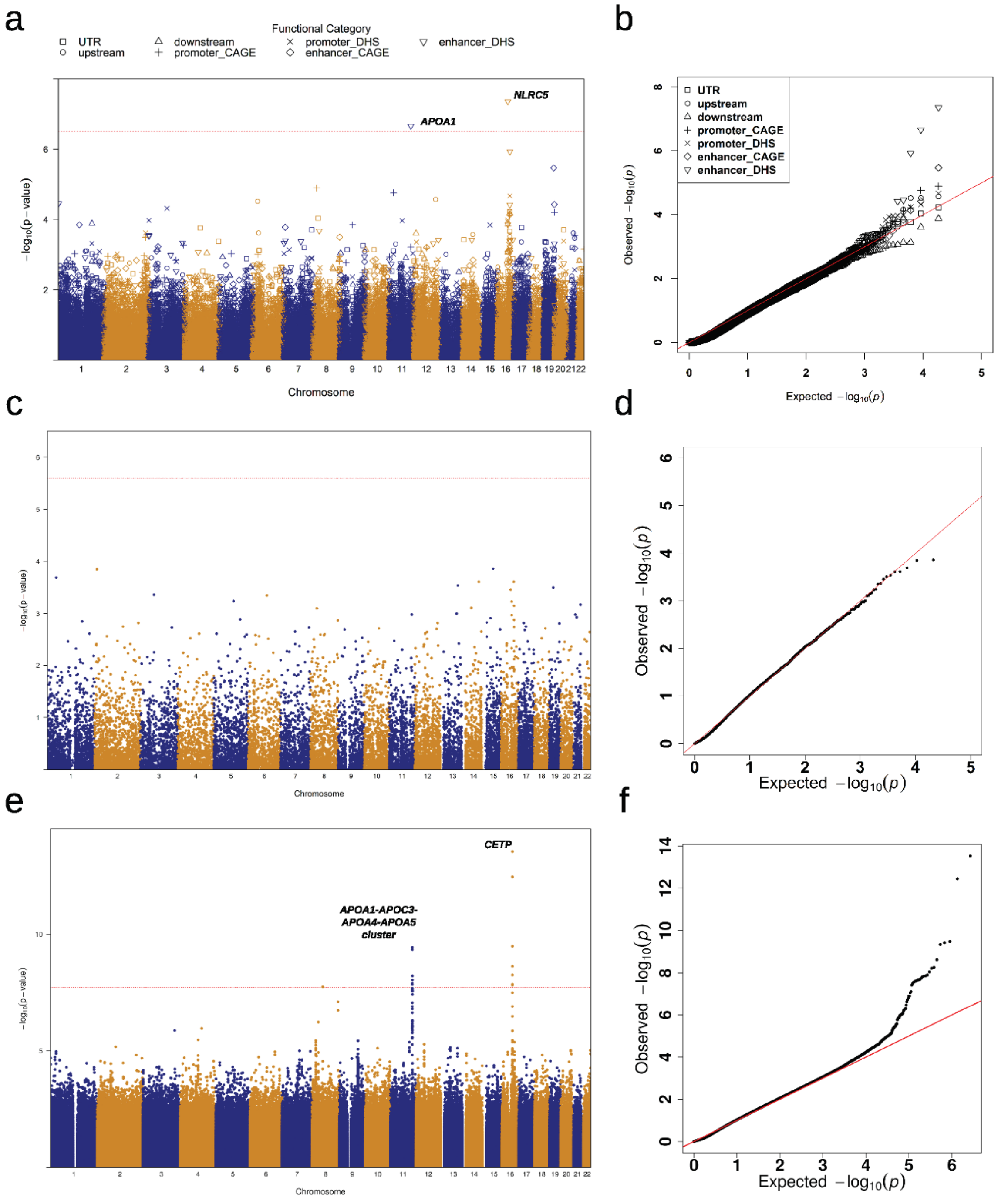

a
Functional Category
UTR
upstream
downstream
promoter_CAGE
promoter_DHS
enhancer_CAGE
enhancer_DHS
NLRC5
APOA1
−log10(p − value)
Chromosome
b
Observed −log10(p)
Expected −log10(p)
c
Chromosome
d
Observed −log10(p)
Expected −log10(p)
e
CETP
APOA1-APOC3-APOA4-APOA5 cluster
Chromosome
f
Observed −log10(p)
Expected −log10(p)

**Supplementary Figure 4. Manhattan plots and Q-Q plots for unconditional gene-centric noncoding analysis and sliding window analysis of low-density lipoprotein cholesterol (LDL-C) in the discovery phase (n=21,015). a**, Manhattan plots for unconditional gene-centric noncoding analysis of protein-coding gene. The horizontal line indicates a genome-wide STAAR-O *P*-value threshold of $3.57 \times 10^{-7}$. Different symbols represent the STAAR-O *P*-value of the protein-coding gene using different functional categories (upstream, downstream, UTR, promoter_CAGE, promoter_DHS, enhancer_CAGE, enhancer_DHS). Promoter_CAGE and promoter_DHS are the promoter with overlap of Cap Analysis of Gene Expression (CAGE) sites and DNase hypersensitivity (DHS) sites for a given gene, respectively. **b**, Quantile-quantile plots for unconditional gene-centric noncoding analysis of protein-coding gene. Different symbols represent the STAAR-O *P*-value of the gene using different functional categories (upstream, downstream, UTR, promoter_CAGE, promoter_DHS, enhancer_CAGE, enhancer_DHS). **c,** Manhattan plots for unconditional gene-centric noncoding analysis of ncRNA gene. The horizontal line indicates a genome-wide STAAR-O *P*-value threshold of $2.50 \times 10^{-6}$. **d,** Quantile-quantile plots for unconditional gene-centric noncoding analysis of ncRNA gene. **e,** Manhattan plot for 2-kb sliding windows. The horizontal line indicates a genome-wide *P*-value threshold of $1.88 \times 10^{-8}$. **f**, Quantile-quantile plot for 2-kb sliding windows.

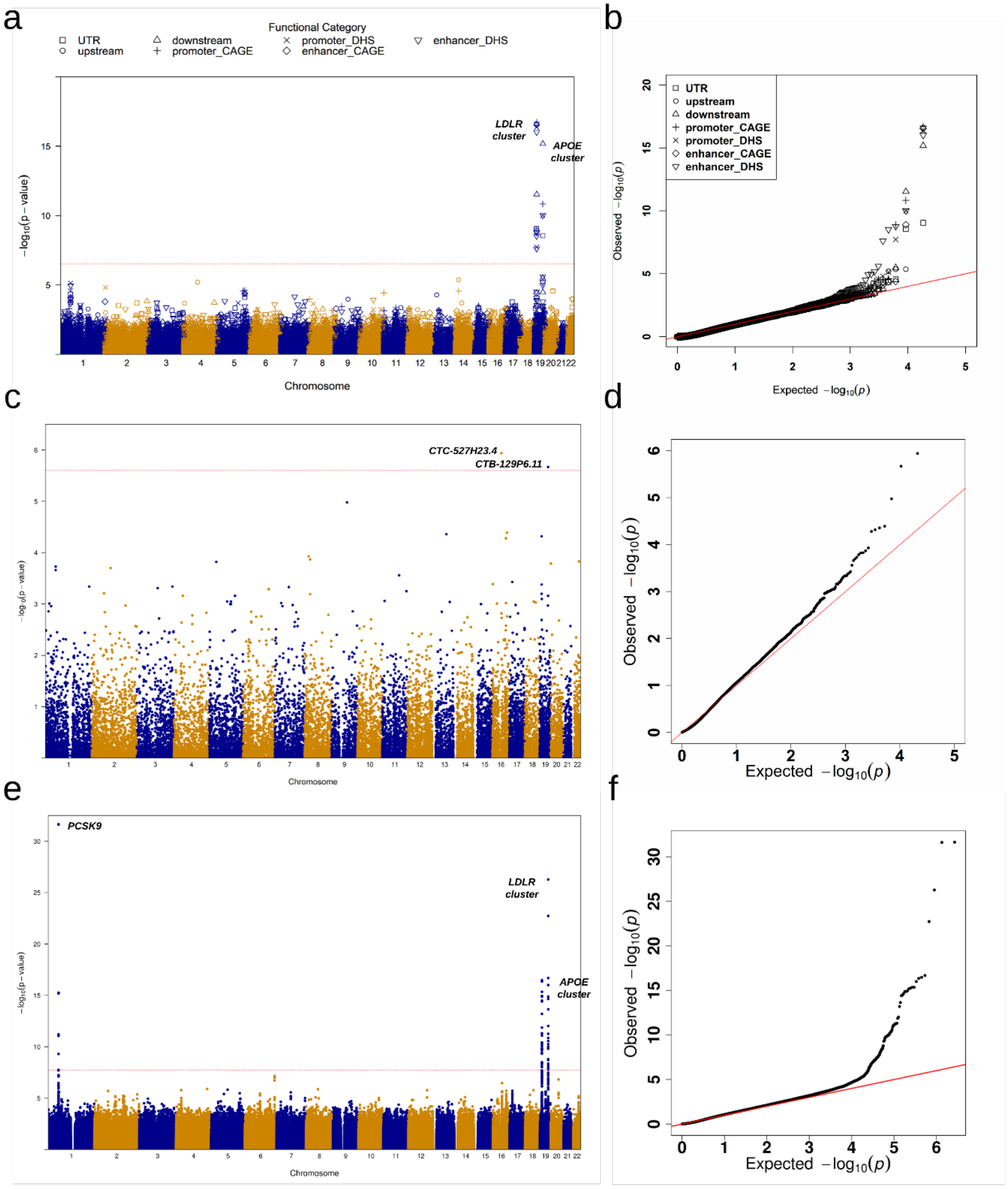
a
Functional Category
UTR
upstream
downstream
promoter_CAGE
promoter_DHS
enhancer_CAGE
enhancer_DHS
LDLR cluster
APOE cluster
$-\log_{10}$(p – value)
Chromosome
b
UTR
upstream
downstream
promoter_CAGE
promoter_DHS
enhancer_CAGE
enhancer_DHS
Observed $-\log_{10}(p)$
Expected $-\log_{10}(p)$
c
CTC-527H23.4
CTB-129P6.11
$-\log_{10}$(p – value)
Chromosome
d
Observed $-\log_{10}(p)$
Expected $-\log_{10}(p)$
e
PCSK9
LDLR cluster
APOE cluster
$-\log_{10}$(p – value)
Chromosome
f
Observed $-\log_{10}(p)$
Expected $-\log_{10}(p)$

**Supplementary Figure 5. Manhattan plots and Q-Q plots for unconditional gene-centric noncoding analysis and sliding window analysis of triglycerides (TG) in the discovery phase (n=21,015). a**, Manhattan plots for unconditional gene-centric noncoding analysis of protein-coding gene. The horizontal line indicates a genome-wide STAAR-O *P*-value threshold of $3.57 \times 10^{-7}$. Different symbols represent the STAAR-O *P*-value of the protein-coding gene using different functional categories (upstream, downstream, UTR, promoter_CAGE, promoter_DHS, enhancer_CAGE, enhancer_DHS). Promoter_CAGE and promoter_DHS are the promoter with overlap of Cap Analysis of Gene Expression (CAGE) sites and DNase hypersensitivity (DHS) sites for a given gene, respectively. **b**, Quantile-quantile plots for unconditional gene-centric noncoding analysis of protein-coding gene. Different symbols represent the STAAR-O *P*-value of the gene using different functional categories (upstream, downstream, UTR, promoter_CAGE, promoter_DHS, enhancer_CAGE, enhancer_DHS). **c,** Manhattan plots for unconditional gene-centric noncoding analysis of ncRNA gene. The horizontal line indicates a genome-wide STAAR-O *P*-value threshold of $2.50 \times 10^{-6}$. **d,** Quantile-quantile plots for unconditional gene-centric noncoding analysis of ncRNA gene. **e,** Manhattan plot for 2-kb sliding windows. The horizontal line indicates a genome-wide *P*-value threshold of $1.88 \times 10^{-8}$. **f**, Quantile-quantile plot for 2-kb sliding windows.

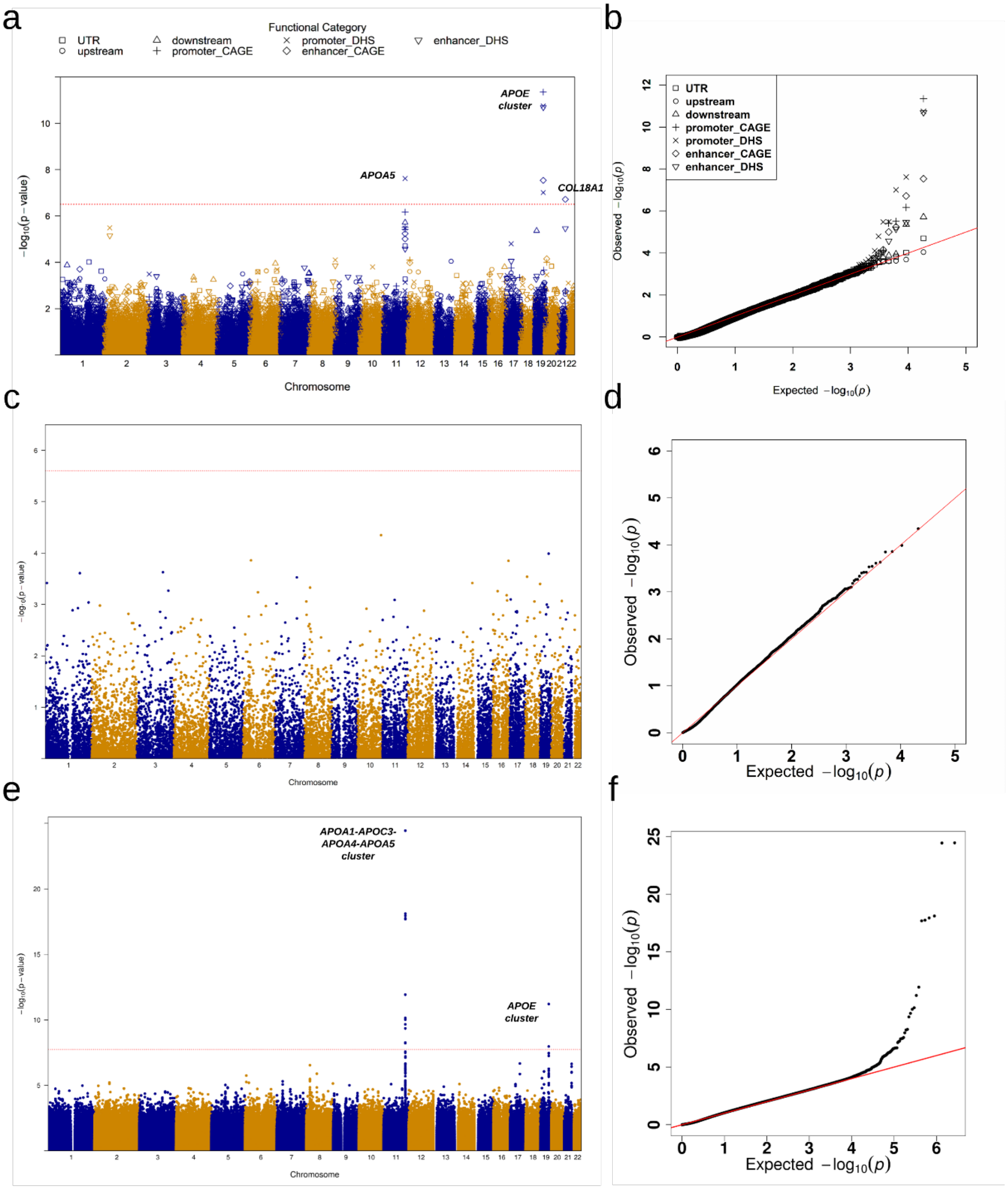

a
Functional Category
UTR
upstream
downstream
promoter_CAGE
promoter_DHS
enhancer_CAGE
enhancer_DHS
APOE cluster
APOA5
COL18A1
−log10(p − value)
Chromosome
b
UTR
upstream
downstream
promoter_CAGE
promoter_DHS
enhancer_CAGE
enhancer_DHS
Observed −log10(p)
Expected −log10(p)
c
Chromosome
d
Observed −log10(p)
Expected −log10(p)
e
APOA1-APOC3-APOA4-APOA5 cluster
APOE cluster
Chromosome
f
Observed −log10(p)
Expected −log10(p)

**Supplementary Figure 6. Manhattan plots and Q-Q plots for unconditional gene-centric noncoding analysis and sliding window analysis of total cholesterol (TC) in the discovery phase (n=21,015). a**, Manhattan plots for unconditional gene-centric noncoding analysis of protein-coding gene. The horizontal line indicates a genome-wide STAAR-O *P*-value threshold of $3.57 \times 10^{-7}$. Different symbols represent the STAAR-O *P*-value of the protein-coding gene using different functional categories (upstream, downstream, UTR, promoter_CAGE, promoter_DHS, enhancer_CAGE, enhancer_DHS). Promoter_CAGE and promoter_DHS are the promoter with overlap of Cap Analysis of Gene Expression (CAGE) sites and DNase hypersensitivity (DHS) sites for a given gene, respectively. **b**, Quantile-quantile plots for unconditional gene-centric noncoding analysis of protein-coding gene. Different symbols represent the STAAR-O *P*-value of the gene using different functional categories (upstream, downstream, UTR, promoter_CAGE, promoter_DHS, enhancer_CAGE, enhancer_DHS). **c,** Manhattan plots for unconditional gene-centric noncoding analysis of ncRNA gene. The horizontal line indicates a genome-wide STAAR-O *P*-value threshold of $2.50 \times 10^{-6}$. **d,** Quantile-quantile plots for unconditional gene-centric noncoding analysis of ncRNA gene. **e,** Manhattan plot for 2-kb sliding windows. The horizontal line indicates a genome-wide *P*-value threshold of $1.88 \times 10^{-8}$. **f**, Quantile-quantile plot for 2-kb sliding windows.

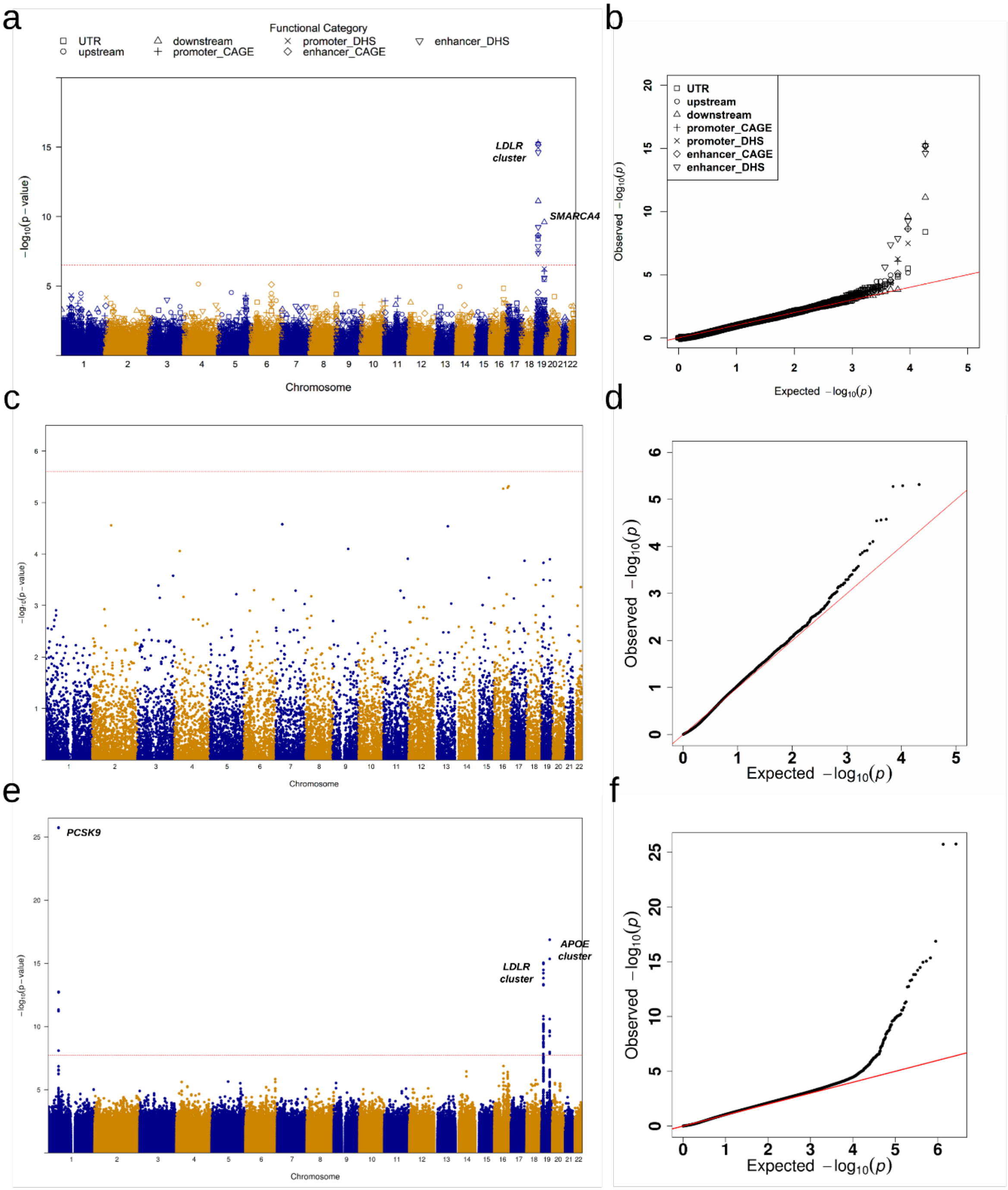
a
Functional Category
UTR
upstream
downstream
promoter_CAGE
promoter_DHS
enhancer_CAGE
enhancer_DHS
LDLR cluster
SMARCA4
−log10(p−value)
Chromosome
b
UTR
upstream
downstream
promoter_CAGE
promoter_DHS
enhancer_CAGE
enhancer_DHS
Observed −log10(p)
Expected −log10(p)
c
Chromosome
d
Observed −log10(p)
Expected −log10(p)
e
PCSK9
LDLR cluster
APOE cluster
Chromosome
f
Observed −log10(p)
Expected −log10(p)

**Supplementary Table 1.** Individual and integrative variant functional annotations used in the STAAR framework. Tissue-specific annotations (liver) DNase, H3K4me3, H3K27ac and H3K27me3 are from ENCODE (https://www.encodeproject.org/report/?type=Experiment). Each annotation Principal Component (aPC) is the first PC calculated from the set of individual functional annotations that measure similar biological function. These aPCs are then transformed into the PHRED-scaled scores (-10*log10(rank/total)) for each variant across the genome. For aPC-LocalDiversity, we consider two separate scores generated by ranks from both directions.

| | **Annotation type** | **Individual annotation scores used to calculate annotation PCs** |
|---|---|---|
| annotation PCs (aPCs) | aPC-EpigeneticActive | EncodeH3K4me1.max, EncodeH3K4me2.max, EncodeH3K4me3.max, EncodeH3K9ac.max, EncodeH3K27ac.max, EncodeH4K20me1.max, EncodeH2AFZ.max |
| | aPC-EpigeneticRepressed | EncodeH3K9me3.max, EncodeH3K27me3.max |
| | aPC-EpigeneticTranscription | EncodeH3K36me3.max, EncodeH3K79me2.max |
| | aPC-Conservation | GerpN, priPhCons, mamPhCons, verPhCons, priPhyloP, mamPhyloP, verPhyloP |
| | aPC-Protein | SIFTval, PolyPhenVal, Grantham, Polyphen2_HDIV_score, Polyphen2_HVAR_score, MutationTaster_score, MutationAssessor_score |
| | aPC-LocalDiversity (+/-) | bStatistic, RecombinationRate, NucleotideDiversity, Freq100bp, Rare100bp, Sngl100bp, Freq1000bp, Rare1000bp, Sngl1000bp, Freq10000bp, Rare10000bp, Sngl10000bp |
| | aPC-TF | RemapOverlapTF, RemapOverlapCL |
| | aPC-Mappability | umap_k100, bismap_k100, umap_k50, bismap_k50, umap_k36, bismap_k36, umap_k24, bismap_k24 |
| | aPC-Liver | DNase (ENCFF032OQN), H3K4me3 (ENCFF958VCA), H3K27ac (ENCFF035NGT), H3K27me3 (ENCFF179RUF) |
| Integrative scores | CADD | |
| | LINSIGHT | |
| | FATHMM-XF | |
| | MetaSVM | |

**Supplementary Table 2.** Study-specific sample sizes and baseline characteristics for 21,015 discovery samples and 9,123 replication samples of study participants. LDL-C (low-density lipoprotein cholesterol); HDL-C (high-density lipoprotein cholesterol); TG (triglycerides); TC (total cholesterol). Count - N; Continuous - Mean (Standard Deviation); Triglycerides are summarized as median [IQR].

| Phase | Study | Age | Female | LDL-C (mg/dl) | HDL-C (mg/dl) | TG (mg/dl) | TC (mg/dl) | White | Black or African American | Asian American | Hispanic/Latino American | Samoan | Sample size |
|---|---|---|---|---|---|---|---|---|---|---|---|---|---|
| Discovery | FHS | 40 (11) | 1922 | 121 (35) | 53 (16) | 89 [61,142] | 198 (39) | 3598 | 0 | 0 | 0 | 0 | 3598 |
| | JHS | 56 (13) | 1821 | 126 (37) | 52 (15) | 91 [65,128] | 199 (41) | 0 | 2924 | 0 | 0 | 0 | 2924 |
| | MESA | 61 (10) | 2510 | 118 (32) | 51 (15) | 107 [76,156] | 194 (35) | 1665 | 1656 | 543 | 927 | 0 | 4791 |
| | OOA | 50 (17) | 503 | 141 (44) | 56 (16) | 63 [45,96] | 212 (47) | 1003 | 0 | 0 | 0 | 0 | 1003 |
| | Samoan | 45 (11) | 673 | 130 (32) | 45 (11) | 106 [77,148] | 199 (37) | 0 | 0 | 0 | 0 | 1103 | 1103 |
| | WHI | 67 (7) | 7596 | 146 (38) | 55 (15) | 130 [93,181] | 231 (41) | 6047 | 1269 | 132 | 148 | 0 | 7596 |
| Replication | ARIC | 55 (6) | 1760 | 139 (39) | 50 (17) | 116 [84,165] | 216 (42) | 3226 | 193 | 0 | 0 | 0 | 3419 |
| | CFS | 46 (16) | 283 | 100 (33) | 43 (13) | 100 [75,143] | 168 (38) | 218 | 273 | 0 | 12 | 0 | 503 |
| | CHS | 73 (5) | 29 | 125 (26) | 56 (19) | 127 [93,161] | 209 (31) | 49 | 0 | 0 | 0 | 0 | 49 |
| | DHS | 60 (9) | 193 | 107 (33) | 49 (14) | 99 [76,145] | 180 (40) | 325 | 0 | 0 | 0 | 0 | 325 |
| | GeneSTAR | 42 (11) | 963 | 128 (40) | 53 (15) | 103 [72,150] | 205 (44) | 904 | 725 | 0 | 0 | 0 | 1629 |
| | GENOA | 57 (11) | 756 | 121 (43) | 56 (18) | 124 [95,170] | 205 (46) | 0 | 1074 | 0 | 0 | 0 | 1074 |
| | GOLDN | 48 (16) | 472 | 121 (31) | 47 (13) | 108 [72,171] | 189 (38) | 893 | 0 | 0 | 0 | 0 | 893 |
| | SAFS | 43 (17) | 716 | 109 (32) | 50 (14) | 115 [82,166] | 184 (39) | 0 | 0 | 0 | 1231 | 0 | 1231 |

**Supplementary Table 3.** Ethinicity distribution, related sample distribution, and variant number distribution for discovery phase, replication phase and pooled samples of TOPMed Freeze 5 data. Common variants (MAF > 5%); low frequency variants (1% ≤ MAF ≤ 5%); rare variants (MAF < 1%). The “Others” category in the replication cohort includes many Hispanic/Latino American as well as a cohort of Samoans.

| | Race/Ethnicity Distribution | | | | | | Related Sample Distribution | | | Variant Number Distribution | | | | |
|---|---|---|---|---|---|---|---|---|---|---|---|---|---|---|
| Phase | White | Black or African American | Asian American | Hispanic/Latino American | Samoan | Total Sample Size | 1st Degree | 2nd Degree | 3rd Degree | Common | Low Frequency | Rare | Rare Noncoding | Total |
| Discovery | 12,313 (58.6%) | 5,849 (27.8%) | 675 (3.2%) | 1,075 (5.1%) | 1,103 (5.3%) | 21,015 | 3,610 (17.2%) | 546 (2.6%) | 472 (2.2%) | 6,250,357 | 4,855,695 | 204,767,089 | 202,213,403 | 215,873,141 |
| Replication | 5,615 (61.5%) | 2,265 (24.8%) | 0 (0.0%) | 1,243 (13.6%) | 0 (0.0%) | 9,123 | 2,819 (30.9%) | 361 (4.0%) | 225 (2.5%) | 6,250,193 | 4,950,351 | 108,794,989 | 107,460,347 | 119,995,533 |
| Pooled | 17,928 (59.5%) | 8,114 (26.9%) | 675 (2.2%) | 2,318 (7.7%) | 1103 (3.7%) | 30,138 | 6,690 (22.2%) | 938 (3.1%) | 769 (2.6%) | 6,252,506 | 4,869,126 | 244,025,643 | 240,976,748 | 255,147,275 |

**Supplementary Table 4**. Known variants list in conditional analysis. The list is generated using the stepwise algorithm decribed in the manuscript and the common and low-frequent phenotype-specific variants (MAF ≥ 1%) indexed in GWAS Catalog.

| Trait | CHR | POS | REF | ALT | #rs |
|---|---|---|---|---|---|
| HDL | 1 | 109,274,968 | G | T | rs12740374 |
| HDL | 1 | 230,162,032 | A | T | rs2281718 |
| HDL | 2 | 21,008,652 | G | A | rs676210 |
| HDL | 2 | 164,694,691 | T | C | rs7607980 |
| HDL | 3 | 36,919,169 | C | A | rs7622114 |
| HDL | 4 | 109,648,464 | C | T | rs78025076 |
| HDL | 5 | 73,630,689 | A | G | rs6881956 |
| HDL | 6 | 43,792,590 | C | G | rs11967262 |
| HDL | 7 | 80,671,133 | T | G | rs3211938 |
| HDL | 7 | 130,747,722 | T | C | rs6971365 |
| HDL | 8 | 9,326,086 | A | G | rs4841132 |
| HDL | 8 | 19,961,928 | A | G | rs326 |
| HDL | 8 | 19,973,410 | C | T | rs10096633 |
| HDL | 8 | 20,070,502 | T | A | rs6999158 |
| HDL | 8 | 120,856,311 | G | T | rs4871137 |
| HDL | 8 | 125,495,147 | C | A | rs2954038 |
| HDL | 9 | 15,304,784 | C | A | rs686030 |
| HDL | 9 | 104,826,853 | A | T | rs4149310 |
| HDL | 9 | 104,884,738 | C | T | rs11789603 |
| HDL | 9 | 104,902,020 | C | T | rs1883025 |
| HDL | 10 | 45,523,383 | A | G | rs11239549 |
| HDL | 11 | 61,824,890 | A | G | rs174566 |
| HDL | 11 | 116,778,201 | G | C | rs964184 |
| HDL | 11 | 117,103,213 | G | C | rs12269901 |
| HDL | 12 | 57,398,797 | C | T | rs11613352 |
| HDL | 12 | 123,975,620 | G | T | rs4765127 |
| HDL | 12 | 124,853,983 | C | T | rs10773112 |
| HDL | 13 | 44,950,491 | G | A | rs9526023 |
| HDL | 14 | 104,806,341 | A | T | rs45490496 |
| HDL | 15 | 43,528,519 | C | T | rs55707100 |
| HDL | 15 | 58,289,171 | T | A | rs12148399 |
| HDL | 15 | 58,391,167 | A | G | rs1532085 |
| HDL | 15 | 58,431,476 | C | T | rs1800588 |
| HDL | 16 | 56,955,678 | C | T | rs247616 |
| HDL | 16 | 56,961,915 | G | A | rs17231520 |
| HDL | 16 | 56,972,678 | C | T | rs7499892 |
| HDL | 16 | 56,973,441 | C | T | rs5883 |
| HDL | 16 | 56,981,179 | G | C | rs5880 |
| HDL | 16 | 67,943,479 | T | C | rs1109166 |
| HDL | 17 | 43,848,758 | C | T | rs72836561 |

| | | | | | |
|---|---|---|---|---|---|
| HDL | 18 | 49,592,028 | T | C | rs9958734 |
| HDL | 18 | 49,645,518 | C | G | rs8086351 |
| HDL | 19 | 8,364,439 | G | A | rs116843064 |
| HDL | 19 | 11,235,874 | C | T | rs3760782 |
| HDL | 19 | 44,908,684 | T | C | rs429358 |
| HDL | 19 | 44,909,976 | G | T | rs1065853 |
| HDL | 19 | 44,945,208 | T | G | rs5167 |
| HDL | 20 | 44,413,724 | C | T | rs1800961 |
| HDL | 20 | 45,923,216 | T | C | rs6073958 |
| HDL | 21 | 42,298,682 | A | G | rs3746915 |
| HDL | 22 | 21,622,645 | T | C | rs7444 |
| LDL | 1 | 55,021,673 | C | G | rs12117661 |
| LDL | 1 | 55,039,974 | G | T | rs11591147 |
| LDL | 1 | 55,055,640 | G | T | rs472495 |
| LDL | 1 | 55,058,182 | G | A | rs28362263 |
| LDL | 1 | 55,063,514 | G | A | rs505151 |
| LDL | 1 | 109,274,968 | G | T | rs12740374 |
| LDL | 1 | 234,717,312 | C | T | rs556107 |
| LDL | 2 | 21,011,100 | T | C | rs533617 |
| LDL | 2 | 21,041,028 | G | A | rs1367117 |
| LDL | 2 | 21,065,354 | G | A | rs563290 |
| LDL | 2 | 43,847,292 | C | T | rs4245791 |
| LDL | 3 | 32,496,755 | C | T | rs3773777 |
| LDL | 4 | 68,475,769 | T | C | rs976058 |
| LDL | 5 | 75,355,259 | A | G | rs3846662 |
| LDL | 5 | 156,971,158 | G | C | rs1501908 |
| LDL | 6 | 160,589,086 | A | G | rs10455872 |
| LDL | 7 | 44,566,618 | C | G | rs217381 |
| LDL | 8 | 9,323,885 | A | G | rs2169387 |
| LDL | 8 | 22,069,949 | C | G | rs7386762 |
| LDL | 8 | 125,466,208 | T | C | rs2001846 |
| LDL | 9 | 133,279,294 | T | G | rs495828 |
| LDL | 10 | 112,190,660 | T | C | rs7096937 |
| LDL | 11 | 61,803,910 | G | A | rs174549 |
| LDL | 11 | 116,778,201 | G | C | rs964184 |
| LDL | 12 | 120,951,159 | A | C | rs2650000 |
| LDL | 13 | 113,844,399 | C | T | rs6602911 |
| LDL | 14 | 24,413,852 | C | T | rs2332328 |
| LDL | 14 | 94,380,925 | T | A | rs17580 |
| LDL | 15 | 63,500,058 | T | C | rs56369308 |
| LDL | 16 | 72,063,928 | C | T | rs3794695 |
| LDL | 16 | 72,182,890 | C | A | rs7200153 |
| LDL | 17 | 66,214,462 | A | C | rs1801689 |
| LDL | 18 | 9,526,186 | T | C | rs328996 |
| LDL | 19 | 11,086,585 | G | A | rs12151108 |

| | | | | | |
|---|---|---|---|---|---|
| LDL | 19 | 11,091,630 | G | T | rs6511720 |
| LDL | 19 | 11,116,926 | C | T | rs688 |
| LDL | 19 | 19,349,732 | G | C | rs73001065 |
| LDL | 19 | 44,908,684 | T | C | rs429358 |
| LDL | 19 | 44,908,822 | C | T | rs7412 |
| LDL | 19 | 44,935,906 | C | G | rs35136575 |
| LDL | 19 | 48,703,160 | A | G | rs492602 |
| LDL | 20 | 41,095,698 | T | C | rs6065311 |
| LDL | 21 | 15,214,362 | T | A | rs12106385 |
| LDL | 22 | 46,231,706 | C | T | rs4253772 |
| TG | 1 | 62,612,551 | A | T | rs10889348 |
| TG | 1 | 230,161,390 | C | T | rs2281721 |
| TG | 2 | 21,002,409 | C | T | rs1042034 |
| TG | 2 | 27,508,073 | T | C | rs1260326 |
| TG | 2 | 164,694,691 | T | C | rs7607980 |
| TG | 3 | 52,482,277 | C | G | rs6800707 |
| TG | 4 | 87,136,201 | G | A | rs1408 |
| TG | 5 | 131,337,265 | G | A | rs193735 |
| TG | 5 | 156,963,286 | T | C | rs6882076 |
| TG | 6 | 43,790,159 | C | A | rs998584 |
| TG | 6 | 139,518,361 | G | C | rs608736 |
| TG | 7 | 73,602,532 | T | C | rs13240994 |
| TG | 8 | 9,326,086 | A | G | rs4841132 |
| TG | 8 | 19,961,928 | A | G | rs326 |
| TG | 8 | 19,965,681 | T | C | rs3289 |
| TG | 8 | 20,070,649 | G | A | rs10106652 |
| TG | 8 | 125,495,066 | T | C | rs2980888 |
| TG | 9 | 84,002,350 | A | G | rs1982151 |
| TG | 10 | 63,364,338 | C | G | rs10822163 |
| TG | 11 | 61,830,500 | A | G | rs1535 |
| TG | 11 | 116,778,201 | G | C | rs964184 |
| TG | 11 | 116,789,970 | G | A | rs2266788 |
| TG | 11 | 116,791,691 | G | C | rs3135506 |
| TG | 11 | 116,794,363 | C | T | rs9804646 |
| TG | 12 | 21,178,615 | T | C | rs4149056 |
| TG | 13 | 113,844,399 | C | T | rs6602911 |
| TG | 14 | 103,824,476 | A | G | rs12893623 |
| TG | 15 | 42,429,647 | G | A | rs184334219 |
| TG | 15 | 43,735,687 | T | C | rs139974673 |
| TG | 15 | 58,387,979 | T | C | rs261291 |
| TG | 16 | 56,956,804 | C | A | rs247617 |
| TG | 16 | 81,501,185 | T | C | rs2925979 |
| TG | 17 | 43,763,481 | C | T | rs77697917 |
| TG | 18 | 289,209 | C | T | rs6506033 |
| TG | 19 | 8,364,439 | G | A | rs116843064 |

| | | | | | |
|---|---|---|---|---|---|
| TG | 19 | 19,321,481 | AG | A | rs140868651 |
| TG | 19 | 44,908,684 | T | C | rs429358 |
| TG | 19 | 44,919,330 | A | G | rs12721054 |
| TG | 19 | 44,927,023 | C | G | rs5112 |
| TG | 19 | 48,756,272 | A | G | rs838133 |
| TG | 20 | 41,152,292 | G | A | rs6093446 |
| TG | 20 | 45,923,216 | T | C | rs6073958 |
| TG | 21 | 39,181,919 | T | C | rs6517522 |
| TG | 22 | 38,204,535 | C | T | rs2267373 |
| TC | 1 | 23,421,503 | GA | G | rs11340914 |
| TC | 1 | 55,021,673 | C | G | rs12117661 |
| TC | 1 | 55,039,974 | G | T | rs11591147 |
| TC | 1 | 55,052,794 | A | G | rs2495477 |
| TC | 1 | 55,058,182 | G | A | rs28362263 |
| TC | 1 | 55,063,514 | G | A | rs505151 |
| TC | 1 | 62,620,326 | T | A | rs12239736 |
| TC | 1 | 109,274,968 | G | T | rs12740374 |
| TC | 1 | 234,722,850 | A | T | rs514230 |
| TC | 2 | 21,011,100 | T | C | rs533617 |
| TC | 2 | 21,041,028 | G | A | rs1367117 |
| TC | 2 | 21,070,463 | T | TAG | rs10692845 |
| TC | 2 | 27,518,370 | T | C | rs780094 |
| TC | 2 | 43,847,292 | C | T | rs4245791 |
| TC | 3 | 32,496,755 | C | T | rs3773777 |
| TC | 4 | 3,471,412 | A | G | rs6831256 |
| TC | 5 | 75,329,662 | C | A | rs7703051 |
| TC | 5 | 156,971,158 | G | C | rs1501908 |
| TC | 6 | 32,622,958 | C | T | rs35062987 |
| TC | 6 | 160,576,086 | A | T | rs74617384 |
| TC | 7 | 44,566,618 | C | G | rs217381 |
| TC | 8 | 9,326,086 | A | G | rs4841132 |
| TC | 8 | 18,415,371 | G | A | rs1495741 |
| TC | 8 | 58,479,765 | G | A | rs9297994 |
| TC | 8 | 125,467,120 | C | T | rs6982502 |
| TC | 9 | 104,826,853 | A | T | rs4149310 |
| TC | 9 | 104,886,314 | A | G | rs3847302 |
| TC | 9 | 104,903,697 | C | G | rs1800978 |
| TC | 9 | 133,279,427 | T | C | rs635634 |
| TC | 10 | 45,517,829 | A | C | rs970548 |
| TC | 11 | 61,803,311 | T | C | rs174547 |
| TC | 11 | 116,715,567 | T | C | rs7350481 |
| TC | 11 | 116,791,691 | G | C | rs3135506 |
| TC | 12 | 120,978,847 | A | C | rs1169288 |
| TC | 13 | 41,034,911 | G | A | rs17532301 |
| TC | 14 | 24,413,852 | C | T | rs2332328 |

| | | | | | |
|---|---|---|---|---|---|
| TC | 14 | 94,380,925 | T | A | rs17580 |
| TC | 15 | 58,387,979 | T | C | rs261291 |
| TC | 15 | 58,431,280 | T | C | rs1077834 |
| TC | 16 | 56,963,321 | G | A | rs1864163 |
| TC | 16 | 67,983,453 | A | G | rs255054 |
| TC | 16 | 72,018,449 | A | G | rs11648003 |
| TC | 16 | 72,054,562 | A | C | rs5471 |
| TC | 17 | 7,176,750 | A | G | rs12945299 |
| TC | 18 | 49,592,028 | T | C | rs9958734 |
| TC | 19 | 11,086,922 | G | T | rs73015024 |
| TC | 19 | 11,091,630 | G | T | rs6511720 |
| TC | 19 | 11,116,926 | C | T | rs688 |
| TC | 19 | 11,239,812 | C | T | rs2278426 |
| TC | 19 | 19,349,732 | G | C | rs73001065 |
| TC | 19 | 44,908,684 | T | C | rs429358 |
| TC | 19 | 44,908,822 | C | T | rs7412 |
| TC | 19 | 44,919,330 | A | G | rs12721054 |
| TC | 19 | 48,703,160 | A | G | rs492602 |
| TC | 20 | 40,551,182 | G | C | rs1883711 |
| TC | 20 | 41,043,978 | T | A | rs6029526 |
| TC | 21 | 31,687,518 | T | C | rs17660708 |
| TC | 22 | 43,928,975 | G | A | rs3747207 |

**Supplementary Table 5.** Gene-centric unconditional analysis results of lipid traits LDL-C, HDL-C, TG and TC in discovery phase using the TOPMed cohort (*n* = 21,015). Results for the significant genes (unconditional STAAR-O *P*-value < 3.57E-07 for 7 different noncoding masks across protein-coding genes; unconditional STAAR-O P-value < 2.50E-06 for ncRNA genes) are presented in the table. Four statistical tests were compared: Burden, SKAT, ACAT-V and STAAR-O. Chr (chromosome); Category (functional category); #SNV (number of rare variants (MAF < 1%) of the particular functional category in the gene); SKAT (SKAT *P*-value); Burden (Burden *P*-value); ACAT-V (ACAT-V *P*-value); STAAR-O (STAAR-O *P*-value); LDL-C (low-density lipoprotein cholesterol); HDL-C (high-density lipoprotein cholesterol); TG (triglycerides); TC (total cholesterol).

| Trait | Gene | Chr | Category | #SNV | SKAT | Burden | ACAT-V | STAAR-O |
|---|---|---|---|---|---|---|---|---|
| LDL-C | ILF3 | 19 | UTR | 523 | 5.60E-06 | 3.16E-04 | 2.36E-10 | 8.95E-10 |
| | TOMM40 | 19 | UTR | 227 | 8.89E-07 | 2.44E-02 | 8.43E-10 | 2.73E-09 |
| | LDLR | 19 | upstream | 68 | 1.61E-10 | 7.45E-03 | 7.90E-18 | 2.35E-17 |
| | SMARCA4 | 19 | downstream | 85 | 1.49E-11 | 1.36E-05 | 1.23E-12 | 2.96E-12 |
| | APOC1 | 19 | downstream | 92 | 2.56E-16 | 1.66E-06 | 8.26E-16 | 6.66E-16 |
| | QTRT1 | 19 | promoter_CAGE | 32 | 1.01E-09 | 5.53E-06 | 7.57E-10 | 1.28E-09 |
| | LDLR | 19 | promoter_CAGE | 131 | 4.39E-13 | 2.03E-06 | 6.57E-18 | 1.88E-17 |
| | APOE | 19 | promoter_CAGE | 91 | 8.33E-12 | 8.25E-02 | 9.65E-11 | 1.45E-11 |
| | QTRT1 | 19 | promoter_DHS | 149 | 6.21E-07 | 2.87E-03 | 1.53E-08 | 1.87E-08 |
| | LDLR | 19 | promoter_DHS | 257 | 4.05E-12 | 2.83E-04 | 1.46E-17 | 4.03E-17 |
| | APOE | 19 | promoter_DHS | 162 | 5.50E-11 | 6.23E-06 | 8.59E-11 | 9.81E-11 |
| | QTRT1 | 19 | enhancer_CAGE | 32 | 1.01E-09 | 5.53E-06 | 7.57E-10 | 1.28E-09 |
| | LDLR | 19 | enhancer_CAGE | 150 | 1.18E-13 | 6.98E-06 | 1.06E-17 | 2.82E-17 |
| | SLC44A2 | 19 | enhancer_DHS | 1716 | 1.55E-05 | 3.59E-01 | 8.83E-10 | 3.04E-09 |
| | QTRT1 | 19 | enhancer_DHS | 149 | 6.36E-05 | 6.70E-01 | 9.70E-09 | 2.44E-08 |
| | SMARCA4 | 19 | enhancer_DHS | 1746 | 1.19E-03 | 7.05E-01 | 6.32E-10 | 1.75E-09 |
| | LDLR | 19 | enhancer_DHS | 560 | 6.40E-13 | 3.26E-04 | 4.17E-17 | 9.75E-17 |
| | APOE | 19 | enhancer_DHS | 239 | 6.29E-09 | 1.72E-01 | 4.12E-10 | 9.84E-11 |
| | CTC-527H23.4 | 16 | ncRNA | 32 | 4.28E-03 | 5.53E-02 | 5.92E-07 | 1.15E-06 |
| | CTB-129P6.11 | 19 | ncRNA | 21 | 2.35E-05 | 1.99E-06 | 2.91E-06 | 2.14E-06 |
| HDL-C | APOA1 | 11 | enhancer_DHS | 1862 | 3.64E-03 | 9.39E-01 | 6.38E-08 | 2.19E-07 |
| | NLRC5 | 16 | enhancer_DHS | 871 | 1.12E-08 | 7.42E-02 | 1.99E-04 | 4.41E-08 |
| TG | APOE | 19 | promoter_CAGE | 92 | 1.01E-12 | 5.45E-02 | 4.11E-08 | 4.45E-12 |
| | APOA5 | 11 | promoter_DHS | 175 | 1.66E-08 | 1.72E-04 | 2.84E-06 | 2.39E-08 |
| | APOE | 19 | promoter_DHS | 163 | 2.72E-11 | 1.70E-01 | 5.42E-08 | 1.80E-11 |
| | APOC1 | 19 | promoter_DHS | 306 | 1.63E-07 | 5.73E-04 | 1.06E-07 | 9.81E-08 |
| | APOE | 19 | enhancer_CAGE | 53 | 1.17E-08 | 1.33E-07 | 3.16E-08 | 2.89E-08 |
| | COL18A1 | 21 | enhancer_CAGE | 256 | 1.81E-06 | 5.57E-04 | 1.28E-07 | 1.92E-07 |
| | APOE | 19 | enhancer_DHS | 241 | 4.72E-12 | 3.73E-01 | 9.29E-08 | 2.02E-11 |
| TC | ILF3 | 19 | UTR | 542 | 1.73E-06 | 3.15E-04 | 1.10E-09 | 4.19E-09 |
| | LDLR | 19 | upstream | 69 | 1.79E-10 | 1.78E-02 | 1.84E-16 | 5.55E-16 |
| | SMARCA4 | 19 | downstream | 89 | 1.24E-11 | 1.40E-04 | 3.67E-12 | 7.65E-12 |
| | APOC1 | 19 | downstream | 95 | 8.21E-11 | 8.24E-03 | 3.02E-10 | 2.54E-10 |
| | QTRT1 | 19 | promoter_CAGE | 33 | 1.66E-09 | 1.33E-05 | 1.48E-09 | 2.31E-09 |
| | LDLR | 19 | promoter_CAGE | 134 | 1.69E-13 | 1.03E-07 | 1.53E-16 | 4.44E-16 |
| | QTRT1 | 19 | promoter_DHS | 155 | 4.46E-07 | 1.64E-02 | 2.71E-08 | 3.32E-08 |
| | LDLR | 19 | promoter_DHS | 267 | 5.70E-13 | 2.21E-04 | 3.40E-16 | 1.11E-15 |
| | QTRT1 | 19 | enhancer_CAGE | 33 | 1.66E-09 | 1.33E-05 | 1.48E-09 | 2.31E-09 |
| | LDLR | 19 | enhancer_CAGE | 154 | 9.84E-14 | 1.01E-06 | 2.45E-16 | 6.66E-16 |
| | SLC44A2 | 19 | enhancer_DHS | 1754 | 1.41E-05 | 3.35E-01 | 4.03E-09 | 1.38E-08 |
| | QTRT1 | 19 | enhancer_DHS | 153 | 2.66E-05 | 1.72E-01 | 1.76E-08 | 4.25E-08 |
| | SMARCA4 | 19 | enhancer_DHS | 1781 | 1.67E-03 | 8.00E-01 | 2.06E-10 | 5.72E-10 |
| | LDLR | 19 | enhancer_DHS | 573 | 2.75E-11 | 6.82E-04 | 9.74E-16 | 2.44E-15 |

**Supplementary Table 6**. Known variants list in conditional analysis. The list is generated using the stepwise algorithm decribed in the manuscript and the common, low-frequent and rare phenotype-specific variants (MAC ≥ 20) indexed in GWAS Catalog.

| Trait | CHR | POS | REF | ALT | #rs |
|---|---|---|---|---|---|
| HDL | 1 | 109,274,968 | G | T | rs12740374 |
| HDL | 1 | 230,162,032 | A | T | rs2281718 |
| HDL | 2 | 21,008,652 | G | A | rs676210 |
| HDL | 2 | 164,694,691 | T | C | rs7607980 |
| HDL | 3 | 36,919,169 | C | A | rs7622114 |
| HDL | 4 | 109,717,608 | A | G | rs41278045 |
| HDL | 5 | 73,630,689 | A | G | rs6881956 |
| HDL | 6 | 43,792,590 | C | G | rs11967262 |
| HDL | 7 | 80,671,133 | T | G | rs3211938 |
| HDL | 7 | 130,747,722 | T | C | rs6971365 |
| HDL | 8 | 9,326,086 | A | G | rs4841132 |
| HDL | 8 | 19,961,928 | A | G | rs326 |
| HDL | 8 | 19,973,410 | C | T | rs10096633 |
| HDL | 8 | 20,070,502 | T | A | rs6999158 |
| HDL | 8 | 120,856,311 | G | T | rs4871137 |
| HDL | 8 | 125,495,147 | C | A | rs2954038 |
| HDL | 9 | 15,304,784 | C | A | rs686030 |
| HDL | 9 | 104,826,853 | A | T | rs4149310 |
| HDL | 9 | 104,858,554 | G | A | rs9282541 |
| HDL | 9 | 104,884,738 | C | T | rs11789603 |
| HDL | 9 | 104,902,020 | C | T | rs1883025 |
| HDL | 10 | 45,523,383 | A | G | rs11239549 |
| HDL | 11 | 61,824,890 | A | G | rs174566 |
| HDL | 11 | 116,778,201 | G | C | rs964184 |
| HDL | 11 | 116,830,638 | G | A | rs138326449 |
| HDL | 11 | 117,103,213 | G | C | rs12269901 |
| HDL | 11 | 117,218,489 | C | T | rs142953140 |
| HDL | 12 | 57,398,797 | C | T | rs11613352 |
| HDL | 12 | 123,975,620 | G | T | rs4765127 |
| HDL | 12 | 124,853,983 | C | T | rs10773112 |
| HDL | 13 | 44,950,491 | G | A | rs9526023 |
| HDL | 14 | 104,806,341 | A | T | rs45490496 |
| HDL | 15 | 43,528,519 | C | T | rs55707100 |
| HDL | 15 | 58,289,171 | T | A | rs12148399 |
| HDL | 15 | 58,391,167 | A | G | rs1532085 |
| HDL | 15 | 58,431,476 | C | T | rs1800588 |
| HDL | 16 | 56,955,678 | C | T | rs247616 |
| HDL | 16 | 56,961,915 | G | A | rs17231520 |
| HDL | 16 | 56,972,678 | C | T | rs7499892 |
| HDL | 16 | 56,973,441 | C | T | rs5883 |

| | | | | | |
|---|---|---|---|---|---|
| HDL | 16 | 56,981,179 | G | C | rs5880 |
| HDL | 16 | 67,943,479 | T | C | rs1109166 |
| HDL | 17 | 43,848,758 | C | T | rs72836561 |
| HDL | 18 | 49,592,028 | T | C | rs9958734 |
| HDL | 18 | 49,645,518 | C | G | rs8086351 |
| HDL | 19 | 8,364,439 | G | A | rs116843064 |
| HDL | 19 | 11,235,874 | C | T | rs3760782 |
| HDL | 19 | 11,240,198 | C | T | rs145464906 |
| HDL | 19 | 44,908,684 | T | C | rs429358 |
| HDL | 19 | 44,909,976 | G | T | rs1065853 |
| HDL | 19 | 44,945,208 | T | G | rs5167 |
| HDL | 20 | 44,413,724 | C | T | rs1800961 |
| HDL | 20 | 45,923,216 | T | C | rs6073958 |
| HDL | 21 | 42,298,682 | A | G | rs3746915 |
| HDL | 22 | 21,622,645 | T | C | rs7444 |
| LDL | 1 | 55,021,673 | C | G | rs12117661 |
| LDL | 1 | 55,039,974 | G | T | rs11591147 |
| LDL | 1 | 55,046,549 | C | G | rs67608943 |
| LDL | 1 | 55,052,749 | C | T | rs72646508 |
| LDL | 1 | 55,055,640 | G | T | rs472495 |
| LDL | 1 | 55,058,129 | A | G | rs28362261 |
| LDL | 1 | 55,058,182 | G | A | rs28362263 |
| LDL | 1 | 55,063,514 | G | A | rs505151 |
| LDL | 1 | 55,063,542 | C | A | rs28362286 |
| LDL | 1 | 109,274,968 | G | T | rs12740374 |
| LDL | 1 | 234,717,312 | C | T | rs556107 |
| LDL | 2 | 21,011,100 | T | C | rs533617 |
| LDL | 2 | 21,041,028 | G | A | rs1367117 |
| LDL | 2 | 21,065,354 | G | A | rs563290 |
| LDL | 2 | 43,847,292 | C | T | rs4245791 |
| LDL | 3 | 32,496,755 | C | T | rs3773777 |
| LDL | 4 | 68,475,769 | T | C | rs976058 |
| LDL | 5 | 75,355,259 | A | G | rs3846662 |
| LDL | 5 | 156,971,158 | G | C | rs1501908 |
| LDL | 6 | 160,589,086 | A | G | rs10455872 |
| LDL | 6 | 160,690,668 | C | T | rs186696265 |
| LDL | 7 | 44,566,618 | C | G | rs217381 |
| LDL | 8 | 9,323,885 | A | G | rs2169387 |
| LDL | 8 | 22,069,949 | C | G | rs7386762 |
| LDL | 8 | 125,466,208 | T | C | rs2001846 |
| LDL | 9 | 133,279,294 | T | G | rs495828 |
| LDL | 10 | 112,190,660 | T | C | rs7096937 |
| LDL | 11 | 61,803,910 | G | A | rs174549 |
| LDL | 11 | 116,778,201 | G | C | rs964184 |
| LDL | 12 | 120,951,159 | A | C | rs2650000 |

| | | | | | |
|---|---|---|---|---|---|
| LDL | 13 | 113,844,399 | C | T | rs6602911 |
| LDL | 14 | 24,413,852 | C | T | rs2332328 |
| LDL | 14 | 94,380,925 | T | A | rs17580 |
| LDL | 15 | 63,500,058 | T | C | rs56369308 |
| LDL | 16 | 72,063,928 | C | T | rs3794695 |
| LDL | 16 | 72,182,890 | C | A | rs7200153 |
| LDL | 17 | 66,214,462 | A | C | rs1801689 |
| LDL | 18 | 49,583,585 | A | G | rs77960347 |
| LDL | 19 | 11,086,585 | G | A | rs12151108 |
| LDL | 19 | 11,089,332 | C | T | rs17249141 |
| LDL | 19 | 11,116,926 | C | T | rs688 |
| LDL | 19 | 19,349,732 | G | C | rs73001065 |
| LDL | 19 | 44,908,684 | T | C | rs429358 |
| LDL | 19 | 44,908,783 | C | T | rs769455 |
| LDL | 19 | 44,908,822 | C | T | rs7412 |
| LDL | 19 | 44,935,906 | C | G | rs35136575 |
| LDL | 19 | 48,703,160 | A | G | rs492602 |
| LDL | 20 | 41,095,698 | T | C | rs6065311 |
| LDL | 21 | 15,214,362 | T | A | rs12106385 |
| LDL | 22 | 46,231,706 | C | T | rs4253772 |
| TG | 1 | 62,612,551 | A | T | rs10889348 |
| TG | 1 | 230,161,390 | C | T | rs2281721 |
| TG | 2 | 21,002,409 | C | T | rs1042034 |
| TG | 2 | 27,508,073 | T | C | rs1260326 |
| TG | 2 | 164,694,691 | T | C | rs7607980 |
| TG | 3 | 52,482,277 | C | G | rs6800707 |
| TG | 4 | 87,136,201 | G | A | rs1408 |
| TG | 5 | 131,337,265 | G | A | rs193735 |
| TG | 5 | 156,963,286 | T | C | rs6882076 |
| TG | 6 | 43,790,159 | C | A | rs998584 |
| TG | 6 | 139,518,361 | G | C | rs608736 |
| TG | 7 | 73,602,532 | T | C | rs13240994 |
| TG | 8 | 9,326,086 | A | G | rs4841132 |
| TG | 8 | 19,961,928 | A | G | rs326 |
| TG | 8 | 19,965,681 | T | C | rs3289 |
| TG | 8 | 20,070,649 | G | A | rs10106652 |
| TG | 8 | 125,495,066 | T | C | rs2980888 |
| TG | 9 | 84,002,350 | A | G | rs1982151 |
| TG | 10 | 50,814,012 | C | T | rs41274050 |
| TG | 11 | 61,830,500 | A | G | rs1535 |
| TG | 11 | 116,778,201 | G | C | rs964184 |
| TG | 11 | 116,779,090 | G | A | rs75198898 |
| TG | 11 | 116,789,970 | G | A | rs2266788 |
| TG | 11 | 116,791,691 | G | C | rs3135506 |
| TG | 11 | 116,794,363 | C | T | rs9804646 |

| | | | | | |
|---|---|---|---|---|---|
| TG | 12 | 21,178,615 | T | C | rs4149056 |
| TG | 13 | 113,844,399 | C | T | rs6602911 |
| TG | 14 | 103,824,476 | A | G | rs12893623 |
| TG | 15 | 42,429,647 | G | A | rs184334219 |
| TG | 15 | 43,735,687 | T | C | rs139974673 |
| TG | 15 | 58,387,979 | T | C | rs261291 |
| TG | 16 | 56,956,804 | C | A | rs247617 |
| TG | 16 | 81,501,185 | T | C | rs2925979 |
| TG | 17 | 43,763,481 | C | T | rs77697917 |
| TG | 18 | 289,209 | C | T | rs6506033 |
| TG | 19 | 8,364,439 | G | A | rs116843064 |
| TG | 19 | 19,285,807 | C | T | rs188247550 |
| TG | 19 | 19,321,481 | AG | A | rs140868651 |
| TG | 19 | 44,908,684 | T | C | rs429358 |
| TG | 19 | 44,908,783 | C | T | rs769455 |
| TG | 19 | 44,908,822 | C | T | rs7412 |
| TG | 19 | 44,919,330 | A | G | rs12721054 |
| TG | 19 | 44,927,023 | C | G | rs5112 |
| TG | 19 | 48,756,272 | A | G | rs838133 |
| TG | 20 | 41,152,292 | G | A | rs6093446 |
| TG | 20 | 45,923,216 | T | C | rs6073958 |
| TG | 21 | 45,455,861 | G | A | rs114139997 |
| TG | 22 | 38,204,535 | C | T | rs2267373 |
| TC | 1 | 23,421,503 | GA | G | rs11340914 |
| TC | 1 | 55,021,673 | C | G | rs12117661 |
| TC | 1 | 55,039,974 | G | T | rs11591147 |
| TC | 1 | 55,046,549 | C | G | rs67608943 |
| TC | 1 | 55,052,794 | A | G | rs2495477 |
| TC | 1 | 55,058,182 | G | A | rs28362263 |
| TC | 1 | 55,063,514 | G | A | rs505151 |
| TC | 1 | 55,063,542 | C | A | rs28362286 |
| TC | 1 | 62,620,326 | T | A | rs12239736 |
| TC | 1 | 109,274,968 | G | T | rs12740374 |
| TC | 1 | 234,722,850 | A | T | rs514230 |
| TC | 2 | 21,011,100 | T | C | rs533617 |
| TC | 2 | 21,041,028 | G | A | rs1367117 |
| TC | 2 | 21,070,463 | T | TAG | rs10692845 |
| TC | 2 | 27,518,370 | T | C | rs780094 |
| TC | 2 | 43,847,292 | C | T | rs4245791 |
| TC | 3 | 32,496,755 | C | T | rs3773777 |
| TC | 4 | 3,471,412 | A | G | rs6831256 |
| TC | 5 | 75,329,662 | C | A | rs7703051 |
| TC | 5 | 156,971,158 | G | C | rs1501908 |
| TC | 6 | 32,622,958 | C | T | rs35062987 |
| TC | 6 | 160,154,334 | G | A | rs11753995 |

| | | | | | |
|---|---|---|---|---|---|
| TC | 6 | 160,576,086 | A | T | rs74617384 |
| TC | 6 | 160,690,668 | C | T | rs186696265 |
| TC | 7 | 44,566,618 | C | G | rs217381 |
| TC | 8 | 9,326,086 | A | G | rs4841132 |
| TC | 8 | 18,415,371 | G | A | rs1495741 |
| TC | 8 | 58,479,765 | G | A | rs9297994 |
| TC | 8 | 125,467,120 | C | T | rs6982502 |
| TC | 9 | 104,826,853 | A | T | rs4149310 |
| TC | 9 | 104,886,314 | A | G | rs3847302 |
| TC | 9 | 104,903,697 | C | G | rs1800978 |
| TC | 9 | 133,279,427 | T | C | rs635634 |
| TC | 10 | 45,517,829 | A | C | rs970548 |
| TC | 11 | 61,803,311 | T | C | rs174547 |
| TC | 11 | 116,715,567 | T | C | rs7350481 |
| TC | 11 | 116,791,691 | G | C | rs3135506 |
| TC | 12 | 120,978,847 | A | C | rs1169288 |
| TC | 13 | 41,034,911 | G | A | rs17532301 |
| TC | 14 | 24,413,852 | C | T | rs2332328 |
| TC | 14 | 94,380,925 | T | A | rs17580 |
| TC | 15 | 58,387,979 | T | C | rs261291 |
| TC | 15 | 58,431,280 | T | C | rs1077834 |
| TC | 16 | 56,963,321 | G | A | rs1864163 |
| TC | 16 | 67,983,453 | A | G | rs255054 |
| TC | 16 | 72,018,449 | A | G | rs11648003 |
| TC | 16 | 72,054,562 | A | C | rs5471 |
| TC | 17 | 7,176,750 | A | G | rs12945299 |
| TC | 18 | 49,583,585 | A | G | rs77960347 |
| TC | 18 | 49,592,028 | T | C | rs9958734 |
| TC | 19 | 11,086,922 | G | T | rs73015024 |
| TC | 19 | 11,089,332 | C | T | rs17249141 |
| TC | 19 | 11,116,926 | C | T | rs688 |
| TC | 19 | 11,239,812 | C | T | rs2278426 |
| TC | 19 | 19,349,732 | G | C | rs73001065 |
| TC | 19 | 44,908,684 | T | C | rs429358 |
| TC | 19 | 44,908,822 | C | T | rs7412 |
| TC | 19 | 44,919,330 | A | G | rs12721054 |
| TC | 19 | 48,703,160 | A | G | rs492602 |
| TC | 20 | 40,551,182 | G | C | rs1883711 |
| TC | 20 | 41,043,978 | T | A | rs6029526 |
| TC | 21 | 31,687,518 | T | C | rs17660708 |
| TC | 22 | 43,928,975 | G | A | rs3747207 |

**Supplementary Table 7**. Gene-centric analysis results from unconditional analysis and analysis conditional on selected common, low-frequency, and rare variants. 21,015 discovery samples and 9,123 replication samples from the NHLBI Trans-Omics for Precision Medicine (TOPMed) program are considered. The 4 replicated conditionally significant genes (adjust for known common and low-frequent variants) are presented. Chr (Chromosome); Category (Functional category); #SNV (Number of rare variants (MAF < 1%) of the particular functional category in the gene); STAAR-O (STAAR-O P-value); LDL-C (Low-density lipoprotein cholesterol); HDL-C (High-density lipoprotein cholesterol); TG (Triglycerides); TC (Total cholesterol); Variants Adjusted (Adjusted variants in conditional analysis).

| Trait | Gene | Chr | Category | Discovery | | | Replication | | | Variants Adjusted |
|---|---|---|---|---|---|---|---|---|---|---|
| | | | | #SNV | STAAR-O (Unconditional) | STAAR-O (Conditional) | #SNV | STAAR-O (Unconditional) | STAAR-O (Conditional) | |
| HDL-C | *APOA1* | 11 | enhancer_DHS | 1862 | 2.19E-07 | 4.02E-02 | 1005 | 1.50E-03 | 2.03E-01 | rs964184,rs138326449,rs12269901,rs142953140 |
| TG | *APOE* | 19 | promoter_CAGE | 92 | 4.45E-12 | 5.34E-01 | 36 | 9.45E-06 | 9.44E-01 | rs429358, rs769455, rs7412, rs12721054, rs5112 |
| TG | *APOE* | 19 | promoter_DHS | 163 | 1.80E-11 | 9.56E-06 | 65 | 2.96E-06 | 5.22E-01 | rs429358, rs769455, rs7412, rs12721054, rs5112 |
| TG | *APOE* | 19 | enhancer_DHS | 241 | 2.02E-11 | 7.53E-01 | 116 | 1.12E-05 | 9.37E-01 | rs429358, rs769455, rs7412, rs12721054, rs5112 |

**Supplementary Table 8.** 2-kb sliding window analysis results of both unconditional analysis and anlaysis conditional on known common and low-frequency variants. 21,015 discovery samples and 9,123 replication samples from the NHLBI Trans-Omics for Precision Medicine (TOPMed) program were considered in the analysis. Results for the significant sliding windows (unconditional STAAR-O *P*-value < 1.88E-08) using discovery samples are presented. Four statistical tests were compared: Burden, SKAT, ACAT-V and STAAR-O. Chr (chromosome); Start Location (start location of the 2-kb sliding window); End Location (end location of the 2-kb sliding window); #SNV (number of rare variants (MAF < 1%) in the 2-kb sliding window); SKAT (SKAT *P*-value); Burden (Burden *P*-value); ACAT-V (ACAT-V *P*-value); STAAR-O (STAAR-O *P*-value); LDL-C (low-density lipoprotein cholesterol); HDL-C (High-density lipoprotein cholesterol); TG (triglycerides); TC (total cholesterol); Variants Adjusted (adjusted variants in conditional analysis). Physical positions of each window are on build hg38.

| Trait | Chr | Start Location | End Location | Gene | Discovery | | | | | | Replication | | | Variants Adjusted |
|---|---|---|---|---|---|---|---|---|---|---|---|---|---|---|
| | | | | | #SNV | SKAT | Burden | ACAT-V | STAAR-O (Unconditional) | STAAR-O (Conditional) | #SNV | STAAR-O (Unconditional) | STAAR-O (Conditional) | |
| | 1 | 55,045,498 | 55,047,497 | *PCSK9* | 160 | 1.20E-08 | 1.14E-03 | 3.91E-12 | 8.62E-12 | 9.72E-02 | 77 | 6.76E-03 | 9.72E-02 | rs11591147, rs28362263, rs505151, rs12117661, rs472495 |
| | 1 | 55,046,498 | 55,048,497 | *PCSK9* | 166 | 1.74E-08 | 2.42E-07 | 2.75E-12 | 6.37E-12 | 2.73E-02 | 81 | 3.28E-03 | 2.73E-02 | rs11591147, rs28362263, rs505151, rs12117661, rs472495 |
| | 1 | 55,057,498 | 55,059,497 | *PCSK9* | 158 | 8.01E-06 | 6.22E-01 | 1.04E-06 | 5.06E-10 | 3.84E-02 | 80 | 1.69E-02 | 3.84E-02 | rs11591147, rs28362263, rs505151, rs12117661, rs472495 |
| | 1 | 55,062,498 | 55,064,497 | *PCSK9* | 182 | 6.29E-12 | 2.91E-02 | 1.38E-32 | 2.43E-32 | 1.72E-12 | 102 | 5.16E-11 | 1.72E-12 | rs11591147, rs28362263, rs505151, rs12117661, rs472495 |
| | 1 | 55,063,498 | 55,065,497 | *PCSK9* | 163 | 4.86E-15 | 1.45E-01 | 9.28E-33 | 2.28E-32 | 1.40E-12 | 84 | 4.46E-11 | 1.40E-12 | rs11591147, rs28362263, rs505151, rs12117661, rs472495 |
| | 1 | 55,291,498 | 55,293,497 | *Intergenic (PCSK9)* | 164 | 9.15E-09 | 2.39E-01 | 7.14E-05 | 1.81E-08 | 2.89E-01 | 77 | 2.17E-02 | 2.89E-01 | rs11591147, rs28362263, rs505151, rs12117661, rs472495 |
| | 1 | 55,333,498 | 55,335,497 | *Intergenic (GOT2P1)* | 171 | 3.67E-05 | 2.16E-02 | 1.30E-16 | 6.66E-16 | 5.81E-07 | 95 | 1.27E-06 | 5.81E-07 | rs11591147, rs28362263, rs505151, rs12117661, rs472495 |
| | 1 | 55,334,498 | 55,336,497 | *Intergenic (GOT2P1)* | 148 | 1.85E-04 | 6.87E-02 | 1.35E-16 | 5.55E-16 | 5.49E-07 | 81 | 1.20E-06 | 5.49E-07 | rs11591147, rs28362263, rs505151, rs12117661, rs472495 |
| | 19 | 10,689,528 | 10,691,527 | *ILF3* | 205 | 7.72E-06 | 1.07E-04 | 1.28E-10 | 4.66E-10 | 8.30E-01 | 113 | 4.85E-01 | 8.30E-01 | rs12151108, rs688, rs6511720 |
| | 19 | 10,690,528 | 10,692,527 | *ILF3* | 243 | 1.45E-06 | 1.07E-04 | 1.05E-10 | 3.78E-10 | 4.96E-01 | 112 | 2.80E-01 | 4.96E-01 | rs12151108, rs688, rs6511720 |
| | 19 | 10,701,528 | 10,703,527 | *QTRT1* | 119 | 1.17E-04 | 4.67E-02 | 4.94E-09 | 8.15E-09 | 5.39E-01 | 61 | 4.00E-01 | 5.39E-01 | rs12151108, rs688, rs6511720 |
| | 19 | 10,938,528 | 10,940,527 | *Intergenic (SMARCA4)* | 143 | 3.78E-05 | 3.39E-01 | 6.86E-11 | 1.85E-10 | 1.63E-01 | 72 | 9.29E-02 | 1.63E-01 | rs12151108, rs688, rs6511720 |
| | 19 | 10,939,528 | 10,941,527 | *Intergenic (SMARCA4)* | 148 | 1.43E-05 | 7.96E-01 | 8.69E-11 | 2.26E-10 | 1.64E-01 | 88 | 9.08E-02 | 1.64E-01 | rs12151108, rs688, rs6511720 |
| | 19 | 10,987,528 | 10,989,527 | *SMARCA4* | 178 | 1.80E-05 | 5.43E-01 | 5.64E-11 | 1.59E-10 | 2.07E-02 | 88 | 1.06E-02 | 2.07E-02 | rs12151108, rs688, rs6511720 |
| | 19 | 10,988,528 | 10,990,527 | *SMARCA4* | 158 | 3.53E-03 | 3.08E-01 | 6.40E-11 | 8.80E-11 | 1.58E-01 | 82 | 9.08E-02 | 1.58E-01 | rs12151108, rs688, rs6511720 |
| | 19 | 11,027,528 | 11,029,527 | *SMARCA4* | 176 | 3.23E-09 | 1.04E-01 | 3.30E-09 | 3.21E-09 | 4.79E-01 | 76 | 3.16E-01 | 4.79E-01 | rs12151108, rs688, rs6511720 |
| | 19 | 11,028,528 | 11,030,527 | *SMARCA4* | 182 | 1.19E-06 | 9.79E-01 | 4.28E-09 | 4.99E-09 | 1.15E-01 | 84 | 1.32E-01 | 1.15E-01 | rs12151108, rs688, rs6511720 |
| | 19 | 11,040,528 | 11,042,527 | *SMARCA4* | 170 | 7.21E-09 | 4.43E-01 | 2.08E-09 | 3.36E-09 | 7.62E-02 | 72 | 6.46E-02 | 7.62E-02 | rs12151108, rs688, rs6511720 |
| | 19 | 11,041,528 | 11,043,527 | *SMARCA4* | 160 | 1.52E-06 | 6.86E-01 | 1.91E-09 | 4.29E-09 | 3.38E-01 | 83 | 1.77E-01 | 3.38E-01 | rs12151108, rs688, rs6511720 |
| | 19 | 11,042,528 | 11,044,527 | *SMARCA4* | 145 | 1.52E-12 | 2.87E-01 | 1.66E-15 | 1.11E-15 | 7.13E-01 | 80 | 4.44E-01 | 7.13E-01 | rs12151108, rs688, rs6511720 |
| | 19 | 11,043,528 | 11,045,527 | *SMARCA4* | 186 | 8.71E-13 | 4.61E-01 | 2.01E-15 | 1.33E-15 | 8.16E-01 | 83 | 4.50E-01 | 8.16E-01 | rs12151108, rs688, rs6511720 |
| | 19 | 11,056,528 | 11,058,527 | *SMARCA4* | 175 | 2.12E-07 | 1.92E-02 | 4.48E-11 | 1.41E-10 | 1.93E-02 | 102 | 1.25E-01 | 1.93E-02 | rs12151108, rs688, rs6511720 |
| | 19 | 11,057,528 | 11,059,527 | *SMARCA4* | 178 | 4.85E-08 | 4.14E-01 | 3.70E-11 | 1.37E-10 | 8.48E-03 | 93 | 4.31E-02 | 8.48E-03 | rs12151108, rs688, rs6511720 |
| | 19 | 11,060,528 | 11,062,527 | *SMARCA4* | 195 | 3.49E-06 | 7.14E-01 | 2.59E-12 | 8.27E-12 | 3.44E-01 | 101 | 2.74E-01 | 3.44E-01 | rs12151108, rs688, rs6511720 |
| | 19 | 11,061,528 | 11,063,527 | *SMARCA4* | 190 | 4.77E-12 | 9.01E-05 | 2.14E-12 | 5.25E-12 | 1.61E-01 | 105 | 4.66E-02 | 1.61E-01 | rs12151108, rs688, rs6511720 |
| | 19 | 11,062,528 | 11,064,527 | *SMARCA4* | 160 | 2.15E-06 | 4.56E-02 | 1.27E-11 | 3.77E-11 | 8.43E-01 | 89 | 2.81E-01 | 8.43E-01 | rs12151108, rs688, rs6511720 |
| | 19 | 11,077,528 | 11,079,527 | *Intergenic (LDLR)* | 153 | 3.85E-07 | 8.02E-01 | 3.05E-07 | 4.86E-12 | 7.99E-01 | 82 | 9.56E-03 | 7.99E-01 | rs12151108, rs688, rs6511720 |
| | 19 | 11,081,528 | 11,083,527 | *Intergenic (LDLR)* | 152 | 3.04E-09 | 4.15E-01 | 3.66E-07 | 9.47E-09 | 6.41E-01 | 90 | 3.70E-02 | 6.41E-01 | rs12151108, rs688, rs6511720 |
| | 19 | 11,084,528 | 11,086,527 | *Intergenic (LDLR)* | 143 | 7.47E-13 | 6.92E-02 | 1.23E-15 | 3.33E-15 | 4.94E-01 | 73 | 3.06E-01 | 4.94E-01 | rs12151108, rs688, rs6511720 |
| LDL-C | 19 | 11,085,528 | 11,087,527 | *Intergenic (LDLR)* | 137 | 7.32E-07 | 1.30E-01 | 1.51E-15 | 3.89E-15 | 2.45E-01 | 73 | 1.09E-01 | 2.45E-01 | rs12151108, rs688, rs6511720 |
| | 19 | 11,087,528 | 11,089,527 | *LDLR* | 137 | 3.99E-07 | 2.07E-01 | 1.33E-17 | 3.38E-17 | 5.66E-01 | 59 | 3.14E-01 | 5.66E-01 | rs12151108, rs688, rs6511720 |
| | 19 | 11,088,528 | 11,090,527 | *LDLR* | 157 | 2.20E-06 | 3.84E-01 | 1.51E-17 | 4.46E-17 | 3.08E-01 | 77 | 6.35E-01 | 3.08E-01 | rs12151108, rs688, rs6511720 |
| | 19 | 11,097,528 | 11,099,527 | *LDLR* | 114 | 7.52E-11 | 6.24E-01 | 1.06E-16 | 4.44E-16 | 7.24E-01 | 68 | 9.81E-01 | 7.24E-01 | rs12151108, rs688, rs6511720 |
| | 19 | 11,098,528 | 11,100,527 | *LDLR* | 135 | 1.25E-10 | 2.09E-01 | 9.59E-17 | 4.44E-16 | 7.60E-01 | 76 | 7.66E-02 | 7.60E-01 | rs12151108, rs688, rs6511720 |
| | 19 | 11,118,528 | 11,120,527 | *LDLR* | 166 | 3.64E-15 | 1.56E-06 | 6.37E-11 | 6.37E-14 | 3.56E-04 | 96 | 1.30E-04 | 3.56E-04 | rs12151108, rs688, rs6511720 |
| | 19 | 11,119,528 | 11,121,527 | *LDLR* | 155 | 4.32E-13 | 1.19E-06 | 3.33E-11 | 1.37E-12 | 1.59E-04 | 79 | 1.10E-04 | 1.59E-04 | rs12151108, rs688, rs6511720 |
| | 19 | 11,169,528 | 11,171,527 | *KANK2* | 167 | 5.90E-05 | 6.91E-02 | 1.62E-11 | 6.21E-11 | 8.78E-01 | 85 | 9.26E-01 | 8.78E-01 | rs12151108, rs688, rs6511720 |
| | 19 | 11,170,528 | 11,172,527 | *KANK2* | 157 | 4.77E-07 | 1.55E-01 | 1.41E-11 | 4.59E-11 | 4.12E-01 | 77 | 9.44E-02 | 4.12E-01 | rs12151108, rs688, rs6511720 |
| | 19 | 44,827,528 | 44,829,527 | *Intergenic (NECTIN2)* | 140 | 6.22E-09 | 2.79E-04 | 1.36E-08 | 4.44E-09 | 7.14E-01 | 77 | 5.04E-01 | 7.14E-01 | rs7412, rs429358, rs35136575 |
| | 19 | 44,828,528 | 44,830,527 | *Intergenic (NECTIN2)* | 126 | 3.77E-09 | 8.55E-04 | 1.58E-08 | 1.62E-08 | 1.43E-02 | 71 | 1.79E-03 | 1.43E-02 | rs7412, rs429358, rs35136575 |
| | 19 | 44,857,528 | 44,859,527 | *NECTIN2* | 159 | 8.53E-07 | 2.12E-03 | 9.64E-07 | 1.00E-08 | 4.55E-02 | 66 | 6.37E-04 | 4.55E-02 | rs7412, rs429358, rs35136575 |
| | 19 | 44,882,528 | 44,884,527 | *NECTIN2* | 145 | 8.72E-09 | 8.12E-03 | 8.03E-09 | 6.38E-09 | 8.61E-02 | 85 | 2.93E-01 | 8.61E-02 | rs7412, rs429358, rs35136575 |
| | 19 | 44,890,528 | 44,892,527 | *TOMM40* | 202 | 1.06E-09 | 3.22E-03 | 7.43E-11 | 1.54E-10 | 3.05E-01 | 92 | 3.68E-02 | 3.05E-01 | rs7412, rs429358, rs35136575 |
| | 19 | 44,891,528 | 44,893,527 | *TOMM40* | 166 | 4.24E-16 | 3.64E-05 | 6.72E-11 | 2.09E-17 | 4.48E-01 | 77 | 6.18E-04 | 4.48E-01 | rs7412, rs429358, rs35136575 |
| | 19 | 44,892,528 | 44,894,527 | *TOMM40* | 169 | 3.68E-15 | 4.38E-03 | 2.71E-11 | 2.29E-14 | 7.61E-01 | 86 | 1.59E-05 | 7.61E-01 | rs7412, rs429358, rs35136575 |
| | 19 | 44,893,528 | 44,895,527 | *TOMM40* | 159 | 3.26E-24 | 2.55E-01 | 2.63E-11 | 1.86E-23 | 2.28E-01 | 79 | 1.47E-05 | 2.28E-01 | rs7412, rs429358, rs35136575 |
| | 19 | 44,894,528 | 44,896,527 | *Intron (TOMM40)* | 174 | 4.82E-16 | 6.33E-04 | 1.78E-10 | 9.97E-17 | 8.94E-02 | 91 | 7.17E-03 | 8.94E-02 | rs7412, rs429358, rs35136575 |
| | 19 | 44,897,528 | 44,899,527 | *Intron (TOMM40)* | 172 | 8.20E-08 | 5.54E-04 | 3.13E-11 | 1.11E-10 | 2.84E-01 | 104 | 1.31E-05 | 2.84E-01 | rs7412, rs429358, rs35136575 |
| | 19 | 44,898,528 | 44,900,527 | *Intron (TOMM40)* | 140 | 5.45E-09 | 8.00E-01 | 3.00E-11 | 1.10E-10 | 5.20E-01 | 78 | 1.71E-05 | 5.20E-01 | rs7412, rs429358, rs35136575 |
| | 19 | 44,900,528 | 44,902,527 | *TOMM40* | 170 | 2.85E-08 | 4.45E-01 | 6.61E-10 | 2.01E-09 | 4.63E-01 | 89 | 2.08E-01 | 4.63E-01 | rs7412, rs429358, rs35136575 |
| | 19 | 44,901,528 | 44,903,527 | *TOMM40* | 181 | 8.37E-07 | 1.26E-01 | 7.10E-10 | 1.62E-09 | 5.27E-01 | 110 | 1.15E-01 | 5.27E-01 | rs7412, rs429358, rs35136575 |
| | 19 | 44,905,528 | 44,907,527 | *APOE* | 148 | 3.79E-12 | 5.29E-02 | 1.30E-10 | 1.38E-11 | 4.09E-02 | 55 | 1.39E-01 | 4.09E-02 | rs7412, rs429358, rs35136575 |
| | 19 | 44,906,528 | 44,908,527 | *APOE* | 143 | 4.37E-11 | 9.96E-06 | 1.05E-10 | 1.03E-10 | 7.15E-02 | 57 | 1.27E-01 | 7.15E-02 | rs7412, rs429358, rs35136575 |
| | 19 | 44,907,528 | 44,909,527 | *APOE* | 175 | 3.95E-11 | 2.10E-04 | 9.69E-11 | 5.60E-11 | 5.73E-02 | 73 | 4.62E-02 | 5.73E-02 | rs7412, rs429358, rs35136575 |
| | 19 | 44,908,528 | 44,910,527 | *APOE* | 199 | 8.37E-12 | 1.74E-04 | 7.61E-11 | 2.74E-11 | 7.33E-02 | 81 | 4.46E-02 | 7.33E-02 | rs7412, rs429358, rs35136575 |
| | 19 | 44,918,528 | 44,920,527 | *APOC1* | 177 | 1.48E-13 | 1.15E-03 | 1.40E-15 | 1.33E-15 | 6.79E-02 | 75 | 1.86E-01 | 6.79E-02 | rs7412, rs429358, rs35136575 |
| | 19 | 44,919,528 | 44,921,527 | *Intergenic (APOC1)* | 180 | 1.21E-27 | 1.18E-15 | 2.10E-15 | 5.32E-27 | 6.87E-02 | 87 | 3.53E-03 | 6.87E-02 | rs7412, rs429358, rs35136575 |
| | 19 | 44,920,528 | 44,922,527 | *Intergenic (APOC1)* | 164 | 3.54E-16 | 7.87E-07 | 1.59E-14 | 2.22E-15 | 4.38E-01 | 88 | 9.74E-05 | 4.38E-01 | rs7412, rs429358, rs35136575 |
| | 19 | 44,930,528 | 44,932,527 | *APOC1P1* | 171 | 1.54E-09 | 2.33E-02 | 7.90E-10 | 5.46E-12 | 6.77E-04 | 104 | 4.47E-08 | 6.77E-04 | rs7412, rs429358, rs35136575 |
| | 19 | 44,931,528 | 44,933,527 | *Intergenic (APOC1P1)* | 148 | 1.03E-10 | 2.25E-01 | 7.34E-10 | 9.84E-13 | 9.98E-04 | 88 | 1.24E-07 | 9.98E-04 | rs7412, rs429358, rs35136575 |
| | 8 | 57,071,644 | 57,073,643 | *Intergenic (IMPAD1)* | 111 | 1.28E-05 | 2.76E-09 | 2.05E-03 | 1.79E-08 | 1.79E-08 | 53 | 8.38E-01 | 8.38E-01 | n/a |
| | 11 | 116,802,930 | 116,804,929 | *Intergenic (ZPR1)* | 135 | 7.81E-02 | 3.37E-02 | 4.15E-09 | 1.25E-08 | 4.31E-08 | 76 | 9.49E-05 | 2.02E-04 | rs964184, rs12269901 |
| | 11 | 116,828,930 | 116,830,929 | *APOC3* | 172 | 3.16E-06 | 1.82E-02 | 2.00E-10 | 4.44E-10 | 1.05E-08 | 103 | 1.57E-06 | 2.55E-06 | rs964184, rs12269901 |

| | | | | | | | | | | | | | | |
|---|---|---|---|---|---|---|---|---|---|---|---|---|---|---|
| | 11 | 116,829,930 | 116,831,929 | *APOC3* | 150 | 1.11E-05 | 1.65E-03 | 1.85E-10 | 3.62E-10 | 7.33E-09 | 97 | 5.99E-07 | 1.21E-06 | rs964184, rs12269901 |
| | 11 | 116,860,930 | 116,862,929 | *SIK3* | 109 | 4.09E-04 | 5.23E-02 | 4.58E-09 | 1.42E-08 | 4.86E-08 | 71 | 9.92E-05 | 2.11E-04 | rs964184, rs12269901 |
| | 11 | 117,146,930 | 117,148,929 | *Intron (PAFAH1B2)* | 165 | 9.64E-04 | 9.97E-02 | 1.55E-09 | 5.98E-09 | 8.28E-08 | 98 | 6.02E-04 | 1.12E-03 | rs964184, rs12269901 |
| HDL-C | 11 | 117,147,930 | 117,149,929 | *Intron (PAFAH1B2)* | 168 | 9.09E-03 | 5.31E-01 | 2.51E-09 | 8.85E-09 | 1.22E-07 | 96 | 8.72E-04 | 1.64E-03 | rs964184, rs12269901 |
| | 16 | 56,760,029 | 56,762,028 | *Intron (NUP93)* | 132 | 3.83E-05 | 5.29E-01 | 7.16E-09 | 1.38E-08 | 9.65E-06 | 68 | 2.45E-01 | 1.15E-01 | rs247616, rs5883, rs7499892, rs17231520, rs5880 |
| | 16 | 56,761,029 | 56,763,028 | *Intron (NUP93)* | 141 | 2.07E-06 | 1.53E-01 | 5.17E-09 | 1.50E-08 | 1.09E-05 | 73 | 5.87E-01 | 2.26E-01 | rs247616, rs5883, rs7499892, rs17231520, rs5880 |
| | 16 | 56,956,029 | 56,958,028 | *CETP* | 149 | 8.90E-11 | 8.79E-01 | 2.99E-09 | 3.22E-10 | 3.48E-01 | 86 | 2.48E-05 | 2.85E-01 | rs247616, rs5883, rs7499892, rs17231520, rs5880 |
| | 16 | 56,957,029 | 56,959,028 | *CETP* | 149 | 1.26E-08 | 4.92E-01 | 2.17E-09 | 5.46E-09 | 2.28E-01 | 80 | 1.93E-05 | 4.40E-01 | rs247616, rs5883, rs7499892, rs17231520, rs5880 |
| | 16 | 56,970,029 | 56,972,028 | *CETP* | 155 | 7.51E-12 | 5.80E-02 | 2.50E-12 | 3.55E-13 | 5.18E-09 | 89 | 1.48E-03 | 2.43E-01 | rs247616, rs5883, rs7499892, rs17231520, rs5880 |
| | 16 | 56,971,029 | 56,973,028 | *CETP* | 156 | 2.03E-12 | 1.34E-02 | 3.28E-12 | 2.93E-14 | 1.00E-08 | 83 | 7.49E-03 | 5.35E-02 | rs247616, rs5883, rs7499892, rs17231520, rs5880 |
| | 16 | 56,977,029 | 56,979,028 | *CETP* | 177 | 1.36E-10 | 5.02E-01 | 8.67E-05 | 2.36E-09 | 4.55E-05 | 102 | 1.30E-03 | 6.16E-03 | rs247616, rs5883, rs7499892, rs17231520, rs5880 |
| | 11 | 116,666,930 | 116,668,929 | *Intergenic (AP000770.1)* | 206 | 1.95E-06 | 8.80E-01 | 2.76E-06 | 6.07E-09 | 1.33E-02 | 137 | 8.44E-02 | 1.09E-01 | rs964184, rs9804646, rs3135506, rs2266788 |
| | 11 | 116,706,930 | 116,708,929 | *Intergenic (AP000770.1)* | 185 | 1.14E-13 | 5.30E-01 | 1.29E-09 | 1.16E-12 | 3.78E-02 | 94 | 5.93E-03 | 3.03E-01 | rs964184, rs9804646, rs3135506, rs2266788 |
| | 11 | 116,707,930 | 116,709,929 | *Intergenic (AP000770.1)* | 169 | 7.63E-09 | 9.35E-01 | 1.29E-09 | 9.66E-11 | 4.69E-02 | 80 | 1.70E-02 | 3.15E-01 | rs964184, rs9804646, rs3135506, rs2266788 |
| | 11 | 116,746,930 | 116,748,929 | *Intergenic (AP000770.1)* | 137 | 5.35E-09 | 1.19E-02 | 2.21E-09 | 5.37E-09 | 9.00E-01 | 58 | 3.50E-02 | 2.03E-01 | rs964184, rs9804646, rs3135506, rs2266788 |
| | 11 | 116,747,930 | 116,749,929 | *BUD13* | 157 | 6.03E-11 | 1.13E-05 | 3.34E-09 | 2.19E-10 | 1.08E-01 | 76 | 1.05E-02 | 5.55E-01 | rs964184, rs9804646, rs3135506, rs2266788 |
| | 11 | 116,777,930 | 116,779,929 | *ZPR1* | 135 | 6.20E-09 | 6.47E-04 | 5.95E-09 | 4.45E-10 | 1.11E-09 | 88 | 3.07E-06 | 7.62E-06 | rs964184, rs9804646, rs3135506, rs2266788 |
| | 11 | 116,778,930 | 116,780,929 | *ZPR1* | 121 | 1.09E-09 | 6.70E-05 | 5.91E-09 | 7.16E-11 | 2.39E-10 | 80 | 9.59E-07 | 2.85E-06 | rs964184, rs9804646, rs3135506, rs2266788 |
| TG | 11 | 116,828,930 | 116,830,929 | *APOC3* | 171 | 2.08E-13 | 2.37E-05 | 1.48E-25 | 3.64E-25 | 1.50E-23 | 100 | 6.23E-21 | 5.88E-20 | rs964184, rs9804646, rs3135506, rs2266788 |
| | 11 | 116,829,930 | 116,831,929 | *APOC3* | 151 | 7.27E-10 | 1.99E-07 | 1.64E-25 | 3.53E-25 | 1.45E-23 | 94 | 5.22E-20 | 7.31E-19 | rs964184, rs9804646, rs3135506, rs2266788 |
| | 11 | 117,146,930 | 117,148,929 | *Intron (PAFAH1B2)* | 164 | 8.65E-09 | 2.04E-02 | 2.03E-19 | 7.81E-19 | 4.13E-18 | 93 | 2.17E-17 | 5.66E-17 | rs964184, rs9804646, rs3135506, rs2266788 |
| | 11 | 117,147,930 | 117,149,929 | *Intron (PAFAH1B2)* | 165 | 8.78E-05 | 8.45E-01 | 3.27E-19 | 1.15E-18 | 6.11E-18 | 94 | 3.47E-17 | 9.13E-17 | rs964184, rs9804646, rs3135506, rs2266788 |
| | 11 | 117,181,930 | 117,183,929 | *SIDT2* | 178 | 9.03E-07 | 5.27E-01 | 3.46E-19 | 2.06E-18 | 1.04E-17 | 88 | 5.16E-17 | 1.34E-16 | rs964184, rs9804646, rs3135506, rs2266788 |
| | 11 | 117,182,930 | 117,184,929 | *SIDT2* | 160 | 8.43E-07 | 6.22E-02 | 3.03E-19 | 1.80E-18 | 9.04E-18 | 80 | 4.37E-17 | 1.14E-16 | rs964184, rs9804646, rs3135506, rs2266788 |
| | 19 | 44,882,528 | 44,884,527 | *Intron (NECTIN2)* | 145 | 2.87E-09 | 7.55E-01 | 1.36E-07 | 1.06E-08 | 2.18E-07 | 88 | 2.71E-02 | 8.28E-02 | rs12721054, rs5112, rs429358 |
| | 19 | 44,905,528 | 44,907,527 | *APOE* | 150 | 1.24E-11 | 7.71E-01 | 6.77E-08 | 6.12E-12 | 1.28E-05 | 57 | 4.10E-06 | 1.69E-05 | rs12721054, rs5112, rs429358 |
| | 1 | 55,045,498 | 55,047,497 | *PCSK9* | 161 | 2.52E-06 | 2.36E-03 | 2.75E-12 | 6.09E-12 | 4.77E-09 | 85 | 2.82E-03 | 1.16E-01 | rs11591147, rs28362263, rs505151, rs12117661, rs2495477 |
| | 1 | 55,046,498 | 55,048,497 | *PCSK9* | 167 | 2.02E-06 | 2.26E-05 | 1.92E-12 | 4.59E-12 | 3.34E-09 | 90 | 2.06E-03 | 1.49E-01 | rs11591147, rs28362263, rs505151, rs12117661, rs2495477 |
| | 1 | 55,062,498 | 55,064,497 | *PCSK9* | 186 | 1.40E-09 | 1.91E-02 | 1.06E-26 | 1.89E-26 | 1.28E-28 | 109 | 1.36E-10 | 4.33E-12 | rs11591147, rs28362263, rs505151, rs12117661, rs2495477 |
| | 1 | 55,063,498 | 55,065,497 | *PCSK9* | 167 | 2.40E-12 | 9.24E-02 | 7.26E-27 | 1.77E-26 | 1.24E-28 | 90 | 1.17E-10 | 3.56E-12 | rs11591147, rs28362263, rs505151, rs12117661, rs2495477 |
| | 1 | 55,291,498 | 55,293,497 | *Intergenic (GOT2P1)* | 167 | 2.22E-08 | 3.89E-01 | 5.35E-06 | 8.06E-09 | 9.94E-04 | 89 | 1.96E-01 | 2.98E-01 | rs11591147, rs28362263, rs505151, rs12117661, rs2495477 |
| | 1 | 55,333,498 | 55,335,497 | *Intergenic (GOT2P1)* | 175 | 1.63E-05 | 5.06E-03 | 4.95E-14 | 1.98E-13 | 3.83E-14 | 101 | 5.84E-07 | 1.88E-07 | rs11591147, rs28362263, rs505151, rs12117661, rs2495477 |
| | 1 | 55,334,498 | 55,336,497 | *Intergenic (GOT2P1)* | 149 | 3.45E-05 | 1.04E-02 | 5.11E-14 | 1.80E-13 | 3.49E-14 | 90 | 5.53E-07 | 1.78E-07 | rs11591147, rs28362263, rs505151, rs12117661, rs2495477 |
| | 19 | 10,689,528 | 10,691,527 | *ILF3* | 213 | 1.95E-06 | 1.33E-04 | 5.94E-10 | 2.16E-09 | 1.04E-01 | 121 | 6.70E-01 | 9.20E-01 | rs73015024, rs688, rs2278426, rs6511720 |
| | 19 | 10,690,528 | 10,692,527 | *ILF3* | 247 | 3.44E-07 | 7.96E-05 | 4.84E-10 | 1.75E-09 | 2.54E-02 | 120 | 5.62E-01 | 9.20E-01 | rs73015024, rs688, rs2278426, rs6511720 |
| | 19 | 10,701,528 | 10,703,527 | *QTRT1* | 122 | 1.24E-04 | 5.12E-02 | 8.72E-09 | 1.46E-08 | 5.13E-01 | 62 | 2.50E-01 | 3.56E-01 | rs73015024, rs688, rs2278426, rs6511720 |
| | 19 | 10,938,528 | 10,940,527 | *Intergenic (SMARCA4)* | 146 | 1.02E-05 | 1.86E-01 | 2.24E-11 | 6.02E-11 | 6.50E-01 | 76 | 9.51E-02 | 7.30E-02 | rs73015024, rs688, rs2278426, rs6511720 |
| | 19 | 10,939,528 | 10,941,527 | *Intergenic (SMARCA4)* | 150 | 1.12E-05 | 8.03E-01 | 2.85E-11 | 7.37E-11 | 2.61E-01 | 89 | 7.64E-02 | 6.05E-02 | rs73015024, rs688, rs2278426, rs6511720 |
| | 19 | 10,987,528 | 10,989,527 | *SMARCA4* | 180 | 3.17E-05 | 1.64E-01 | 1.99E-10 | 5.61E-10 | 1.55E-01 | 100 | 2.61E-01 | 2.62E-01 | rs73015024, rs688, rs2278426, rs6511720 |
| | 19 | 10,988,528 | 10,990,527 | *SMARCA4* | 161 | 3.64E-03 | 1.43E-01 | 2.25E-10 | 2.45E-10 | 1.31E-01 | 88 | 2.61E-01 | 3.12E-01 | rs73015024, rs688, rs2278426, rs6511720 |
| | 19 | 11,027,528 | 11,029,527 | *SMARCA4* | 179 | 1.67E-09 | 1.75E-01 | 2.53E-09 | 2.12E-09 | 5.40E-01 | 82 | 8.26E-01 | 9.92E-01 | rs73015024, rs688, rs2278426, rs6511720 |
| | 19 | 11,028,528 | 11,030,527 | *SMARCA4* | 183 | 7.34E-07 | 9.93E-01 | 3.25E-09 | 3.50E-09 | 2.05E-01 | 89 | 7.14E-01 | 9.14E-01 | rs73015024, rs688, rs2278426, rs6511720 |
| | 19 | 11,040,528 | 11,042,527 | *SMARCA4* | 171 | 2.27E-09 | 3.84E-01 | 1.32E-09 | 1.66E-09 | 1.23E-01 | 82 | 2.18E-03 | 6.77E-03 | rs73015024, rs688, rs2278426, rs6511720 |
| | 19 | 11,041,528 | 11,043,527 | *SMARCA4* | 162 | 7.89E-07 | 9.00E-01 | 1.21E-09 | 2.50E-09 | 7.56E-02 | 93 | 5.37E-01 | 8.19E-01 | rs73015024, rs688, rs2278426, rs6511720 |
| | 19 | 11,042,528 | 11,044,527 | *SMARCA4* | 146 | 1.03E-12 | 8.48E-02 | 8.32E-15 | 3.33E-15 | 1.55E-02 | 88 | 6.88E-01 | 8.08E-01 | rs73015024, rs688, rs2278426, rs6511720 |
| | 19 | 11,043,528 | 11,045,527 | *SMARCA4* | 187 | 2.29E-12 | 1.14E-01 | 1.00E-14 | 6.00E-15 | 2.48E-02 | 92 | 3.28E-01 | 4.34E-01 | rs73015024, rs688, rs2278426, rs6511720 |
| | 19 | 11,056,528 | 11,058,527 | *SMARCA4* | 178 | 3.68E-08 | 5.98E-02 | 4.22E-11 | 1.30E-10 | 3.08E-01 | 105 | 8.30E-01 | 6.82E-02 | rs73015024, rs688, rs2278426, rs6511720 |
| | 19 | 11,057,528 | 11,059,527 | *SMARCA4* | 181 | 2.85E-08 | 3.51E-01 | 3.51E-11 | 1.29E-10 | 1.38E-01 | 93 | 3.16E-01 | 3.24E-02 | rs73015024, rs688, rs2278426, rs6511720 |
| | 19 | 11,060,528 | 11,062,527 | *SMARCA4* | 201 | 3.08E-06 | 2.18E-01 | 8.12E-12 | 2.60E-11 | 7.72E-02 | 110 | 1.96E-01 | 7.38E-01 | rs73015024, rs688, rs2278426, rs6511720 |
| | 19 | 11,061,528 | 11,063,527 | *SMARCA4* | 196 | 7.56E-12 | 2.69E-03 | 6.35E-12 | 1.50E-11 | 2.82E-02 | 115 | 9.65E-01 | 8.76E-01 | rs73015024, rs688, rs2278426, rs6511720 |
| | 19 | 11,062,528 | 11,064,527 | *SMARCA4* | 165 | 1.59E-06 | 1.04E-01 | 2.90E-11 | 8.68E-11 | 5.32E-02 | 95 | 3.06E-01 | 4.66E-01 | rs73015024, rs688, rs2278426, rs6511720 |
| TC | 19 | 11,077,528 | 11,079,527 | *Intergenic (LDLR)* | 157 | 1.24E-05 | 5.86E-01 | 4.39E-07 | 7.57E-10 | 9.83E-01 | 89 | 8.30E-03 | 9.36E-01 | rs73015024, rs688, rs2278426, rs6511720 |
| | 19 | 11,084,528 | 11,086,527 | *Intergenic (LDLR)* | 147 | 1.57E-12 | 1.45E-01 | 1.73E-14 | 4.55E-14 | 1.72E-01 | 75 | 1.65E-01 | 2.48E-01 | rs73015024, rs688, rs2278426, rs6511720 |
| | 19 | 11,085,528 | 11,087,527 | *Intergenic (LDLR)* | 140 | 9.56E-07 | 4.18E-02 | 2.12E-14 | 5.35E-14 | 8.15E-02 | 77 | 7.98E-02 | 1.61E-01 | rs73015024, rs688, rs2278426, rs6511720 |
| | 19 | 11,087,528 | 11,089,527 | *LDLR* | 139 | 4.99E-07 | 1.68E-01 | 3.08E-16 | 8.88E-16 | 2.71E-02 | 63 | 3.72E-01 | 5.16E-01 | rs73015024, rs688, rs2278426, rs6511720 |
| | 19 | 11,088,528 | 11,090,527 | *LDLR* | 161 | 1.73E-06 | 2.73E-01 | 3.51E-16 | 1.11E-15 | 3.34E-02 | 84 | 2.66E-01 | 2.37E-01 | rs73015024, rs688, rs2278426, rs6511720 |
| | 19 | 11,097,528 | 11,099,527 | *LDLR* | 117 | 9.45E-10 | 3.99E-01 | 4.19E-15 | 1.47E-14 | 3.47E-02 | 71 | 5.84E-02 | 8.76E-01 | rs73015024, rs688, rs2278426, rs6511720 |
| | 19 | 11,098,528 | 11,100,527 | *LDLR* | 136 | 1.36E-09 | 1.55E-01 | 3.81E-15 | 1.48E-14 | 4.77E-02 | 78 | 3.31E-02 | 8.89E-01 | rs73015024, rs688, rs2278426, rs6511720 |
| | 19 | 11,118,528 | 11,120,527 | *LDLR* | 173 | 7.34E-11 | 1.05E-05 | 5.74E-08 | 1.02E-09 | 3.15E-08 | 105 | 5.91E-03 | 3.91E-02 | rs73015024, rs688, rs2278426, rs6511720 |
| | 19 | 11,119,528 | 11,121,527 | *LDLR* | 158 | 9.68E-10 | 9.45E-06 | 3.02E-08 | 4.26E-09 | 1.87E-08 | 88 | 1.75E-02 | 2.93E-02 | rs73015024, rs688, rs2278426, rs6511720 |
| | 19 | 11,169,528 | 11,171,527 | *KANK2* | 171 | 3.18E-05 | 2.02E-01 | 2.71E-11 | 1.04E-10 | 2.58E-02 | 89 | 2.36E-01 | 8.50E-01 | rs73015024, rs688, rs2278426, rs6511720 |
| | 19 | 11,170,528 | 11,172,527 | *KANK2* | 162 | 3.97E-07 | 5.27E-01 | 2.37E-11 | 7.65E-11 | 9.04E-02 | 80 | 3.12E-02 | 2.87E-01 | rs73015024, rs688, rs2278426, rs6511720 |
| | 19 | 11,210,528 | 11,212,527 | *DOCK6* | 144 | 4.89E-10 | 5.78E-02 | 5.67E-09 | 2.38E-09 | 2.80E-02 | 84 | 3.07E-02 | 6.02E-02 | rs73015024, rs688, rs2278426, rs6511720 |
| | 19 | 11,211,528 | 11,213,527 | *DOCK6* | 148 | 1.16E-09 | 6.11E-03 | 3.57E-09 | 3.54E-09 | 3.40E-02 | 77 | 4.99E-02 | 4.28E-01 | rs73015024, rs688, rs2278426, rs6511720 |
| | 19 | 11,319,528 | 11,321,527 | *Intron (TSPAN16)* | 218 | 3.46E-11 | 2.07E-02 | 1.38E-10 | 8.95E-11 | 4.00E-04 | 133 | 1.93E-01 | 2.27E-01 | rs73015024, rs688, rs2278426, rs6511720 |
| | 19 | 11,320,528 | 11,322,527 | *TSPAN16* | 222 | 1.34E-10 | 3.42E-02 | 1.42E-10 | 1.74E-10 | 7.19E-04 | 139 | 2.24E-01 | 5.57E-01 | rs73015024, rs688, rs2278426, rs6511720 |
| | 19 | 44,891,528 | 44,893,527 | *TOMM40* | 171 | 6.94E-11 | 2.44E-04 | 1.30E-08 | 2.57E-11 | 7.09E-01 | 85 | 1.99E-02 | 7.23E-01 | rs7412, rs429358, rs12721054 |
| | 19 | 44,892,528 | 44,894,527 | *TOMM40* | 175 | 1.91E-11 | 1.23E-02 | 2.63E-09 | 2.15E-10 | 2.47E-01 | 93 | 1.85E-03 | 8.62E-01 | rs7412, rs429358, rs12721054 |
| | 19 | 44,893,528 | 44,895,527 | *TOMM40* | 166 | 3.77E-17 | 3.27E-01 | 3.22E-09 | 4.44E-16 | 5.02E-08 | 82 | 4.85E-04 | 4.37E-01 | rs7412, rs429358, rs12721054 |
| | 19 | 44,894,528 | 44,896,527 | *Intron (TOMM40)* | 180 | 1.79E-10 | 8.31E-03 | 1.19E-06 | 2.73E-10 | 8.95E-08 | 97 | 2.68E-03 | 4.22E-01 | rs7412, rs429358, rs12721054 |
| | 19 | 44,897,528 | 44,899,527 | *Intron (TOMM40)* | 177 | 4.61E-06 | 2.26E-03 | 3.24E-09 | 1.15E-08 | 1.84E-01 | 110 | 3.23E-04 | 5.48E-02 | rs7412, rs429358, rs12721054 |
| | 19 | 44,898,528 | 44,900,527 | *Intron (TOMM40)* | 143 | 1.82E-06 | 6.26E-01 | 3.11E-09 | 1.16E-08 | 5.50E-01 | 81 | 1.67E-03 | 6.19E-01 | rs7412, rs429358, rs12721054 |

| | | | | | | | | | | | | | |
|---|---|---|---|---|---|---|---|---|---|---|---|---|---|
| 19 | 44,918,528 | 44,920,527 | *APOC1* | 181 | 2.88E-08 | 2.14E-02 | 5.13E-10 | 5.50E-10 | 1.68E-01 | 79 | 7.36E-01 | 6.28E-01 | rs7412, rs429358, rs12721054 |
| 19 | 44,919,528 | 44,921,527 | *Intergenic (APOC1)* | 184 | 4.76E-18 | 1.07E-10 | 1.88E-10 | 1.35E-17 | 9.67E-01 | 92 | 5.79E-02 | 6.32E-01 | rs7412, rs429358, rs12721054 |
| 19 | 44,920,528 | 44,922,527 | *Intergenic (APOC1)* | 168 | 5.16E-11 | 2.08E-05 | 2.94E-10 | 2.25E-10 | 9.75E-01 | 93 | 1.53E-03 | 8.65E-01 | rs7412, rs429358, rs12721054 |
| 19 | 44,930,528 | 44,932,527 | *APOC1P1* | 173 | 2.78E-06 | 5.94E-02 | 9.40E-08 | 1.43E-08 | 2.04E-01 | 109 | 6.65E-06 | 9.18E-02 | rs7412, rs429358, rs12721054 |
| 19 | 44,931,528 | 44,933,527 | *Intergenic (APOC1P1)* | 151 | 8.89E-07 | 1.90E-01 | 8.70E-08 | 1.02E-08 | 2.97E-01 | 95 | 1.56E-05 | 1.99E-01 | rs7412, rs429358, rs12721054 |

**Supplementary Table 9.** 2-kb sliding window analysis results of both unconditional analysis and anlaysis conditional on known common, low-frequency and rare variants. 21,015 discovery samples and 9,123 replication samples from the NHLBI Trans-Omics for Precision Medicine (TOPMed) program were considered in the analysis. The 9 replicated conditionally significant sliding windows (adjust for known common and low-frequent variants) are presented. Chr (chromosome); Start Location (start location of the 2-kb sliding window); End Location (end location of the 2-kb sliding window); #SNV (number of rare variants (MAF < 1%) in the 2-kb sliding window); STAAR-O (STAAR-O *P*-value); LDL-C (low-density lipoprotein cholesterol); HDL-C (High-density lipoprotein cholesterol); TG (triglycerides); TC (total cholesterol); Variants Adjusted (adjusted variants in conditional analysis). Physical positions of each window are on build hg38.

| Trait | Chr | Start Location | End Location | Gene | Discovery | | | Replication | | | Variants Adjusted |
|---|---|---|---|---|---|---|---|---|---|---|---|
| | | | | | #SNV | STAAR-O (Unconditional) | STAAR-O (Conditional) | #SNV | STAAR-O (Unconditional) | STAAR-O (Conditional) | |
| LDL-C | 1 | 55,333,498 | 55,335,497 | *Intergenic (GOT2P1)* | 171 | 6.66E-16 | 2.02E-02 | 95 | 1.27E-06 | 3.90E-02 | rs12117661, rs11591147, rs67608943, rs72646508, rs472495, rs28362261, rs28362263, rs505151, rs28362286 |
| LDL-C | 1 | 55,334,498 | 55,336,497 | *Intergenic (GOT2P1)* | 148 | 5.55E-16 | 2.76E-02 | 81 | 1.20E-06 | 3.95E-02 | rs12117661, rs11591147, rs67608943, rs72646508, rs472495, rs28362261, rs28362263, rs505151, rs28362286 |
| HDL-C | 11 | 116,802,930 | 116,804,929 | *Intergenic (ZPR1)* | 135 | 1.25E-08 | 9.94E-03 | 76 | 9.49E-05 | 9.75E-01 | rs964184,rs138326449,rs12269901,rs142953140 |
| HDL-C | 11 | 117,146,930 | 117,148,929 | *Intron (PAFAH1B2)* | 165 | 5.98E-09 | 9.69E-01 | 98 | 6.02E-04 | 2.35E-01 | rs964184,rs138326449,rs12269901,rs142953140 |
| HDL-C | 11 | 117,147,930 | 117,149,929 | *Intron (PAFAH1B2)* | 168 | 8.85E-09 | 3.79E-01 | 96 | 8.72E-04 | 9.62E-01 | rs964184,rs138326449,rs12269901,rs142953140 |
| TG | 11 | 117,146,930 | 117,148,929 | *Intron (PAFAH1B2)* | 164 | 7.81E-19 | 4.29E-18 | 93 | 2.17E-17 | 5.63E-17 | rs964184, rs75198898, rs2266788, rs3135506, rs9804646 |
| TG | 11 | 117,147,930 | 117,149,929 | *Intron (PAFAH1B2)* | 165 | 1.15E-18 | 6.34E-18 | 94 | 3.47E-17 | 9.09E-17 | rs964184, rs75198898, rs2266788, rs3135506, rs9804646 |
| TC | 1 | 55,333,498 | 55,335,497 | *Intergenic (GOT2P1)* | 175 | 1.98E-13 | 7.16E-03 | 101 | 5.84E-07 | 7.48E-02 | rs12117661, rs11591147, rs67608943, rs2495477, rs28362263, rs505151, rs28362286 |
| TC | 1 | 55,334,498 | 55,336,497 | *Intergenic (GOT2P1)* | 149 | 1.80E-13 | 5.30E-03 | 90 | 5.53E-07 | 5.49E-02 | rs12117661, rs11591147, rs67608943, rs2495477, rs28362263, rs505151, rs28362286 |

**Supplementary Table 10.** Dynamic window analysis results of both unconditional analysis and anlaysis conditional on known common and low-frequency variants. 21,015 discovery samples and 9,123 replication samples from the NHLBI Trans-Omics for Precision Medicine (TOPMed) program were considered in the analysis. Results for the significant sliding windows (unconditional genome-wide error rate < 0.05) using discovery samples are presented. Chr (chromosome); Start Location (start location of the dynamic window); End Location (end location of the dynamic window); #SNV (number of rare variants (MAF < 1%) in the dynamic window); STAAR-S (STAAR-S *P*-value); GWER (genome-wide error rate); LDL-C (low-density lipoprotein cholesterol); HDL-C (High-density lipoprotein cholesterol); TG (triglycerides); TC (total cholesterol); Variants Adjusted (adjusted variants in conditional analysis). Physical positions of each window are on build hg38.

| Trait | Chr | Start Location | End Location | Gene | Discovery | | | | Replication | | | Variants Adjusted |
|---|---|---|---|---|---|---|---|---|---|---|---|---|
| | | | | | #SNV | GWER | STAAR-S (Unconditional) | STAAR-S (Conditional) | #SNV | STAAR-S (Unconditional) | STAAR-S (Conditional) | |
| | 1 | 54,982,518 | 54,986,806 | *TMEM61* | 300 | <0.0005 | 7.20E-12 | 1.46E-06 | 178 | 4.20E-01 | 5.51E-01 | rs11591147, rs28362263, rs505151, rs12117661, rs472495 |
| | 1 | 55,046,481 | 55,046,987 | *PCSK9* | 50 | <0.0005 | 1.11E-14 | 1.08E-10 | 27 | 1.75E-03 | 2.90E-02 | rs11591147, rs28362263, rs505151, rs12117661, rs472495 |
| | 1 | 55,057,440 | 55,059,168 | *PCSK9* | 140 | 0.0005 | 1.34E-10 | 8.34E-08 | 74 | 6.02E-03 | 1.58E-02 | rs11591147, rs28362263, rs505151, rs12117661, rs472495 |
| | 1 | 55,063,375 | 55,063,715 | *PCSK9* | 40 | <0.0005 | 1.18E-33 | 3.10E-36 | 19 | 2.11E-12 | 6.71E-14 | rs11591147, rs28362263, rs505151, rs12117661, rs472495 |
| | 1 | 55,291,895 | 55,293,502 | *Intergenic (PCSK9)* | 140 | 0.0025 | 3.08E-10 | 1.99E-04 | 64 | 6.69E-02 | 3.53E-01 | rs11591147, rs28362263, rs505151, rs12117661, rs472495 |
| | 1 | 55,335,150 | 55,335,701 | *Intergenic (GOT2P1)* | 40 | <0.0005 | 8.58E-18 | 7.49E-19 | 21 | 9.29E-07 | 4.80E-07 | rs11591147, rs28362263, rs505151, rs12117661, rs472495 |
| | 19 | 10,690,690 | 10,691,276 | *ILF3* | 60 | 0.0045 | 4.53E-10 | 7.41E-02 | 27 | 5.74E-01 | 8.55E-01 | rs12151108, rs688, rs6511720 |
| | 19 | 10,701,935 | 10,704,176 | *QTRT1* | 140 | 0.0015 | 1.94E-10 | 1.30E-01 | 85 | 8.20E-02 | 3.08E-01 | rs12151108, rs688, rs6511720 |
| | 19 | 10,939,669 | 10,940,283 | *Intergenic (SMARCA4)* | 40 | <0.0005 | 2.20E-11 | 1.13E-01 | 23 | 6.90E-02 | 1.36E-01 | rs12151108, rs688, rs6511720 |
| | 19 | 10,988,220 | 10,990,055 | *SMARCA4* | 140 | <0.0005 | 7.86E-12 | 5.38E-02 | 74 | 2.82E-02 | 4.99E-02 | rs12151108, rs688, rs6511720 |
| | 19 | 11,028,128 | 11,030,669 | *SMARCA4* | 230 | <0.0005 | 5.31E-12 | 1.22E-01 | 109 | 3.62E-04 | 1.67E-02 | rs12151108, rs688, rs6511720 |
| | 19 | 11,039,183 | 11,042,158 | *SMARCA4* | 260 | <0.0005 | 7.20E-11 | 2.22E-01 | 112 | 2.54E-02 | 9.42E-02 | rs12151108, rs688, rs6511720 |
| | 19 | 11,044,139 | 11,047,012 | *SMARCA4* | 280 | <0.0005 | 5.62E-17 | 2.40E-02 | 140 | 2.02E-01 | 3.12E-01 | rs12151108, rs688, rs6511720 |
| | 19 | 11,057,659 | 11,058,062 | *SMARCA4* | 40 | <0.0005 | 6.72E-12 | 1.14E-02 | 18 | 1.14E-01 | 3.39E-01 | rs12151108, rs688, rs6511720 |
| | 19 | 11,061,825 | 11,062,376 | *SMARCA4* | 60 | <0.0005 | 6.76E-14 | 4.29E-03 | 30 | 8.73E-02 | 6.11E-01 | rs12151108, rs688, rs6511720 |
| | 19 | 11,062,626 | 11,063,132 | *SMARCA4* | 40 | <0.0005 | 7.67E-12 | 1.48E-01 | 24 | 7.95E-02 | 2.66E-01 | rs12151108, rs688, rs6511720 |
| | 19 | 11,076,615 | 11,080,398 | *Intergenic (LDLR)* | 290 | <0.0005 | 1.14E-12 | 4.59E-02 | 156 | 2.94E-02 | 4.22E-01 | rs12151108, rs688, rs6511720 |
| | 19 | 11,080,965 | 11,083,431 | *Intergenic (LDLR)* | 190 | 0.0395 | 2.92E-09 | 3.16E-02 | 111 | 1.91E-02 | 4.61E-01 | rs12151108, rs688, rs6511720 |
| | 19 | 11,085,211 | 11,085,673 | *Intergenic (LDLR)* | 40 | <0.0005 | 3.88E-21 | 1.83E-02 | 23 | 1.60E-01 | 2.96E-01 | rs12151108, rs688, rs6511720 |
| | 19 | 11,089,332 | 11,091,125 | *LDLR* | 150 | <0.0005 | 1.20E-17 | 7.56E-03 | 77 | 2.92E-01 | 3.90E-01 | rs12151108, rs688, rs6511720 |
| LDL-C | 19 | 11,099,458 | 11,100,236 | *LDLR* | 50 | <0.0005 | 7.35E-18 | 3.62E-05 | 22 | 6.51E-03 | 5.43E-02 | rs12151108, rs688, rs6511720 |
| | 19 | 11,113,482 | 11,114,026 | *LDLR* | 40 | 0.042 | 3.13E-09 | 4.76E-03 | 24 | 1.74E-02 | 5.91E-02 | rs12151108, rs688, rs6511720 |
| | 19 | 11,118,910 | 11,120,647 | *LDLR* | 140 | <0.0005 | 1.17E-14 | 1.24E-11 | 82 | 4.88E-05 | 1.82E-04 | rs12151108, rs688, rs6511720 |
| | 19 | 11,170,154 | 11,170,656 | *KANK2* | 40 | 0.0015 | 2.57E-10 | 8.74E-03 | 25 | 4.66E-01 | 5.44E-01 | rs12151108, rs688, rs6511720 |
| | 19 | 11,252,192 | 11,253,418 | *DOCK6* | 120 | <0.0005 | 3.25E-12 | 4.89E-05 | 58 | 3.31E-01 | 3.86E-01 | rs12151108, rs688, rs6511720 |
| | 19 | 11,319,992 | 11,320,870 | *Intron (TSPAN16)* | 60 | 0.02 | 1.44E-09 | 3.16E-05 | 41 | 5.04E-01 | 5.10E-01 | rs12151108, rs688, rs6511720 |
| | 19 | 44,761,480 | 44,764,386 | *Intergenic (SNORA70)* | 220 | 0.0145 | 1.11E-09 | 2.48E-01 | 105 | 6.15E-01 | 7.51E-01 | rs7412, rs429358, rs35136575 |
| | 19 | 44,794,403 | 44,796,975 | *Intron (CBLC)* | 220 | <0.0005 | 4.21E-11 | 3.30E-01 | 105 | 6.46E-02 | 2.49E-02 | rs7412, rs429358, rs35136575 |
| | 19 | 44,799,323 | 44,799,706 | *Intorn (CBLC)* | 40 | 0.0165 | 1.27E-09 | 4.65E-01 | 23 | 5.28E-03 | 2.37E-01 | rs7412, rs429358, rs35136575 |
| | 19 | 44,827,581 | 44,829,103 | *Intergenic (NECTIN2)* | 100 | 0.0035 | 4.19E-10 | 5.48E-01 | 56 | 1.54E-01 | 8.46E-01 | rs7412, rs429358, rs35136575 |
| | 19 | 44,858,550 | 44,862,878 | *NECTIN2* | 300 | 0.018 | 1.32E-09 | 1.08E-03 | 150 | 8.45E-03 | 4.41E-02 | rs7412, rs429358, rs35136575 |
| | 19 | 44,868,901 | 44,873,030 | *NECTIN2* | 300 | 0.0025 | 3.39E-10 | 8.89E-04 | 163 | 2.21E-02 | 1.54E-01 | rs7412, rs429358, rs35136575 |
| | 19 | 44,882,954 | 44,886,763 | *NECTIN2* | 300 | <0.0005 | 1.70E-11 | 1.36E-08 | 168 | 1.29E-01 | 3.81E-01 | rs7412, rs429358, rs35136575 |
| | 19 | 44,891,486 | 44,895,103 | *TOMM40* | 290 | <0.0005 | 7.99E-36 | 1.44E-11 | 145 | 3.26E-06 | 2.37E-01 | rs7412, rs429358, rs35136575 |
| | 19 | 44,898,834 | 44,899,255 | *Intron (TOMM40)* | 40 | <0.0005 | 1.51E-19 | 5.87E-01 | 24 | 6.43E-07 | 8.86E-02 | rs7412, rs429358, rs35136575 |
| | 19 | 44,901,341 | 44,901,824 | *TOMM40* | 50 | <0.0005 | 5.90E-12 | 3.00E-01 | 31 | 3.62E-01 | 1.93E-01 | rs7412, rs429358, rs35136575 |
| | 19 | 44,905,923 | 44,907,217 | *APOE* | 90 | <0.0005 | 1.60E-12 | 9.22E-12 | 35 | 8.70E-02 | 5.74E-02 | rs7412, rs429358, rs35136575 |
| | 19 | 44,907,264 | 44,908,411 | *APOE* | 80 | 0.0005 | 9.11E-11 | 3.52E-12 | 34 | 1.94E-01 | 2.00E-01 | rs7412, rs429358, rs35136575 |
| | 19 | 44,908,574 | 44,911,921 | *APOE* | 290 | <0.0005 | 2.62E-12 | 8.92E-13 | 121 | 7.28E-02 | 6.44E-02 | rs7412, rs429358, rs35136575 |
| | 19 | 44,919,667 | 44,923,321 | *Intergenic (APOC1)* | 300 | <0.0005 | 6.31E-32 | 3.16E-02 | 156 | 2.12E-05 | 3.75E-01 | rs7412, rs429358, rs35136575 |
| | 19 | 44,931,619 | 44,935,295 | *Intergenic (APOC1P1)* | 260 | <0.0005 | 2.38E-14 | 1.27E-03 | 145 | 7.72E-08 | 1.32E-03 | rs7412, rs429358, rs35136575 |
| | 11 | 116,830,552 | 116,831,135 | *APOC3* | 40 | <0.0005 | 1.11E-14 | 3.49E-13 | 29 | 1.83E-09 | 5.53E-09 | rs964184, rs12269901 |
| | 11 | 116,866,780 | 116,867,288 | *Intron (SIK3)* | 40 | 0.0295 | 2.24E-09 | 8.45E-09 | 19 | 2.22E-05 | 5.46E-05 | rs964184, rs12269901 |
| HDL-C | 11 | 116,928,564 | 116,929,045 | *Intron (PAFAH1B2)* | 40 | 0.0025 | 1.50E-10 | 4.43E-10 | 18 | 7.81E-04 | 1.06E-03 | rs964184, rs12269901 |
| | 16 | 56,954,390 | 56,957,607 | *Intergenic (CETP)* | 240 | <0.0005 | 1.66E-13 | 1.63E-01 | 122 | 4.72E-06 | 7.10E-01 | rs247616, rs5883, rs7499892, rs17231520, rs5880 |
| | 16 | 56,970,772 | 56,974,225 | *CETP* | 290 | <0.0005 | 2.63E-15 | 8.77E-09 | 157 | 3.38E-05 | 2.67E-01 | rs247616, rs5883, rs7499892, rs17231520, rs5880 |
| | 11 | 116,667,290 | 116,669,099 | *Intergenic (AP000770.1)* | 170 | 0.003 | 2.07E-10 | 2.33E-02 | 117 | 6.31E-02 | 2.17E-01 | rs964184, rs9804646, rs3135506, rs2266788 |
| | 11 | 116,707,856 | 116,708,989 | *Intergenic (AP000770.1)* | 110 | <0.0005 | 2.34E-13 | 6.97E-03 | 56 | 3.94E-03 | 6.66E-01 | rs964184, rs9804646, rs3135506, rs2266788 |
| | 11 | 116,748,452 | 116,751,869 | *BUD13* | 280 | <0.0005 | 2.11E-14 | 5.00E-05 | 127 | 1.74E-04 | 2.24E-02 | rs964184, rs9804646, rs3135506, rs2266788 |
| | 11 | 116,756,044 | 116,759,808 | *BUD13* | 290 | 0.0195 | 1.36E-09 | 2.77E-01 | 152 | 3.84E-04 | 5.73E-01 | rs964184, rs9804646, rs3135506, rs2266788 |
| | 11 | 116,764,132 | 116,767,275 | *BUD13* | 240 | 0.0335 | 2.23E-09 | 1.33E-01 | 113 | 6.69E-04 | 6.19E-01 | rs964184, rs9804646, rs3135506, rs2266788 |
| | 11 | 116,778,853 | 116,779,606 | *ZPR1* | 60 | <0.0005 | 3.89E-12 | 4.46E-12 | 40 | 2.70E-09 | 3.43E-08 | rs964184, rs9804646, rs3135506, rs2266788 |
| | 11 | 116,789,243 | 116,792,720 | *APOA5* | 270 | <0.0005 | 5.23E-12 | 2.06E-08 | 132 | 1.95E-05 | 5.92E-04 | rs964184, rs9804646, rs3135506, rs2266788 |
| TG | 11 | 116,830,498 | 116,831,114 | *APOC3* | 40 | <0.0005 | 2.40E-33 | 8.46E-32 | 28 | 5.53E-27 | 6.49E-26 | rs964184, rs9804646, rs3135506, rs2266788 |
| | 11 | 117,147,061 | 117,148,086 | *Intron (PAFAH1B2)* | 80 | <0.0005 | 5.10E-16 | 8.55E-15 | 41 | 9.48E-19 | 3.44E-18 | rs964184, rs9804646, rs3135506, rs2266788 |
| | 11 | 117,182,856 | 117,183,310 | *Intron (SIDT2)* | 40 | <0.0005 | 3.96E-12 | 1.08E-11 | 15 | 3.77E-14 | 6.53E-14 | rs964184, rs9804646, rs3135506, rs2266788 |
| | 11 | 117,349,560 | 117,350,171 | *Intron (CEP164)* | 50 | 0.013 | 1.08E-09 | 1.26E-09 | 29 | 4.12E-11 | 6.39E-11 | rs964184, rs9804646, rs3135506, rs2266788 |
| | 19 | 44,882,972 | 44,886,717 | *NECTIN2* | 300 | 0.027 | 1.70E-09 | 6.15E-07 | 174 | 9.27E-03 | 2.88E-01 | rs12721054, rs5112, rs429358 |
| | 19 | 44,906,026 | 44,907,048 | *APOE* | 80 | <0.0005 | 8.44E-13 | 4.44E-06 | 30 | 1.72E-05 | 5.20E-05 | rs12721054, rs5112, rs429358 |
| | 19 | 44,912,630 | 44,915,305 | *APOC1* | 260 | 0.003 | 2.13E-10 | 9.78E-02 | 140 | 5.69E-02 | 7.09E-01 | rs12721054, rs5112, rs429358 |
| | 1 | 54,982,463 | 54,986,097 | *TMEM61* | 240 | 0.0155 | 8.58E-10 | 1.31E-04 | 142 | 3.26E-01 | 5.24E-01 | rs11591147, rs28362263, rs505151, rs12117661, rs2495477 |
| | 1 | 55,046,513 | 55,046,818 | *PCSK9* | 40 | <0.0005 | 4.47E-14 | 3.60E-11 | 25 | 1.80E-04 | 8.28E-03 | rs11591147, rs28362263, rs505151, rs12117661, rs2495477 |
| | 1 | 55,063,387 | 55,063,740 | *PCSK9* | 40 | <0.0005 | 8.87E-28 | 6.67E-30 | 21 | 5.30E-12 | 1.63E-13 | rs11591147, rs28362263, rs505151, rs12117661, rs2495477 |
| | 1 | 55,291,905 | 55,293,502 | *Intergenic (GOT2P1)* | 140 | 0.0055 | 3.17E-10 | 8.77E-05 | 68 | 4.76E-01 | 2.30E-01 | rs11591147, rs28362263, rs505151, rs12117661, rs2495477 |
| | 1 | 55,335,119 | 55,335,584 | *Intergenic (GOT2P1)* | 40 | <0.0005 | 1.63E-15 | 4.44E-16 | 26 | 2.23E-07 | 7.03E-08 | rs11591147, rs28362263, rs505151, rs12117661, rs2495477 |
| | 19 | 10,690,690 | 10,691,129 | *ILF3* | 50 | 0.0125 | 6.60E-10 | 1.41E-01 | 24 | 6.64E-01 | 8.90E-01 | rs73015024, rs688, rs2278426, rs6511720 |
| | 19 | 10,702,032 | 10,704,176 | *QTRT1* | 140 | 0.0085 | 3.99E-10 | 1.41E-01 | 87 | 1.24E-01 | 4.05E-01 | rs73015024, rs688, rs2278426, rs6511720 |
| | 19 | 10,939,669 | 10,940,283 | *Intergenic (SMARCA4)* | 40 | 0.0005 | 8.21E-12 | 1.58E-01 | 23 | 3.31E-01 | 3.11E-01 | rs73015024, rs688, rs2278426, rs6511720 |
| | 19 | 10,989,311 | 10,990,058 | *SMARCA4* | 40 | 0.0005 | 1.98E-11 | 4.87E-03 | 23 | 8.85E-02 | 1.69E-01 | rs73015024, rs688, rs2278426, rs6511720 |
| | 19 | 11,028,636 | 11,030,669 | *SMARCA4* | 190 | 0.0005 | 3.18E-11 | 1.07E-01 | 96 | 2.20E-01 | 7.18E-01 | rs73015024, rs688, rs2278426, rs6511720 |
| | 19 | 11,039,067 | 11,042,158 | *SMARCA4* | 270 | 0.001 | 9.56E-11 | 1.11E-01 | 125 | 7.79E-04 | 8.47E-03 | rs73015024, rs688, rs2278426, rs6511720 |
| | 19 | 11,044,078 | 11,046,273 | *SMARCA4* | 230 | <0.0005 | 1.67E-16 | 3.00E-02 | 107 | 4.12E-01 | 6.70E-01 | rs73015024, rs688, rs2278426, rs6511720 |
| | 19 | 11,057,659 | 11,058,062 | *SMARCA4* | 40 | 0.0005 | 5.99E-12 | 8.82E-03 | 18 | 4.46E-01 | 8.18E-01 | rs73015024, rs688, rs2278426, rs6511720 |
| | 19 | 11,061,835 | 11,062,369 | *SMARCA4* | 60 | <0.0005 | 6.92E-13 | 3.41E-03 | 32 | 1.46E-01 | 5.46E-01 | rs73015024, rs688, rs2278426, rs6511720 |
| TC | 19 | 11,062,558 | 11,062,989 | *SMARCA4* | 40 | 0.0005 | 1.49E-11 | 1.42E-02 | 25 | 2.76E-01 | 6.57E-01 | rs73015024, rs688, rs2278426, rs6511720 |
| | 19 | 11,076,654 | 11,080,398 | *Intergenic (LDLR)* | 290 | 0.001 | 1.08E-10 | 2.53E-01 | 163 | 3.43E-02 | 7.60E-01 | rs73015024, rs688, rs2278426, rs6511720 |
| | 19 | 11,085,212 | 11,085,656 | *Intergenic (LDLR)* | 40 | <0.0005 | 3.18E-20 | 2.01E-02 | 22 | 1.14E-01 | 4.95E-01 | rs73015024, rs688, rs2278426, rs6511720 |
| | 19 | 11,089,332 | 11,091,204 | *LDLR* | 160 | <0.0005 | 1.67E-16 | 2.41E-02 | 86 | 3.89E-01 | 2.35E-01 | rs73015024, rs688, rs2278426, rs6511720 |
| | 19 | 11,099,458 | 11,100,236 | *LDLR* | 50 | <0.0005 | 3.64E-16 | 1.26E-03 | 23 | 1.01E-02 | 3.82E-02 | rs73015024, rs688, rs2278426, rs6511720 |
| | 19 | 11,118,898 | 11,120,556 | *LDLR* | 140 | 0.004 | 1.95E-10 | 1.46E-08 | 86 | 1.75E-03 | 2.12E-02 | rs73015024, rs688, rs2278426, rs6511720 |
| | 19 | 11,169,980 | 11,170,656 | *KANK2* | 60 | <0.0005 | 3.17E-12 | 1.36E-03 | 31 | 1.10E-01 | 2.82E-01 | rs73015024, rs688, rs2278426, rs6511720 |
| | 19 | 11,211,092 | 11,214,437 | *DOCK6* | 230 | 0.0035 | 1.91E-10 | 2.38E-03 | 126 | 1.82E-01 | 7.53E-01 | rs73015024, rs688, rs2278426, rs6511720 |
| | 19 | 11,252,192 | 11,253,246 | *DOCK6* | 100 | 0.001 | 8.82E-11 | 3.33E-04 | 49 | 7.26E-02 | 1.18E-01 | rs73015024, rs688, rs2278426, rs6511720 |
| | 19 | 11,309,640 | 11,310,134 | *Intron (TSPAN16)* | 40 | 0.033 | 2.07E-09 | 1.70E-03 | 20 | 1.19E-02 | 7.38E-02 | rs73015024, rs688, rs2278426, rs6511720 |
| | 19 | 11,319,627 | 11,320,925 | *Intron (TSPAN16)* | 110 | <0.0005 | 2.95E-12 | 2.32E-05 | 75 | 3.40E-01 | 5.90E-01 | rs73015024, rs688, rs2278426, rs6511720 |
| | 19 | 11,320,995 | 11,321,274 | *TSPAN16* | 40 | 0.032 | 1.93E-09 | 9.84E-04 | 23 | 7.09E-02 | 3.09E-01 | rs73015024, rs688, rs2278426, rs6511720 |
| | 19 | 44,891,807 | 44,895,104 | *TOMM40* | 280 | <0.0005 | 3.56E-24 | 6.80E-07 | 142 | 6.79E-05 | 6.96E-01 | rs7412, rs429358, rs12721054 |
| | 19 | 44,898,755 | 44,899,179 | *Intron (TOMM40)* | 40 | <0.0005 | 5.82E-15 | 7.07E-01 | 22 | 4.81E-05 | 7.52E-01 | rs7412, rs429358, rs12721054 |
| | 19 | 44,919,785 | 44,923,374 | *Intergenic (APOC1)* | 300 | <0.0005 | 7.23E-20 | 1.85E-01 | 165 | 3.68E-03 | 4.99E-01 | rs7412, rs429358, rs12721054 |
| | 19 | 44,931,072 | 44,932,057 | *Intergenic (APOC1P1)* | 80 | 0.0115 | 5.64E-10 | 3.20E-01 | 50 | 5.17E-07 | 8.60E-02 | rs7412, rs429358, rs12721054 |

**Supplementary Table 11.** Dynamic window analysis results of both unconditional analysis and anlaysis conditional on known common, low-frequency and rare variants. 21,015 discovery samples and 9,123 replication samples from the NHLBI Trans-Omics for Precision Medicine (TOPMed) program were considered in the analysis. The 7 replicated conditionally significant sliding windows (adjust for known common and low-frequent variants) are presented. Chr (chromosome); Start Location (start location of the dynamic window); End Location (end location of the dynamic window); #SNV (number of rare variants (MAF < 1%) in the dynamic window); STAAR-S (STAAR-S *P* -value); GWER (genome-wide error rate); LDL-C (low-density lipoprotein cholesterol); TG (triglycerides); TC (total cholesterol); Variants Adjusted (adjusted variants in conditional analysis). Physical positions of each window are on build hg38.

| Trait | Chr | Start Location | End Location | Gene | Discovery | | | | Replication | | | Variants Adjusted |
|---|---|---|---|---|---|---|---|---|---|---|---|---|
| | | | | | #SNV | GWER | STAAR-S (Unconditional) | STAAR-S (Conditional) | #SNV | STAAR-S (Unconditional) | STAAR-S (Conditional) | |
| LDL-C | 1 | 55,335,150 | 55,335,701 | *Intergenic (GOT2P1)* | 40 | <0.0005 | 8.58E-18 | 2.52E-02 | 21 | 9.29E-07 | 8.00E-01 | rs12117661, rs11591147, rs67608943, rs72646508, rs472495, rs28362261, rs28362263, rs505151, rs28362286 |
| HDL-C | 11 | 116,866,780 | 116,867,288 | *Intron (SIK3)* | 40 | 0.0295 | 2.24E-09 | 1.33E-03 | 19 | 2.22E-05 | 3.76E-02 | rs964184,rs138326449,rs12269901,rs142953140 |
| HDL-C | 11 | 116,928,564 | 116,929,045 | *Intron (PAFAH1B2)* | 40 | 0.0025 | 1.50E-10 | 5.25E-04 | 18 | 7.81E-04 | 6.08E-01 | rs964184,rs138326449,rs12269901,rs142953140 |
| TG | 11 | 117,147,061 | 117,148,086 | *Intron (PAFAH1B2)* | 80 | <0.0005 | 5.10E-16 | 9.99E-15 | 41 | 9.48E-19 | 3.29E-18 | rs964184, rs75198898, rs2266788, rs3135506, rs9804646 |
| TG | 11 | 117,182,856 | 117,183,310 | *Intron (SIDT2)* | 40 | <0.0005 | 3.96E-12 | 1.12E-11 | 15 | 3.77E-14 | 6.37E-14 | rs964184, rs75198898, rs2266788, rs3135506, rs9804646 |
| TG | 11 | 117,349,560 | 117,350,171 | *Intron (CEP164)* | 50 | 0.013 | 1.08E-09 | 1.44E-09 | 29 | 4.12E-11 | 7.12E-11 | rs964184, rs75198898, rs2266788, rs3135506, rs9804646 |
| TC | 1 | 55,335,119 | 55,335,584 | *Intergenic (GOT2P1)* | 40 | <0.0005 | 1.63E-15 | 2.80E-02 | 26 | 2.23E-07 | 4.19E-01 | rs12117661, rs11591147, rs67608943, rs2495477, rs28362263, rs505151, rs28362286 |

### Supplementary Note

##### TOPMed study participants and acknowledgements

##### Discovery phase (n = 21,015)

<u>Framingham Heart Study (FHS)</u>

The FHS is a three generational prospective cohort that has been described in detail previously[1]. Individuals were initially recruited in 1948 in Framingham, USA to evaluate cardiovascular disease risk factors. The second generation cohort (5,124 offspring of the original cohort) was recruited between 1971 and 1975[2,3]. The third generation cohort (4,095 grandchildren of the original cohort) was collected between 2002 and 2005. Fasting lipid levels were measured at exam 1 of the Offspring (1971-1975) and third generation (2002-2005) cohorts, using standard LRC protocols.

Whole genome sequencing (WGS) for the Trans-Omics in Precision Medicine (TOPMed) program was supported by the National Heart, Lung and Blood Institute (NHLBI). WGS for "NHLBI TOPMed: Whole Genome Sequencing and Related Phenotypes in the Framingham Heart Study" (phs000974.v1.p1) was performed at the Broad Institute of MIT and Harvard (HHSN268201500014C).

The Framingham Heart Study (FHS) acknowledges the support of contracts NO1-HC-25195, HHSN268201500001I and 75N92019D00031 from the National Heart, Lung and Blood Institute and grant supplement R01 HL092577-06S1 for this research. We also acknowledge the dedication of the FHS study participants without whom this research would not be possible. Dr. Vasan is supported in part by the Evans Medical Foundation and the Jay and Louis Coffman Endowment from the Department of Medicine, Boston University School of Medicine.

<u>Old Order Amish (OOA)</u>

The Amish Complex Disease Research Program includes a set of large community-based studies focused largely on cardiometabolic health carried out in the Old Order Amish (OOA) community of Lancaster, Pennsylvania[4]. The OOA population of

Lancaster County, PA immigrated to the Colonies from Western Europe in the early 1700's. There are now over 38,000 OOA individuals in the Lancaster area, nearly all of whom can trace their ancestry back 12-14 generations to approximately 400 founders. Investigators at the University of Maryland School of Medicine have been studying the genetic determinants of cardiometabolic health in this population since 1993. To date, over 8,000 Amish adults have participated in one or more of our studies.
The 1,123 Amish subjects included in the TOPMed program were enrolled in studies supported by NIH grants R01 AG18728, U01 HL072515, R01 HL088119, R01 HL121007, and P30 DK072488. WGS for “NHLBI TOPMed: Genetics of Cardiometabolic Health in the Amish” (phs000956) was performed at the Broad Institute of MIT and Harvard (3R01HL121007-01S1).

Jackson Heart Study (JHS)

The JHS is a large, population-based observational study evaluating the etiology of cardiovascular, renal, and respiratory diseases among African Americans residing in the three counties (Hinds, Madison, and Rankin) that make up the Jackson, Mississippi metropolitan area[5,6]. Data and biologic materials have been collected from 5,306 participants, including a nested family cohort of 1,498 members of 264 families. The age at enrollment for the unrelated cohort was 35-84 years; the family cohort included related individuals >21 years old. Participants provided an extensive medical and social history and had an array of physical and biochemical measurements and diagnostic procedures, and a subset of participants provided genomic DNA during a baseline examination (2000-2004) and two follow-up examinations (2005-2008 and 2009-2012), with a fourth examination ongoing. Annual follow-up interviews and cohort surveillance are ongoing.

Whole genome sequencing (WGS) for the Trans-Omics in Precision Medicine (TOPMed) program was supported by the National Heart, Lung and Blood Institute (NHLBI). WGS for “NHLBI TOPMed: The Jackson Heart Study” (phs000964.v1.p1) was performed at the University of Washington Northwest Genomics Center (HHSN268201100037C).

The Jackson Heart Study (JHS) is supported and conducted in collaboration with Jackson State University (HHSN268201800013I), Tougaloo College (HHSN268201800014I), the Mississippi State Department of Health (HHSN268201800015I/HHSN26800001) and the University of Mississippi Medical Center (HHSN268201800010I, HHSN268201800011I and HHSN268201800012I) contracts from the National Heart, Lung, and Blood Institute (NHLBI) and the National Institute for Minority Health and Health Disparities (NIMHD). The authors also wish to thank the staffs and participants of the JHS.

Multi-Ethnic Study of Atherosclerosis (MESA)

The Multi-Ethnic Study of Atherosclerosis is a National Heart, Lung and Blood Institute-sponsored, population-based investigation of subclinical cardiovascular disease and its progression[7]. A total of 6,814 individuals, aged 45 to 84 years, were recruited from six US communities (Baltimore City and County, MD; Chicago, IL; Forsyth County, NC; Los Angeles County, CA; New York, NY; and St. Paul, MN) between July 2000 and August 2002. Participants were excluded if they had physician-diagnosed cardiovascular disease prior to enrollment, including angina, myocardial infarction, heart failure, stroke or TIA, resuscitated cardiac arrest or a cardiovascular intervention (e.g., CABG, angioplasty, valve replacement, or pacemaker/defibrillator placement). Pre-specified recruitment plans identified four racial/ethnic groups (White European-American, African-American, Hispanic-American, and Chinese-American) for enrollment, with targeted oversampling of minority groups to enhance statistical power.

Whole genome sequencing (WGS) for the Trans-Omics in Precision Medicine (TOPMed) program was supported by the National Heart, Lung and Blood Institute (NHLBI). WGS for “NHLBI TOPMed: Multi-Ethnic Study of Atherosclerosis (MESA)” (phs001416.v1.p1) was performed at the Broad Institute of MIT and Harvard (3U54HG003067-13S1). Centralized read mapping and genotype calling, along with variant quality metrics and filtering were provided by the TOPMed Informatics Research Center (3R01HL-117626-02S1; contract HHSN268201800002I). Phenotype harmonization, data management, sample-identity QC, and general study coordination, were provided by the TOPMed Data Coordinating Center (3R01HL-120393-02S1; contract HHSN268201800001I).

The MESA project is conducted and supported by the National Heart, Lung, and Blood Institute (NHLBI) in collaboration with MESA investigators. Support for MESA is provided by contracts 75N92020D00001, HHSN268201500003I, N01-HC-95159, 75N92020D00005, N01-HC-95160, 75N92020D00002, N01-HC-95161, 75N92020D00003, N01-HC-95162, 75N92020D00006, N01-HC-95163, 75N92020D00004, N01-HC-95164, 75N92020D00007, N01-HC-95165, N01-HC-95166, N01-HC-95167, N01-HC-95168, N01-HC-95169, UL1-TR-000040, UL1-TR-001079, UL1-TR-001420. Also supported in part by the National Center for Advancing Translational Sciences, CTSI grant UL1TR001881, and the National Institute of Diabetes and Digestive and Kidney Disease Diabetes Research Center (DRC) grant DK063491 to the Southern California Diabetes Endocrinology Research Center.

MESA Family is conducted and supported by the National Heart, Lung, and Blood Institute (NHLBI) in collaboration with MESA investigators. Support is provided by grants and contracts R01HL071051, R01HL071205, R01HL071250, R01HL071251, R01HL071258, R01HL071259, by the National Center for Research Resources, Grant UL1RR033176. Also supported in part by the National Center for Advancing Translational Sciences, CTSI grant UL1TR001881, and the National Institute of Diabetes and Digestive and Kidney Disease Diabetes Research Center (DRC) grant DK063491 to the Southern California Diabetes Endocrinology Research Center.

Genome-wide Association Study of Adiposity in Samoans (Samoan)

The parent Samoan study is a population-based genome-wide association study (GWAS) of adiposity and cardiometabolic phenotypes among adults, 25-65 years of age, from the independent nation of Samoa in the South Pacific. The research goal of this study is to identify genetic variation that increases susceptibility to obesity and cardiometabolic phenotypes. Biomarker and questionnaire data were collected to assess cardiometabolic phenotypes. DNA was collected and the Affymetrix 6.0 chip used for SNP genotyping. After quality control checks on genotyping and excluding individuals with key missing data we have a final sample of 3,122 adults with high-quality genome-wide marker data[8]. Participation in TOPMed provided whole genome sequence data for 1,285 individuals from the GWAS sample chosen for maximal informativity for our Samoan-specific imputation panel.

Whole genome sequencing (WGS) for the Trans-Omics in Precision Medicine (TOPMed) program was supported by the National Heart, Lung and Blood Institute (NHLBI). WGS for “NHLBI TOPMed: Genome-wide Association Study of Adiposity in Samoans” (phs000972) was performed at the University of Washington Northwest Genomics Center (HHSN268201100037C) and the New York Genome Center (HHSN268201500016C).

Data collection was funded by NIH grant R01-HL093093 and R01-HL133040. We thank the Samoan participants of the study and local village authorities. We acknowledge the support of the Samoan Ministry of Health and the Samoa Bureau of Statistics for their support of this research.

<u>Women’s Health Initiative (WHI)</u>

The Women's Health Initiative (WHI) is a large study of postmenopausal women’s health investigating risk factors for cancer, CVD, age-related fractures and chronic disease[9]. It began in 1993 as a set of randomized controlled clinical trials (CT) and an observational study (OS). Specifically, the CT (n=68,132) included three overlapping components: The Hormone Therapy (HT) Trials (n=27,347), Dietary Modification (DM) Trial (n=48,835), and Calcium and Vitamin D (CaD) Trial (n=36,282). Eligible women could be randomized into as many as all three CTs components. Women who were ineligible or unwilling to join the CT were then invited to join the OS (n=93,676).

Whole genome sequencing (WGS) for the Trans-Omics in Precision Medicine (TOPMed) program was supported by the National Heart, Lung and Blood Institute

(NHLBI). WGS for “NHLBI TOPMed: Women’s Health Initiative” (phs001237) was performed at the Broad Institute of MIT and Harvard (HHSN268201500014C).

The WHI program is funded by the National Heart, Lung, and Blood Institute, National Institutes of Health, U.S. Department of Health and Human Services through contracts 75N92021D00001, 75N92021D00002, 75N92021D00003, 75N92021D00004, 75N92021D00005. The authors thank the WHI investigators and staff for their dedication, and the study participants for making the program possible. A full listing of WHI investigators can be found at: http://www.whi.org/researchers/Documents%20%20Write%20a%20Paper/WHI%20Investigator%20Long%20List.pdf.

The content is solely the responsibility of the authors and does not necessarily represent the official views of the National Institutes of Health.

**Replication phase (n = 9,123)**

Atherosclerosis Risk in Communities Study (ARIC)

The ARIC study is a population-based prospective cohort study of cardiovascular disease sponsored by the National Heart, Lung, and Blood Institute (NHLBI). ARIC included 15,792 individuals, predominantly European American and African American, aged 45-64 years at baseline (1987-89), chosen by probability sampling from four US communities. Cohort members completed three additional triennial follow-up examinations, a fifth exam in 2011-2013, a sixth exam in 2016-2017, a seventh exam in 2018-2019, and an eighth exam in 2020. The ARIC study has been described in detail previously[10].

Whole genome sequencing (WGS) for the Trans-Omics in Precision Medicine (TOPMed) program was supported by the National Heart, Lung and Blood Institute (NHLBI). WGS for “NHLBI TOPMed: Atherosclerosis Risk in Communities (ARIC)” (phs001211) was performed at the Baylor College of Medicine Human Genome

Sequencing Center (HHSN268201500015C and 3U54HG003273-12S2) and the Broad Institute of MIT and Harvard (3R01HL092577-06S1). Centralized read mapping and genotype calling, along with variant quality metrics and filtering were provided by the TOPMed Informatics Research Center (3R01HL-117626-02S1; contract HHSN268201800002I). Phenotype harmonization, data management, sample-identity QC, and general study coordination, were provided by the TOPMed Data Coordinating Center (3R01HL-120393-02S1; contract HHSN268201800001I). We gratefully acknowledge the studies and participants who provided biological samples and data for TOPMed.

The Atherosclerosis Risk in Communities study has been funded in whole or in part with Federal funds from the National Heart, Lung, and Blood Institute, National Institutes of Health, Department of Health and Human Services (contract numbers HHSN268201700001I, HHSN268201700002I, HHSN268201700003I, HHSN268201700004I and HHSN268201700005I). The authors thank the staff and participants of the ARIC study for their important contributions.

Cleveland Family Study (CFS)

The CFS is a family-based longitudinal study that includes participants with laboratory diagnosed sleep apnea, their family members and neighborhood control families followed between 1990 and 2006. Four examinations over 16 years provided measurements of sleep apnea with overnight polysomnography, anthropometry, and other related phenotypes, as detailed previously[4,11]. After an overnight fast, blood was collected which was assayed for lipid levels at the University of Vermont Laboratory for Clinical Biochemistry Research. Lipids (triglycerides, HDL cholesterol) from fasted blood serum were measured by enzymatic methods using Centers for Disease Control and Prevention guidelines[12].

Whole genome sequencing (WGS) for the Trans-Omics in Precision Medicine (TOPMed) program was supported by the National Heart, Lung and Blood Institute (NHLBI). WGS for “NHLBI TOPMed: Cleveland Family Study” (phs000954) was performed at the University of Washington Northwest Genomics Center (3R01HL098433-05S1).

This research was supported by grants HL 046389; HL113338;1R35HL135818 from the National Heart, Lung, and Blood Institute (NHLBI).

Cardiovascular Health Study (CHS)

The Cardiovascular Health Study is a prospective population-based cohort study of risk factors for CHD and stroke in adults 65 years and older[13]. The main objective is to identify factors related to the onset and course of heart disease and stroke. The four Field Centers are located in Forsyth County, NC; Sacramento County, CA; Washington County, MD; and Pittsburgh, PA. The original cohort of 5201 elderly were recruited in 1989-1990; and in 1992-1993, 687 additional minority participants were recruited and examined. Each community sample was obtained from random samples of the Medicare eligibility lists of the Health Care Financing Administration (HCFA). Eligible to participate were persons living in the household of each sampled individual who were: 1) 65 yr or older; 2) non-institutionalized; 3) expected to remain in the area for 3 yr; and 4) able to give informed consent. Excluded were those wheelchair-bound, receiving hospice care or cancer treatment. The minority cohort was recruited using similar methods. Participants were eligible whether or not they had clinically apparent cardiovascular disease. Subjects were followed with semi-annual contacts, alternating between telephone calls and surveillance clinic visits.

Whole genome sequencing (WGS) for the Trans-Omics in Precision Medicine (TOPMed) program was supported by the National Heart, Lung and Blood Institute (NHLBI). WGS for “NHLBI TOPMed: Cardiovascular Health Study” (phs001368) was performed at the Baylor College of Medicine Human Genome Sequencing Center (HHSN268201500015C).

This research was supported by contracts HHSN268201200036C, HHSN268200800007C, HHSN268201800001C, N01HC55222, N01HC85079,

N01HC85080, N01HC85081, N01HC85082, N01HC85083, N01HC85086, 75N92021D00006, and grants U01HL080295 and U01HL130114 from the National Heart, Lung, and Blood Institute (NHLBI), with additional contribution from the National Institute of Neurological Disorders and Stroke (NINDS). Additional support was provided by R01AG023629 from the National Institute on Aging (NIA). A full list of principal CHS investigators and institutions can be found at CHS-NHLBI.org.

Diabetes Heart Study (DHS)

The Diabetes Heart Study (DHS) began as a family-based study enriched for type 2 diabetes (T2D). The initial cohort included 1443 European American and African American participants from 564 families with multiple cases of type 2 diabetes recruited between 1998 and 2006[14]. As an ancillary study, the African American Diabetes Heart Study (AA-DHS) expanded the total number of African Americans to 691 by recruiting additional unrelated participants with type 2 diabetes from 2007 and 2010[15]. All participants were extensively phenotyped for measures of subclinical CVD and other known CVD risk factors. Primary outcomes were quantified burden of vascular calcified plaque in the coronary artery, carotid artery, and abdominal aorta all determined from non-contrast computed tomography scans. For TOPMed, DHS and AA-DHS African American participants with CAC were selected for WGS, prioritizing the inclusion of families.

Whole genome sequencing (WGS) for the Trans-Omics in Precision Medicine (TOPMed) program was supported by the National Heart, Lung and Blood Institute (NHLBI). WGS for “NHLBI TOPMed: Diabetes Heart Study” (phs001412) was performed at the Broad Institute of MIT and Harvard (HHSN268201500014C).

This work was supported by R01 HL92301, R01 HL67348, R01 NS058700, R01 AR48797, R01 DK071891, R01 AG058921, the General Clinical Research Center of the Wake Forest University School of Medicine (M01 RR07122, F32 HL085989), the American Diabetes Association, and a pilot grant from the Claude Pepper Older

Americans Independence Center of Wake Forest University Health Sciences (P60 AG10484).

Genetic Study of Atherosclerosis Risk (GeneSTAR)

GeneSTAR is an ongoing family-based prospective study designed to determine environmental, phenotypic, and genetic causes of premature cardiovascular disease. GeneSTAR was originally conducted in healthy adult European- and African-American siblings of probands with documented early onset coronary disease under 60 years of age at the time of hospitalization in any of 10 Baltimore area hospitals from 1982-2006. Participants were screened for traditional coronary disease and stroke risk factors and have been followed regularly to ascertain incident cardiovascular disease[16]. Commencing in 2003, the siblings, their offspring, and the coparent of the offspring who were free of cardiovascular disease participated in a 2 week trial of aspirin 81 mg/day with pre and post ex vivo platelet function assessed using multiple agonists and were screened for traditional coronary disease and stroke risk factors[17]. Of the total 3949 participants, 1786 were selected for TOPMed prioritized on complete platelet function measures and largest family size.

Whole genome sequencing (WGS) for the Trans-Omics in Precision Medicine (TOPMed) program was supported by the National Heart, Lung and Blood Institute (NHLBI). WGS for “NHLBI TOPMed: Genetic Study of Atherosclerosis Risk” (phs001218) was performed at Psomagen (formerly Macrogen; 3R01HL112064-04S1), Illumina (R01HL112064), and the Broad Institute of MIT and Harvard (HHSN268201500014C).

GeneSTAR was supported by grants from the National Institutes of Health/National Heart, Lung, and Blood Institute (U01 HL72518, HL087698, HL49762, HL59684, HL58625, HL071025, HL112064), by a grant from the National Institutes of Health/National Institute of Nursing Research (NR0224103), and by a grant from the National Institutes of Health/National Center for Research Resources (M01-RR000052) to the Johns Hopkins General Clinical Research Center.

Genetic Epidemiology Network of Arteriopathy (GENOA)

The Genetic Epidemiology Network of Arteriopathy (GENOA) is one of four networks in the NHLBI Family-Blood Pressure Program (FBPP)[18]. GENOA's long-term objective is to elucidate the genetics of target organ complications of hypertension, including both atherosclerotic and arteriolosclerotic complications involving the heart, brain, kidneys, and peripheral arteries[19]. The longitudinal GENOA Study recruited European-American and African-American sibships with at least 2 individuals with clinically diagnosed essential hypertension before age 60 years. All other members of the sibship were invited to participate regardless of their hypertension status. Participants were diagnosed with hypertension if they had either 1) a previous clinical diagnosis of hypertension by a physician with current anti-hypertensive treatment, or 2) an average systolic blood pressure ≥ 140 mm Hg or diastolic blood pressure ≥ 90 mm Hg based on the second and third readings at the time of their clinic visit. Only participants of the African-American Cohort were sequenced through TOPMed.

During the first exam (Phase 1; 1996-2000), 1,583 European-Americans from Rochester, MN and 1,854 African-Americans from Jackson, MS were examined. Between 2000 and 2004 (Phase 2), 1,241 participants of the European-American Cohort and 1,482 participants of the African-American cohort returned for a second examination. The second examination of the European-American cohort included computed tomography scans for coronary artery calcification while the second examination of the African-American cohort included an echocardiogram. Between 2009 and 2011, an examination that included computed tomography scans for coronary artery calcification (CAC Study) was conducted on 752 participants of the African-American Cohort.

Every participant with an echocardiogram was selected for whole genome sequencing (WGS) through TOPMed. We then selected 106 African-American participants who had a computed tomography scan for coronary artery calcification but not an echocardiogram or were a sibling of someone already selected for WGS. Finally, we excluded individuals whom we knew were already being whole genome sequenced through TOPMed or another sequencing effort (GENOA participants who overlap with ARIC or JHS participants).

Support for GENOA was provided by the National Heart, Lung and Blood Institute (HL054457, HL054464, HL054481, HL119443, HL085571, and HL087660) of the National Institutes of Health. DNA extraction for "NHLBI TOPMed: Genetic Epidemiology Network of Arteriopathy" (phs001345) was performed at the Mayo Clinic Genotyping Core, and WGS was performed at the DNA Sequencing and Gene Analysis Center at the University of Washington (3R01HL055673-18S1) and the Broad Institute (HHSN268201500014C). We would like to thank the GENOA participants.

Genetics of Lipid Lowering Drugs and Diet Network (GOLDN)

GOLDN is a family-based study of European descent individuals recruited in Minneapolis and Salt Lake City (two of the NHLBI Family Heart Study sites). It aims to uncover genetic predictors of variability in lipid phenotypes, which include both fasting and postprandial lipids quantified using traditional methods, NMR, and high-throughput lipidomics. During the initial screening of ~1,350 individuals, the following criteria were used for exclusion: age < 18 years; fasting triglycerides ≥1500 mg/dL; recent history of myocardial infarction, coronary bypass surgery, or coronary angioplasty; self-report of a positive history of liver, kidney, pancreas, or gallbladder disease, or a history of nutrient malabsorption; current use of insulin; abnormal liver or kidney function; in women of childbearing potential, pregnancy, breastfeeding, not using an acceptable form of contraception. Of those who enrolled, 1,048 individuals consented to the use of their DNA in research; 893 participants with data on all exposures, outcomes, and covariates were included in the current study.

GOLDN biospecimens, baseline phenotype data, and intervention phenotype data were collected with funding from National Heart, Lung and Blood Institute (NHLBI) grant U01 HL072524. Whole-genome sequencing in GOLDN was funded by NHLBI grant R01 HL104135-04S1.

Whole genome sequencing (WGS) for the Trans-Omics in Precision Medicine (TOPMed) program was supported by the National Heart, Lung and Blood Institute (NHLBI). WGS for “NHLBI TOPMed: Genetics of Lipid Lowering Drugs and Diet Network” (phs001359) was performed at the University of Washington Northwest Genomics Center (3R01HL104135-04S1).

San Antonio Family Heart Study (SAFS)

The SAFHS began in 1991, and included 1,431 individuals in 42 extended families at baseline. Probands were 40 to 60 year old low-income Mexican Americans selected at random without regard to presence or absence of disease, almost exclusively from Mexican American census tracts in San Antonio, Texas. All first, second, and third degree relatives of the proband and of the proband's spouse, aged 16 years or above, were eligible to participate in the study. As part of our ongoing studies, we have recruited new family members from the original families, expanding the cohort to almost 3,099 individuals primarily from 73 families. Our study is a mixed longitudinal design. Subjects have been seen between 1 and 4 times with an average of 1.95 examinations.

Whole genome sequencing (WGS) for the Trans-Omics in Precision Medicine (TOPMed) program was supported by the National Heart, Lung and Blood Institute (NHLBI). WGS for “NHLBI TOPMed: San Antonio Family Heart Study” (phs001215) was performed at the Illumina Genomic Services (3R01HL113323-03S1).

Collection of the San Antonio Family Study data was supported in part by National Institutes of Health (NIH) grants R01 HL045522, MH078143, MH078111 and MH083824; and whole genome sequencing of SAFS subjects was supported by U01 DK085524 and R01 HL113323. We are very grateful to the participants of the San Antonio Family Study for their continued involvement in our research programs.

**NHLBI Trans-Omics for Precision Medicine (TOPMed) Consortium**

Namiko Abe[47], Gonçalo Abecasis[48], Francois Aguet[49], Christine Albert[50], Laura Almasy[51], Alvaro Alonso[52], Seth Ament[53], Peter Anderson[54], Pramod Anugu[55], Deborah Applebaum-Bowden[56], Kristin Ardlie[49], Dan Arking[57], Donna K Arnett[58], Allison Ashley-Koch[59], Stella Aslibekyan[60], Tim Assimes[61], Paul Auer[62], Dimitrios Avramopoulos[57], Najib Ayas[63], Adithya Balasubramanian[64], John Barnard[65], Kathleen Barnes[66], R. Graham Barr[67], Emily Barron-Casella[57], Lucas Barwick[68], Terri Beaty[57], Gerald Beck[69], Diane Becker[70], Lewis Becker[57], Rebecca Beer[71], Amber Beitelshees[53], Emelia Benjamin[72], Takis Benos[73], Marcos Bezerra[74], Larry Bielak[48], Joshua Bis[75], Thomas Blackwell[48], John Blangero[76], Eric Boerwinkle[77], Donald W. Bowden[78], Russell Bowler[79], Jennifer Brody[54], Ulrich Broeckel[80], Jai Broome[54], Deborah Brown[81], Karen Bunting[47], Esteban Burchard[82], Carlos Bustamante[83], Erin Buth[84], Brian Cade[85], Jonathan Cardwell[86], Vincent Carey[87], Julie Carrier[88], Cara Carty[89], Richard Casaburi[90], Juan P Casas Romero[91], James Casella[57], Peter Castaldi[92], Mark Chaffin[49], Christy Chang[53], Yi-Cheng Chang[93], Daniel Chasman[94], Sameer Chavan[86], Bo-Juen Chen[47], Wei-Min Chen[95], Yii-Der Ida Chen[96], Michael Cho[87], Seung Hoan Choi[49], Lee-Ming Chuang[97], Mina Chung[98], Ren-Hua Chung[99], Clary Clish[100], Suzy Comhair[101], Matthew Conomos[84], Elaine Cornell[102], Adolfo Correa[103], Carolyn Crandall[90], James Crapo[104], L. Adrienne Cupples[105], Joanne Curran[106], Jeffrey Curtis[48], Brian Custer[107], Coleen Damcott[53], Dawood Darbar[108], Sean David[109], Colleen Davis[54], Michelle Daya[86], Mariza de Andrade[110], Lisa de las Fuentes[111], Paul de Vries[112], Michael DeBaun[113], Ranjan Deka[114], Dawn DeMeo[87], Scott Devine[53], Huyen Dinh[64], Harsha Doddapaneni[115], Qing Duan[116], Shannon Dugan-Perez[64], Ravi Duggirala[117], Jon Peter Durda[102], Susan K. Dutcher[118], Charles Eaton[119], Lynette Ekunwe[55], Adel El Boueiz[120], Patrick Ellinor[121],

Leslie Emery[54], Serpil Erzurum[65], Charles Farber[95], Jesse Farek[64], Tasha Fingerlin[122], Matthew Flickinger[48], Myriam Fornage[77], Nora Franceschini[123], Chris Frazar[54], Mao Fu[53], Stephanie M. Fullerton[54], Lucinda Fulton[124], Stacey Gabriel[49], Weiniu Gan[71], Shanshan Gao[86], Yan Gao[55], Margery Gass[125], Heather Geiger[126], Bruce Gelb[127], Mark Geraci[128], Soren Germer[47], Robert Gerszten[129], Auyon Ghosh[87], Richard Gibbs[64], Chris Gignoux[61], Mark Gladwin[73], David Glahn[130], Stephanie Gogarten[54], Da-Wei Gong[53], Harald Goring[131], Sharon Graw[66], Kathryn J. Gray[132], Daniel Grine[86], Colin Gross[48], C. Charles Gu[124], Yue Guan[53], Xiuqing Guo[96], Namrata Gupta[49], David M. Haas[133], Jeff Haessler[125], Michael Hall[134], Yi Han[64], Patrick Hanly[135], Daniel Harris[136], Nicola L. Hawley[137], Jiang He[138], Ben Heavner[84], Susan Heckbert[54], Ryan Hernandez[82], David Herrington[139], Craig Hersh[140], Bertha Hidalgo[60], James Hixson[77], Brian Hobbs[87], John Hokanson[86], Elliott Hong[53], Karin Hoth[141], Chao (Agnes) Hsiung[142], Jianhong Hu[64], Yi-Jen Hung[143], Haley Huston[144], Chii Min Hwu[145], Marguerite Ryan Irvin[60], Rebecca Jackson[146], Deepti Jain[54], Cashell Jaquish[71], Jill Johnsen[147], Andrew Johnson[71], Craig Johnson[54], Rich Johnston[52], Kimberly Jones[57], Hyun Min Kang[148], Robert Kaplan[149], Sharon Kardia[48], Shannon Kelly[150], Eimear Kenny[127], Michael Kessler[53], Alyna Khan[54], Ziad Khan[64], Wonji Kim[151], John Kimoff[152], Greg Kinney[153], Barbara Konkle[144], Charles Kooperberg[125], Holly Kramer[154], Christoph Lange[155], Ethan Lange[86], Leslie Lange[86], Cathy Laurie[54], Cecelia Laurie[54], Meryl LeBoff[87], Jiwon Lee[87], Sandra Lee[64], Wen-Jane Lee[145], Jonathon LeFaive[48], David Levine[54], Dan Levy[71], Joshua Lewis[53], Xiaohui Li[96], Yun Li[116], Henry Lin[96], Honghuang Lin[156], Xihong Lin[157], Simin Liu[158], Yongmei Liu[159], Yu Liu[160], Ruth J.F. Loos[161], Steven Lubitz[121], Kathryn Lunetta[156], James Luo[71], Ulysses Magalang[162], Michael Mahaney[106], Barry Make[57], Ani Manichaikul[95], Alisa Manning[163], JoAnn Manson[87], Lisa Martin[164], Melissa Marton[126], Susan Mathai[86], Rasika Mathias[57], Susanne May[84], Patrick McArdle[53], Merry-Lynn McDonald[60], Sean McFarland[151], Stephen McGarvey[119], Daniel McGoldrick[165], Caitlin McHugh[84], Becky McNeil[166], Hao Mei[55], James Meigs[167], Vipin Menon[64], Luisa Mestroni[66], Ginger Metcalf[64], Deborah A Meyers[168], Emmanuel Mignot[169], Julie Mikulla[71], Nancy Min[55], Mollie Minear[170], Ryan L Minster[73], Braxton D. Mitchell[53], Matt Moll[92], Zeineen Momin[64], May E. Montasser[53], Courtney Montgomery[171], Donna Muzny[64], Josyf C Mychaleckyj[95], Girish Nadkarni[127], Rakhi Naik[57], Take Naseri[172], Pradeep Natarajan[49], Sergei Nekhai[173], Sarah C.

Nelson[84], Bonnie Neltner[86], Caitlin Nessner[64], Deborah Nickerson[174], Osuji Nkechinyere[64], Kari North[116], Jeff O'Connell[175], Tim O'Connor[53], Heather Ochs-Balcom[176], Geoffrey Okwuonu[64], Allan Pack[177], David T. Paik[178], Nicholette Palmer[179], James Pankow[180], George Papanicolaou[71], Cora Parker[181], Gina Peloso[182], Juan Manuel Peralta[117], Marco Perez[61], James Perry[53], Ulrike Peters[183], Patricia Peyser[48], Lawrence S Phillips[52], Jacob Pleiness[48], Toni Pollin[53], Wendy Post[184], Julia Powers Becker[185], Meher Preethi Boorgula[86], Michael Preuss[127], Bruce Psaty[54], Pankaj Qasba[71], Dandi Qiao[87], Zhaohui Qin[52], Nicholas Rafaels[186], Laura Raffield[187], Mahitha Rajendran[64], Vasan S. Ramachandran[156], D.C. Rao[124], Laura Rasmussen-Torvik[188], Aakrosh Ratan[95], Susan Redline[87], Robert Reed[53], Catherine Reeves[189], Elizabeth Regan[104], Alex Reiner[190], Muagututi'a Sefuiva Reupena[191], Ken Rice[54], Stephen Rich[95], Rebecca Robillard[192], Nicolas Robine[126], Dan Roden[193], Carolina Roselli[49], Jerome Rotter[96], Ingo Ruczinski[57], Alexi Runnels[126], Pamela Russell[86], Sarah Ruuska[144], Kathleen Ryan[53], Ester Cerdeira Sabino[194], Danish Saleheen[195], Shabnam Salimi[53], Sejal Salvi[64], Steven Salzberg[57], Kevin Sandow[196], Vijay G. Sankaran[197], Jireh Santibanez[64], Karen Schwander[124], David Schwartz[86], Frank Sciurba[73], Christine Seidman[198], Jonathan Seidman[199], Frédéric Sériès[200], Vivien Sheehan[201], Stephanie L. Sherman[202], Amol Shetty[53], Aniket Shetty[86], Wayne Hui-Heng Sheu[145], M. Benjamin Shoemaker[203], Brian Silver[204], Edwin Silverman[87], Robert Skomro[205], Albert Vernon Smith[206], Jennifer Smith[48], Josh Smith[54], Nicholas Smith[207], Tanja Smith[47], Sylvia Smoller[149], Beverly Snively[208], Michael Snyder[61], Tamar Sofer[87], Nona Sotoodehnia[54], Adrienne M. Stilp[54], Garrett Storm[209], Elizabeth Streeten[53], Jessica Lasky Su[87], Yun Ju Sung[124], Jody Sylvia[87], Adam Szpiro[54], Daniel Taliun[48], Hua Tang[210], Margaret Taub[57], Kent D. Taylor[211], Matthew Taylor[66], Simeon Taylor[53], Marilyn Telen[59], Timothy A. Thornton[54], Machiko Threlkeld[212], Lesley Tinker[125], David Tirschwell[54], Sarah Tishkoff[213], Hemant Tiwari[214], Catherine Tong[215], Russell Tracy[216], Michael Tsai[180], Dhananjay Vaidya[57], David Van Den Berg[217], Peter VandeHaar[48], Scott Vrieze[180], Tarik Walker[86], Robert Wallace[141], Avram Walts[86], Fei Fei Wang[54], Heming Wang[218], Jiongming Wang[219], Karol Watson[90], Jennifer Watt[64], Daniel E. Weeks[73], Joshua Weinstock[148], Bruce Weir[54], Scott T Weiss[220], Lu-Chen Weng[121], Jennifer Wessel[221], Cristen Willer[222], Kayleen Williams[84], L. Keoki Williams[223], Carla Wilson[87], James

Wilson[224], Lara Winterkorn[126], Quenna Wong[54], Joseph Wu[178], Huichun Xu[53], Lisa Yanek[57], Ivana Yang[86], Ketian Yu[48], Seyedeh Maryam Zekavat[49], Yingze Zhang[225], Snow Xueyan Zhao[104], Wei Zhao[226], Xiaofeng Zhu[227], Michael Zody[47], Sebastian Zoellner[48]

47 - New York Genome Center, New York, New York, 10013, US; 48 - University of Michigan, Ann Arbor, Michigan, 48109, US; 49 - Broad Institute, Cambridge, Massachusetts, 2142, US; 50 - Cedars Sinai, Boston, Massachusetts, 2114, US; 51 - Children's Hospital of Philadelphia, University of Pennsylvania, Philadelphia, Pennsylvania, 19104, US; 52 - Emory University, Atlanta, Georgia, 30322, US; 53 - University of Maryland, Baltimore, Maryland, 21201, US; 54 - University of Washington, Seattle, Washington, 98195, US; 55 - University of Mississippi, Jackson, Mississippi, 38677, US; 56 - National Institutes of Health, Bethesda, Maryland, 20892, US; 57 - Johns Hopkins University, Baltimore, Maryland, 21218, US; 58 - University of Kentucky, Lexington, Kentucky, 40506, US; 59 - Duke University, Durham, North Carolina, 27708, US; 60 - University of Alabama, Birmingham, Alabama, 35487, US; 61 - Stanford University, Stanford, California, 94305, US; 62 - University of Wisconsin Milwaukee, Milwaukee, Wisconsin, 53211, US; 63 - Providence Health Care, Medicine, Vancouver, CA; 64 - Baylor College of Medicine Human Genome Sequencing Center, Houston, Texas, 77030, US; 65 - Cleveland Clinic, Cleveland, Ohio, 44195, US; 66 - University of Colorado Anschutz Medical Campus, Aurora, Colorado, 80045, US; 67 - Columbia University, New York, New York, 10032, US; 68 - The Emmes Corporation, LTRC, Rockville, Maryland, 20850, US; 69 - Cleveland Clinic, Quantitative Health Sciences, Cleveland, Ohio, 44195, US; 70 - Johns Hopkins University, Medicine, Baltimore, Maryland, 21218, US; 71 - National Heart, Lung, and Blood Institute, National Institutes of Health, Bethesda, Maryland, 20892, US; 72 - Boston University, Massachusetts General Hospital, Boston University School of Medicine, Boston, Massachusetts, 2114, US; 73 - University of Pittsburgh, Pittsburgh, Pennsylvania, 15260, US; 74 - Fundação de Hematologia e Hemoterapia de Pernambuco - Hemope, Recife, 52011-000, BR; 75 - University of Washington, Cardiovascular Health Research Unit, Department of Medicine, Seattle, Washington, 98195, US; 76 - University of Texas Rio Grande Valley

School of Medicine, Human Genetics, Brownsville, Texas, 78520, US; 77 - University of Texas Health at Houston, Houston, Texas, 77225, US; 78 - Wake Forest Baptist Health, Department of Biochemistry, Winston-Salem, North Carolina, 27157, US; 79 - National Jewish Health, National Jewish Health, Denver, Colorado, 80206, US; 80 - Medical College of Wisconsin, Milwaukee, Wisconsin, 53226, US; 81 - University of Texas Health at Houston, Pediatrics, Houston, Texas, 77030, US; 82 - University of California, San Francisco, San Francisco, California, 94143, US; 83 - Stanford University, Biomedical Data Science, Stanford, California, 94305, US; 84 - University of Washington, Biostatistics, Seattle, Washington, 98195, US; 85 - Brigham & Women's Hospital, Brigham and Women's Hospital, Boston, Massachusetts, 2115, US; 86 - University of Colorado at Denver, Denver, Colorado, 80204, US; 87 - Brigham & Women's Hospital, Boston, Massachusetts, 2115, US; 88 - University of Montreal, US; 89 - Washington State University, Pullman, Washington, 99164, US; 90 - University of California, Los Angeles, Los Angeles, California, 90095, US; 91 - Brigham & Women's Hospital, US; 92 - Brigham & Women's Hospital, Medicine, Boston, Massachusetts, 2115, US; 93 - National Taiwan University, Taipei, 10617, TW; 94 - Brigham & Women's Hospital, Division of Preventive Medicine, Boston, Massachusetts, 2215, US; 95 - University of Virginia, Charlottesville, Virginia, 22903, US; 96 - Lundquist Institute, Torrance, California, 90502, US; 97 - National Taiwan University, National Taiwan University Hospital, Taipei, 10617, TW; 98 - Cleveland Clinic, Cleveland Clinic, Cleveland, Ohio, 44195, US; 99 - National Health Research Institute Taiwan, Miaoli County, 350, TW; 100 - Broad Institute, Metabolomics Platform, Cambridge, Massachusetts, 2142, US; 101 - Cleveland Clinic, Immunity and Immunology, Cleveland, Ohio, 44195, US; 102 - University of Vermont, Burlington, Vermont, 5405, US; 103 - University of Mississippi, Population Health Science, Jackson, Mississippi, 39216, US; 104 - National Jewish Health, Denver, Colorado, 80206, US; 105 - Boston University, Biostatistics, Boston, Massachusetts, 2115, US; 106 - University of Texas Rio Grande Valley School of Medicine, Brownsville, Texas, 78520, US; 107 - Vitalant Research Institute, San Francisco, California, 94118, US; 108 - University of Illinois at Chicago, Chicago, Illinois, 60607, US; 109 - University of Chicago, Chicago, Illinois, 60637, US; 110 - Mayo Clinic, Health Quantitative Sciences Research, Rochester,

Minnesota, 55905, US; 111 - Washington University in St Louis, Department of Medicine, Cardiovascular Division, St. Louis, Missouri, 63110, US; 112 - University of Texas Health at Houston, Human Genetics Center, Department of Epidemiology, Human Genetics, and Environmental Sciences, Houston, Texas, 77030, US; 113 - Vanderbilt University, Nashville, Tennessee, 37235, US; 114 - University of Cincinnati, Cincinnati, Ohio, 45220, US; 115 - Baylor College of Medicine Human Genome Sequencing Center, Houston, Texas, 77030; 116 - University of North Carolina, Chapel Hill, North Carolina, 27599, US; 117 - University of Texas Rio Grande Valley School of Medicine, Edinburg, Texas, 78539, US; 118 - Washington University in St Louis, Genetics, St Louis, Missouri, 63110, US; 119 - Brown University, Providence, Rhode Island, 2912, US; 120 - Harvard University, Channing Division of Network Medicine, Cambridge, Massachusetts, 2138, US; 121 - Massachusetts General Hospital, Boston, Massachusetts, 2114, US; 122 - National Jewish Health, Center for Genes, Environment and Health, Denver, Colorado, 80206, US; 123 - University of North Carolina, Epidemiology, Chapel Hill, North Carolina, 27599, US; 124 - Washington University in St Louis, St Louis, Missouri, 63130, US; 125 - Fred Hutchinson Cancer Research Center, Seattle, Washington, 98109, US; 126 - New York Genome Center, New York City, New York, 10013, US; 127 - Icahn School of Medicine at Mount Sinai, New York, New York, 10029, US; 128 - University of Pittsburgh, Pittsburgh, Pennsylvania, US; 129 - Beth Israel Deaconess Medical Center, Boston, Massachusetts, 2215, US; 130 - Boston Children's Hospital, Harvard Medical School, Department of Psychiatry, Boston, Massachusetts, 2115, US; 131 - University of Texas Rio Grande Valley School of Medicine, San Antonio, Texas, 78229, US; 132 - Mass General Brigham, Obstetrics and Gynecology, Boston, Massachusetts, 2115, US; 133 - Indiana University, OB/GYN, Indianapolis, Indiana, 46202, US; 134 - University of Mississippi, Cardiology, Jackson, Mississippi, 39216, US; 135 - University of Calgary, Medicine, Calgary, CA; 136 - University of Maryland, Genetics, Philadelphia, Pennsylvania, 19104, US; 137 - Yale University, Department of Chronic Disease Epidemiology, New Haven, Connecticut, 6520, US; 138 - Tulane University, New Orleans, Louisiana, 70118, US; 139 - Wake Forest Baptist Health, Winston-Salem, North Carolina, 27157, US; 140 - Brigham & Women's Hospital, Channing Division of

Network Medicine, Boston, Massachusetts, 2115, US; 141 - University of Iowa, Iowa City, Iowa, 52242, US; 142 - National Health Research Institute Taiwan, Institute of Population Health Sciences, NHRI, Miaoli County, 350, TW; 143 - Tri-Service General Hospital National Defense Medical Center, TW; 144 - Blood Works Northwest, Seattle, Washington, 98104, US; 145 - Taichung Veterans General Hospital Taiwan, Taichung City, 407, TW; 146 - Oklahoma State University Medical Center, Internal Medicine, DIvision of Endocrinology, Diabetes and Metabolism, Columbus, Ohio, 43210, US; 147 - Blood Works Northwest, Research Institute, Seattle, Washington, 98104, US; 148 - University of Michigan, Biostatistics, Ann Arbor, Michigan, 48109, US; 149 - Albert Einstein College of Medicine, New York, New York, 10461, US; 150 - University of California, San Francisco, San Francisco, California, 94118, US; 151 - Harvard University, Cambridge, Massachusetts, 2138, US; 152 - McGill University, Montréal, QC H3A 0G4, CA; 153 - University of Colorado at Denver, Epidemiology, Aurora, Colorado, 80045, US; 154 - Loyola University, Public Health Sciences, Maywood, Illinois, 60153, US; 155 - Harvard School of Public Health, Biostats, Boston, Massachusetts, 2115, US; 156 - Boston University, Boston, Massachusetts, 2215, US; 157 - Harvard School of Public Health, Boston, Massachusetts, 2115, US; 158 - Brown University, Epidemiology and Medicine, Providence, Rhode Island, 2912, US; 159 - Duke University, Cardiology, Durham, North Carolina, 27708, US; 160 - Stanford University, Cardiovascular Institute, Stanford, California, 94305, US; 161 - Icahn School of Medicine at Mount Sinai, The Charles Bronfman Institute for Personalized Medicine, New York, New York, 10029, US; 162 - Ohio State University, Division of Pulmonary, Critical Care and Sleep Medicine, Columbus, Ohio, 43210, US; 163 - Broad Institute, Harvard University, Massachusetts General Hospital; 164 - George Washington University, cardiology, Washington, District of Columbia, 20037, US; 165 - University of Washington, Genome Sciences, Seattle, Washington, 98195, US; 166 - RTI International, US; 167 - Massachusetts General Hospital, Medicine, Boston, Massachusetts, 2114, US; 168 - University of Arizona, Tucson, Arizona, 85721, US; 169 - Stanford University, Center For Sleep Sciences and Medicine, Palo Alto, California, 94304, US; 170 - National Institute of Child Health and Human Development, National Institutes of Health, Bethesda, Maryland, 20892, US; 171 - Oklahoma Medical Research Foundation, Genes and Human Disease, Oklahoma

City, Oklahoma, 73104, US; 172 - Ministry of Health, Government of Samoa, Apia, WS; 173 - Howard University, Washington, District of Columbia, 20059, US; 174 - University of Washington, Department of Genome Sciences, Seattle, Washington, 98195, US; 175 - University of Maryland, Balitmore, Maryland, 21201, US; 176 - University at Buffalo, Buffalo, New York, 14260, US; 177 - University of Pennsylvania, Division of Sleep Medicine/Department of Medicine, Philadelphia, Pennsylvania, 19104-3403, US; 178 - Stanford University, Stanford Cardiovascular Institute, Stanford, California, 94305, US; 179 - Wake Forest Baptist Health, Biochemistry, Winston-Salem, North Carolina, 27157, US; 180 - University of Minnesota, Minneapolis, Minnesota, 55455, US; 181 - RTI International, Biostatistics and Epidemiology Division, Research Triangle Park, North Carolina, 27709-2194, US; 182 - Boston University, Department of Biostatistics, Boston, Massachusetts, 2118, US; 183 - Fred Hutchinson Cancer Research Center, Fred Hutch and UW, Seattle, Washington, 98109, US; 184 - Johns Hopkins University, Cardiology/Medicine, Baltimore, Maryland, 21218, US; 185 - University of Colorado at Denver, Medicine, Denver, Colorado, 80204, US; 186 - University of Colorado at Denver, Denver, Colorado, 80045, US; 187 - University of North Carolina, Genetics, Chapel Hill, North Carolina, 27599, US; 188 - Northwestern University, Chicago, Illinois, 60208, US; 189 - New York Genome Center, New York Genome Center, New York City, New York, 10013, US; 190 - Fred Hutchinson Cancer Research Center, University of Washington, Seattle, Washington, 98109, US; 191 - Lutia I Puava Ae Mapu I Fagalele, Apia, WS; 192 - University of Ottawa, Sleep Research Unit, University of Ottawa Institute for Mental Health Research, Ottawa, ON K1Z 7K4, CA; 193 - Vanderbilt University, Medicine, Pharmacology, Biomedicla Informatics, Nashville, Tennessee, 37235, US; 194 - Universidade de Sao Paulo, Faculdade de Medicina, Sao Paulo, 1310000, BR; 195 - Columbia University, New York, New York, 10027, US; 196 - Lundquist Institute, TGPS, Torrance, California, 90502, US; 197 - Harvard University, Division of Hematology/Oncology, Boston, Massachusetts, 2115, US; 198 - Harvard Medical School, Genetics, Boston, Massachusetts, 2115, US; 199 - Harvard Medical School, Boston, Massachusetts, 2115, US; 200 - Université Laval, Quebec City, G1V 0A6, CA; 201 - Emory University, Pediatrics, Atlanta, Georgia, 30307, US; 202 - Emory University, Human Genetics, Atlanta, Georgia, 30322, US; 203 - Vanderbilt University,

Medicine/Cardiology, Nashville, Tennessee, 37235, US; 204 - UMass Memorial Medical Center, Worcester, Massachusetts, 1655, US; 205 - University of Saskatchewan, Saskatoon, SK S7N 5C9, CA; 206 - University of Michigan; 207 - University of Washington, Epidemiology, Seattle, Washington, 98195, US; 208 - Wake Forest Baptist Health, Biostatistical Sciences, Winston-Salem, North Carolina, 27157, US; 209 - University of Colorado at Denver, Genomic Cardiology, Aurora, Colorado, 80045, US; 210 - Stanford University, Genetics, Stanford, California, 94305, US; 211 - Lundquist Institute, Institute for Translational Genomics and Populations Sciences, Torrance, California, 90502, US; 212 - University of Washington, University of Washington, Department of Genome Sciences, Seattle, Washington, 98195, US; 213 - University of Pennsylvania, Genetics, Philadelphia, Pennsylvania, 19104, US; 214 - University of Alabama, Biostatistics, Birmingham, Alabama, 35487, US; 215 - University of Washington, Department of Biostatistics, Seattle, Washington, 98195, US; 216 - University of Vermont, Pathology & Laboratory Medicine, Burlington, Vermont, 5405, US; 217 - University of Southern California, USC Methylation Characterization Center, University of Southern California, California, 90033, US; 218 - Brigham & Women's Hospital, Mass General Brigham, Boston, Massachusetts, 2115, US; 219 - University of Michigan, US; 220 - Brigham & Women's Hospital, Channing Division of Network Medicine, Department of Medicine, Boston, Massachusetts, 2115, US; 221 - Indiana University, Epidemiology, Indianapolis, Indiana, 46202, US; 222 - University of Michigan, Internal Medicine, Ann Arbor, Michigan, 48109, US; 223 - Henry Ford Health System, Detroit, Michigan, 48202, US; 224 - Beth Israel Deaconess Medical Center, Cardiology, Cambridge, Massachusetts, 2139, US; 225 - University of Pittsburgh, Medicine, Pittsburgh, Pennsylvania, 15260, US; 226 - University of Michigan, Department of Epidemiology, Ann Arbor, Michigan, 48109, US; 227 - Case Western Reserve University, Department of Population and Quantitative Health Sciences, Cleveland, Ohio, 44106, US

**TOPMed Lipids Working Group**

Moustafa Abdalla, Gonçalo Abecasis, Donna K Arnett, Stella Aslibekyan, Tim Assimes, Elizabeth Atkinson, Christie Ballantyne, Wei Bao, David Beame, Amber Beitelshees,

Larry Bielak, Joshua Bis, Corneliu Bodea, Eric Boerwinkle, Donald W. Bowden, Michael Bowers, Jennifer Brody, Brian Cade, Sarah Calvo, Jenna Carlson, I-Shou Chang, Yii-Der Ida Chen, Sung Chun, Ren-Hua Chung, Adolfo Correa, L. Adrienne Cupples, Coleen Damcott, Paul de Vries, Ana F. Diallo, Ron Do, Amanda Elliott, Mao Fu, Andrea Ganna, Dawei Gong, Sarah Graham, Mary Haas, Bernhard Haring, Jiang He, Susan Heckbert, Blanca Himes, James Hixson, Marguerite Ryan Irvin, Deepti Jain, Gail Jarvik, Jicai Jiang, Paule Valery Joseph, Goo Jun, Rita Kalyani, Sharon Kardia, Sekar Kathiresan, Addison Keely, Amit Khera, Sumeet Khetarpal, Derek Klarin, Charles Kooperberg, Satoshi Koyama, Brian Kral, Leslie Lange, Cathy Laurie, Cecelia Laurie, Rozenn Lemaitre, Zilin Li, Xihao Li, Xihong Lin, Yingchang Lu, Michael Mahaney, Ani Manichaikul, Lisa Martin, Rasika Mathias, Ravi Mathur, Stephen McGarvey, Caitlin McHugh, John McLenithan, Julie Mikulla, Braxton D. Mitchell, May E. Montasser, Vamsi Mootha, Andrew Moran, Alanna C Morrison, Tetsushi Nakao, Pradeep Natarajan, Deborah Nickerson, Kari North, Jeff O'Connell, Christopher O'Donnell, Nicholette Palmer, Akhil Pampana, Kaavya Paruchuri, Aniruddh Patel, Gina Peloso, James Perry, Ulrike Peters, Mary Pettinger, Patricia Peyser, James Pirruccello, Toni Pollin, Michael Preuss, Bruce Psaty, Jennifer Anne Purnell, Susan Redline, Robert Reed, Alex Reiner, Stephen Rich, Samantha Rosenthal, Jerome Rotter, Jenny Schoenberg, Margaret Sunitha Selvaraj, Wayne Hui-Heng Sheu, Jennifer Smith, Tamar Sofer, Adrienne M. Stilp, Shamil R Sunyaev, Ida Surakka, Carole Sztalryd, Hua Tang, Kent D. Taylor, Mark Trinder, Michael Tsai, Md Mesbah Uddin, Sarah Urbut, Marie Verbanck, Ann Von Holle, Heming Wang, Yuxuan Wang, Kate Wehr, Kerri Wiggins, John Wilkins, Cristen Willer, James Wilson, Brooke Wolford, Huichun Xu, Lisa Yanek, Zhi Yu, Norann Zaghloul, Seyedeh Maryam Zekavat, Jingwen Zhang, Ying Zhou

**The Samoan Obesity, Lifestyle and Genetic Adaptations Study (OLaGA) Group**

Ranjan Deka, Dept. of Environmental Health, University of Cincinnati;
Nicola L. Hawley, Dept. of Chronic Disease Epidemiology, Yale University;
Stephen T McGarvey, Dept. of Epidemiology and International Health Institute, and Dept. of Anthropology, Brown University;
Ryan L Minster, Dept. of Human Genetics, University of Pittsburgh;

Take Naseri, Ministry of Health, Government of Samoa;
Muagututi‘a Sefuiva Reupena, Lutia I Puava Ae Mapu I Fagalele;
Daniel E. Weeks, Depts. of Human Genetics and Biostatistics, University of Pittsburgh.
